# Genetic characterization of the *apterous* Life Span Enhancer in *Drosophila melanogaster*

**DOI:** 10.64898/2025.12.04.692058

**Authors:** Cindy Reinger, Michèle Sickmann, Dimitri Bieli, Klemens E. Fröhlich, Alexander Schmidt, Markus Affolter, Martin Müller

## Abstract

The selector gene *apterous* (*ap*) of *Drosophila melanogaster* is best known for its prominent role in wing formation. *ap^null^* flies are viable but have no wings. However, *ap^null^* flies are associated with another phenotype whose analysis was largely neglected in the past 40 years: adults die precociously 2-3 days after eclosion. We have recently published a comprehensive analysis on the physiology of this striking phenotype. We showed that Ap protein is essential early in metamorphosis during hindgut remodeling where it is required for rectal papillae formation in the rectal ampulla. Wild-type flies have four rectal papillae that are involved in selective reabsorption of essential ions and metabolites from the primary urine produced by the Malpighian tubules. In *ap^null^* flies, rectal papillae formation is aborted prematurely. The corresponding papillar cells remain stuck in the rectal lumen. This leads to complete intestinal blockage and a cascade of pathologies resulting in premature death. In this study, we genetically identify and functionally dissect a novel tissue specific *cis-* regulatory element that directs Ap expression in the hindgut. This Life Span Enhancer (apLSE) maps to a ∼400 bp DNA fragment that is activated via only two small regulatory modules. Genetic analysis of *ap^LSE^* alleles proves that they are essential for papilla formation and thus for adult survival. Our studies solve the 110-year-old enigma about the *ap* syndrome. They establish the apLSE as a key player linking hindgut morphogenesis and survival. We propose that several aspects of the *ap* syndrome are a consequence of drastic changes in endocrine homeostasis.

## Introduction

The basic principles of eukaryotic genetics were elucidated by Morgan et al., 1915 (Morgan et al., 1915). Their work demonstrated that the hereditary rules postulated by Mendel based on his work with the garden pea *Pisum sativum* also apply to animals and that the chromosomes are the bearers of the hereditary material. The work of Morgan et al relied on the isolation of many spontaneous mutations and their careful genetic characterization. The vast majority of these mutations were homozygous viable and fertile and displayed an obvious recessive phenotype. An exceptional case was the allele *apterous^1^* (*ap^1^*). *ap^1^* homozygotes had no wings and halteres. However, females were also sterile and both sexes survived only for a few days (Metz, 1914). Since balancer chromosomes were not available yet and a homozygous *ap^1^* stock could not be maintained, the genetic analysis of the *ap^1^* allele was cumbersome. The genotype of heterozygous parents could only be deduced in retrospect from the presence of the recessive phenotypes in the progeny! Not surprisingly, the *ap^1^* allele is long defunct. But later isolates of *ap* alleles reproduced the phenotypes described by Metz (Butterworth & King, 1965). Decades later, cloning of the gene revealed that *ap* encodes a transcription factor of the LIM/homeodomain class (Bourgouin et al., 1992; Cohen et al., 1992). Thereafter, the role of *ap* during wing development was studied in great detail. There, Ap acts as a selector gene which specifies the identity of cells located in the presumptive dorsal wing compartment (reviewed Irvine & Rauskolb, 2001).

The second group of phenotypes described by Metz, female sterility due to a block of vitellogenesis and precocious adult death, were also investigated by several labs (King & Bodenstein, 1965; Butterworth & King, 1965; Postlethwait & Weiser, 1973; Postlethwait & Handler, 1978; Wilson, 1979, 1981a, 1981b, 1982; Stevens & Bryant, 1985, 1986). These studies unearthed further phenotypes associated with *ap* alleles. For example, the adipose tissue of strong *ap* alleles behaves abnormally (Butterworth & King, 1964a; Butterworth & King, 1964b; Butterworth, 1972). In wild-type flies, the adult fat body grows while the larval fat body degenerates. In strong *ap* alleles, the larval fat body shows no sign of degenerating, and the adult fat body maintains an immature appearance until death. Other, more obscure, phenotypes concerned aberrant sexual behavior in both sexes (Ringo et al., 1991, 1992). With one exception, these phenotypes were attributed to an endocrine lesion, namely a deficiency in juvenile hormone. In fact, measurements of juvenile hormone production in the corpus allatum gland indicated that it is reduced in *ap* alleles (Bownes, 1982; Altaratz et al., 1991). Furthermore, external application of juvenile hormone to the abdomen of *ap* females alleviated the block to vitellogenesis and larval fat body histolysis. These observations suggested that Ap protein might play a role in juvenile hormone synthesis in the corpus allatum. However, Ap expression in that tissue could not be detected, suggesting that the effect of *ap* on juvenile hormone synthesis is indirect (Cohen et al., 1992).

As alluded to above, one of the phenotypes belonging to the second group described by Metz was refractory to rescue by juvenile hormone treatment: the precocious adult death phenotype. In a recent study, we have described a comprehensive set of experiments that explain this fact (Reinger et al., 2025). Starting from the analysis of a hindgut specific enhancer, it could be demonstrated that early in metamorphosis, *ap* plays a crucial role during the formation of rectal papillae in the rectal ampulla. The *Drosophila* ampulla normally contains four papillae which are involved in reabsorption of molecules from the primary urin produced by the Malpighian tubules prior to excretion. In strong *ap* alleles, papilla cells remain clustered together leading to a complete blockage of the intestinal lumen. Consequently, freshly eclosed *ap* flies are unable to excrete their meconium. In addition, they barely ingest food. A succession of further pathologies reminiscent of ileus disease in humans ensues, leading to the untimely death of *ap* flies.

In this study, we have investigated and characterized the role of the responsible hindgut specific enhancer, the *apterous* Life Span Enhancer (apLSE). By careful genetic analysis of many *ap^LSE^* alleles, we show that the activity of this enhancer is under the control of two short regulatory modules. Analyses of *ap* gain-of-function conditions and the dominant *ap^Xa^* allele indicate that for normal papilla development, Ap expression must be restricted to two of the four papillae. Furthermore, we show that female fertility also depends on the presence of the apLSE. We propose that most *ap* phenotypes attributed to juvenile hormone dysfunction are a secondary consequence of a perturbed gastro-endocrine system due to complete hindgut blockage.

## Results

### 1. Mapping the Life Span Enhancer region within the intergenic spacer between *apterous* and *l(2)09851*

In the course of our genetic dissection of the *ap* regulatory region, numerous deletions with precisely defined end points have been generated (Bieli, et al., 2015a; Bieli et al., 2015b). Complementation crosses indicated that some of these deletions reproduced the precocious adult death phenotype previously described for strong *ap* alleles (Metz, 1914; Butterworth & King, 1965). In analogy to our approach to localize enhancers important for *ap* function during wing development (Bieli et al., 2015a, Bieli et al., 2015b), these preliminary observations suggested that it should be possible to map a regulatory region linked to this phenotype. Towards that end, the effect on life span of many deletions was carefully analyzed (for details, see materials and methods). The data for the most telling ones is presented in Fig. 1.

**Figure 1:**
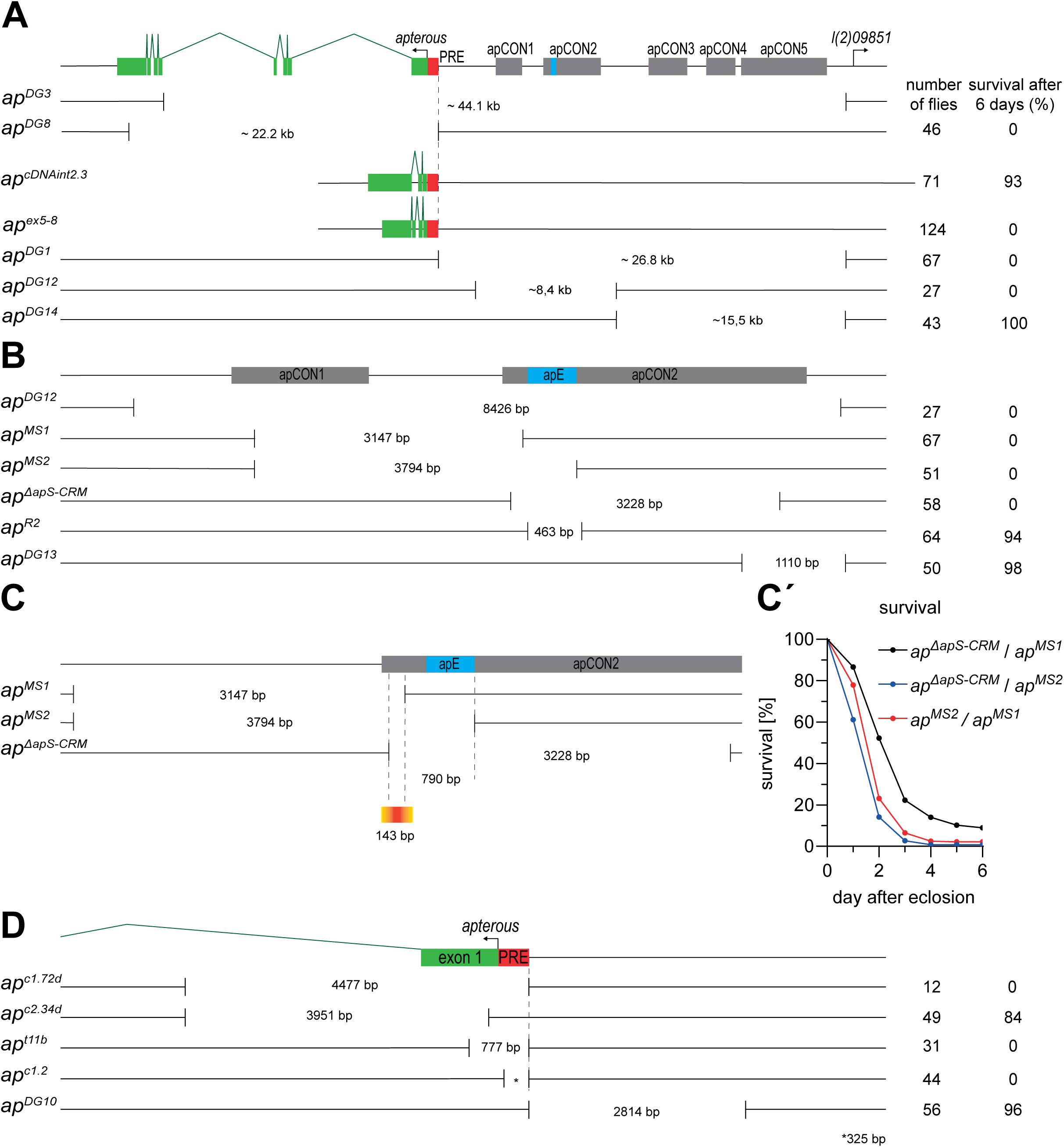
Functional identification of a putative *apterous* related Life Span Enhancer through targeted deletions and *in vivo* rescue assays. At the top, the *apterous* (*ap*) locus is depicted. The exons/intron structure is shown in green, the apPRE in red, the apE enhancer in blue and conserved stretches of regulatory DNA in grey according to Bieli et al., 2015b. On the right, the 5’ end of gene *l(2)09851* is indicated. Below, the position and size of various deletions are depicted. Survival of all deletions was tested over *ap^DG3^*. Survival data for each genotype is indicated in the two columns to the right. **A**: Mapping the Life Span Enhancer with larger deletions. As expected, lack of Ap protein in *ap^DG8^/ap^DG3^* flies leads to precocious adult death. Among the three deletions affecting the 27 kb intergenic spacer, *ap^DG12^* is the smallest deletion resulting in precocious death. Survival of allele *ap^cDNAint2.3^* indicates that the two large introns do not contain relevant regulatory input. Also, note that the 469 aa Ap protein expressed by *ap^cDNAint2.3^* is sufficient for survival and wing formation. In contrast, *ap^ex5-8^* produces only the 246 aa Ap protein and fails to rescue these two phenotypes. **B**: Zoom-in on the DNA interval defined by *ap^DG12^*. Deletion of the proximal apCON2 region in *ap^MS1^*, *ap^MS2^* and *ap^ΔapS-CRM^* is sufficient to elicit precocious adult death. This strongly suggests that the putative apLSE maps to this region. **C-Ć**: Complementation crosses between *ap^MS1^*, *ap^MS2^* and *ap^ΔapS-CRM^* result in precocious adult death in trans-heterozygous flies. Therefore, the 143 bp overlap between *ap^MS1^* and *ap^ΔapS-CRM^* defines the smallest DNA fragment containing the apLSE. However, survival of such flies is reproducibly slightly better than that of *ap^MS2^*/*ap^ΔapS-CRM^* flies. This suggests that more DNA to the right of the distal *ap^MS1^* break is required for full apLSE function. **D**: the apPRE also plays a role in survival. Several deletions mapping to the 5’ end of *ap* were tested *in trans* to *ap^DG3^*. The 325 bp interval (indicated by *) deleted by *ap^c1.2b^* contains the apPRE. *ap^c1.2b^* elicits precocious adult death. It therefore defines a second regulatory element required for survival. However, *ap^c1.2b^*/*ap^DG3^* flies also show a strong wing phenotype, indicating that the apPRE plays a role in more than one tissue. Alleles *ap^c1.72d^* and *ap^c2.34d^* share an identical proximal break point in intron 1 but differ in the position of the distal break. While *ap^c1.72d^* is a *bona fide ap^null^*, *ap^c2.34d^* is not. This deletion removes the coding part of the first exon including the ATG codon. Nevertheless, it produces a truncated version of the 469 aa as well as the 246 aa protein (see Supplementary Material). Although hemizygous *ap^c2.34d^* flies show a strong wing phenotype, they survive fairly well.

In a first set of experiments, larger *ap* deletions were assayed for their effect on survival (Fig. 1A). *ap^DG3^* removes the whole intergenic spacer as well as most of the open reading frame. *ap^DG8^* deletes all coding exons. Proteomics data indicate that *ap^DG3^* and *ap^DG8^* are *bona fide* null alleles (see Supplementary Material). *ap^cDNAint2.3^* lacks the two large introns 1 and 4. *ap^DG1^*, *ap^DG12^* and *ap^DG14^* are restricted to the intergenic spacer between *ap* and *l(2)09851*. *ap^DG1^* removes the whole 26.8 kb and therefore all five conserved regions apCON1 to apCON5 (Gohl et al., 2008; Bieli et al., 2015b). *ap^DG12^* lacks only apCON1 and apCON2, while *ap^DG14^* removes the distal three conserved regions.

The only genotypes which survive similarly well as *+/ap^DG3^* control flies are *ap^DG14^*/*ap^DG3^* and *ap^cDNAint2.3^*/*ap^DG3^*. The other three complementation crosses produce flies that die precociously before reaching day 6. Based on these observations, two conclusions can be drawn. Because no Ap protein is produced in *ap^DG8^* / *ap^DG3^* flies, this genotype strongly indicates that in wild-type flies, this protein is essential for survival. Among the deletions that do not affect the open reading frame, *ap^DG12^* is the smallest to elicit the precocious adult death phenotype. Hence, DNA missing in this allele must be essential for survival. Or with other words: a putative regulatory element should map to the 8426 bp interval containing apCON1 and apCON2.

This 8426 bp interval was further delimited with the help of *ap* rescue alleles that we generated for the analysis of wing specific enhancers (Bieli et al., 2015b). In brief, a set of alleles containing various combinations of the conserved regions apCON1 to apCON5 were crossed with *ap^DG3^/SM6a*. In the next generation, *ap^[rescue construct]^*/*ap^DG3^* flies were collected and their survival rate scored (Supp. Fig. 1). As expected for a negative control in which the whole 26.8 kb intergenic spacer is missing, *ap^attPΔEnh^/ap^DG3^* flies are all dead within a few days. In contrast, when all five conserved DNA intervals of the intergenic spacer are present, > 95% of *ap^C12345^/ap^DG3^* flies survive more than 6 days. This demonstrates that the genetic assay system is robust and useful to address our question. The remaining observations can be summarized as follows. Whenever apCON2 is lacking as for example in *ap^C1345^/ap^DG3^* flies, survival rate drops to 0%. On the other hand, all *ap^[rescue construct]^*/*ap^DG3^* variants that contain apCON2 survive. Furthermore, the same result was obtained with a 873 bp sub-fragment of apCON2 called apRXa (Bieli et al., 2015a). These results establish that within *ap^DG12^*, it is the proximal most part of apCON2 that confers the survival function to the *ap* gene. The associated putative tissue specific enhancer is referred to as *apterous* Life Span Enhancer (apLSE; Reinger et al., 2025).

### 2. A 409 bp interval in apCON2 contains the apLSE

Five smaller deletions that overlap with *ap^DG12^* were used to further delimit the location of the putative tissue specific enhancer. They are *ap^MS1^*, *ap^MS2^*, *ap^R2^*, *ap^ΔapS-CRM^* and *ap^DG13^* (Fig. 1B). As above, their effect on survival was tested in hemizygous flies. *ap^R2^*/*ap^DG3^* and *ap^DG13^*/*ap^DG3^* flies survive well. On the other hand, *ap^MS1^*/*ap^DG3^*, *ap^MS2^*/*ap^DG3^* and *ap^ΔapS-CRM^*/*ap^DG3^* flies die within the first six days. The overlap between the latter three deletions immediately pinpoints the location of apLSE: it maps to the proximal end of apCON2, right next to the apE wing enhancer. This conclusion was further corroborated by complementation crosses between *ap^MS1^*, *ap^MS2^* and *ap^ΔapS-CRM^* (Fig. 1C-C’). The three possible trans-heterozygous combinations produce flies with a short live span. The overlap between *ap^MS1^* and *ap^ΔapS-CRM^* is only 143 bp and defines the smallest interval for the position of the LSE. Also, *ap^MS1^*/*ap^ΔapS-CRM^* flies have normal wings, indicating that the function of the apE wing enhancer is not affected. However, we observed that life span reduction in *ap^MS2^*/ *ap^ΔapS-CRM^* flies was reproducibly slightly stronger, suggesting that *ap^MS1^* might not delete the entire apLSE.

### 3. The apPRE is also essential for survival

Apart from the apLSE, only one other region within the ∼50 kb *ap* locus was found that elicits the precocious adult death phenotype (Fig. 1D). Comparison of deletions *ap^c1.72d^* and *ap^c2.34d^* indicates that a 500 bp interval at the 5’ end of *ap* is essential for survival. In contrast, *ap^DG10^*, a deletion flanking that interval, confers good viability. *ap^t11b^* and *ap^c1.2b^* are two small deletions at the 5’ end which both show the precocious adult death phenotype. The smaller of the two, *ap^c1.2b^*, is only 325 bp in size and contains the apPRE (Schwartz et al., 2006; Oktaba et al., 2008). However, this deletion does not only affect survival but also wing and haltere development (Bieli et al, 2015b). With other words, it is not defining a tissue specific regulatory region but rather an element required for multiple *ap* functions.

To conclude, the data presented in the first three sections indicates that a tissue specific regulatory element called apLSE is contained near the proximal end of apCON2 region. It is strictly required for *ap* dependent survival.

### 4. Dissection of apLSE in the *ap^MS2^* landing site reveals two crucial modules for apLSE function

With the aim to further dissect the apLSE, we generated two deletions containing an attP landing site for ɸC31-mediated transgenesis as well as a FRT site for the removal of the transposition marker *mini-yellow* (for details, see materials and methods and Supp. Fig. 5). They are *ap^MS1^* and *ap^MS2^* (Fig. 1C). For the following analysis, *ap^MS2^* is the landing site of choice because it removes the complete apLSE. In a first set of experiments, 10 fragments spanning parts of the *ap^MS2^* deletion interval were inserted into the *ap^MS2^* landing site and their activity was assayed in hemizygous condition (Fig. 2A; Supp. Fig. 2A). The results were in sync with our previous observations. All relevant sequences for apLSE function are located at the proximal end of apCON2, between the proximal end of fragment 9 and the distal end of fragment 4. In a second set of experiments, four minimal LSE fragments were analyzed. Robust survival was obtained with minLSE (562 bp) and minLSE1 (409 bp), but not with minLSE2 (300bp) and minLSE3 (224 bp) (Fig. 2B; Supp. Fig. 2B). In order to locate important regulatory modules within the apLSE, 11 small internal deletions were induced in the context of the minLSE fragment and the corresponding alleles were generated (Supp. Table 3). Only two of these failed to rescue the precocious adult death phenotype in hemizygous flies: the 21 bp *ap^LSEΔ3.2^* and the 79 bp *ap^LSEΔ5^* deletions (Fig. 2C; Supp. Fig. 2C). Each of these intervals was subjected to another round of fine mapping. Within *ap^LSEΔ3.2^*, 11 bp defined by deletion *ap^LSEΔ3.2_C^* are essential (Fig. 2D; Supp. Fig. 2D; Supp. Table 3). Within *ap^LSEΔ5^*, 6 bp defined by deletion *ap^LSEΔ5.1^* are essential (Fig. 2E; Supp. Fig. 2E; Supp. Table 3). Finally, a minLSE variant containing deletions *ap^LSEΔ3.2_C^* and *ap^LSEΔ5.1^* was also generated (Fig. 2F; Supp. Fig. 2F). As expected, it completely inactivated the apLSE in hemizygous flies.

**Figure 2:**
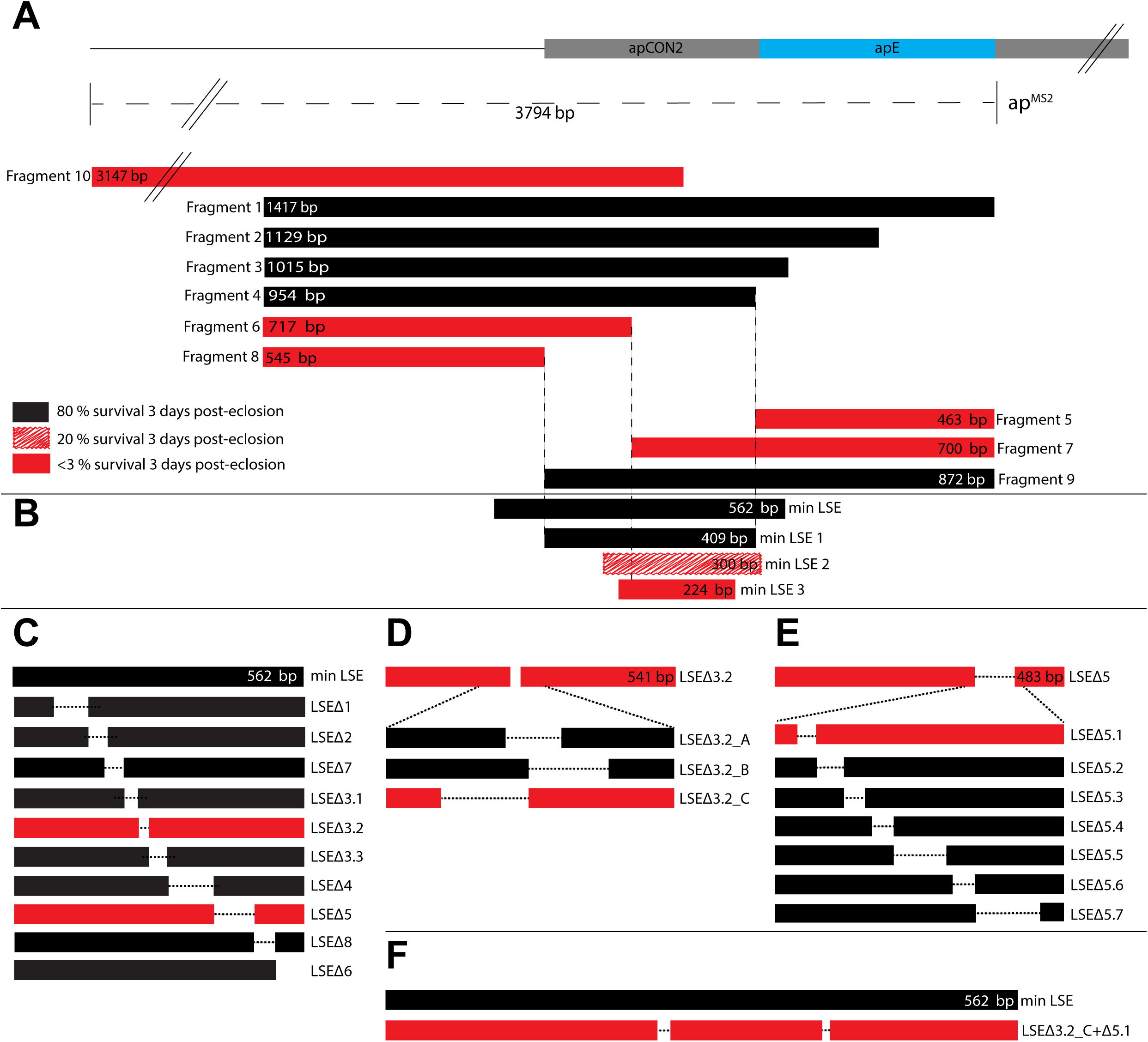
Within the apLSE, regions LSE-3.2_C and LSE-5.1 are required for survival. At the top, the proximal end of apCON2 is indicated in grey and the position of the apE enhancer in blue. Below, the deletion interval defined by *ap^MS2^* is shown. The dashed line represents the missing DNA. *ap^MS2^* serves as the host chromosome for the analysis of putative apLSE DNA fragments. *ap^MS2^* contains an attP landing site as well as an FRT which are required for ФC31-integrase mediated insertion of transgenic constructs and removal of the *mini-yellow* marker, respectively (for details, see Supp. Fig. 5C-E). All alleles were tested in hemizygous condition over *ap^DG3^*. Tested DNA fragments are indicated either in red or black. Red indicates “no rescue of precocious adult death”, black indicates “rescue of precocious adult death” (also applies to Fig. 4; the corresponding survival curves are depicted in Supp. Fig. 2A). Fragment sizes are shown within the colored bars. Exact end points and sizes of the deletions in A-F can be found in the Supp. Tables 1 and 3. **A**: 10 fragments spanning different parts of the DNA deleted in *ap^MS2^* were tested for their *in vivo* rescue activity. The deletion analysis indicates that the region between the proximal end of Fragment 9 and the distal end of Fragment 4 contains all necessary information for a functional life span enhancer (LSE). **B**: Four minimal LSE fragments (minLSE and minLSE1 to 3) of different length were tested. While minLSE and minLSE1 yielded robust rescue activity (see also Supp. Fig. 2B), it dropped considerably in minLSE2 and could no longer be detected in minLSE3. Based on these observations, minLSE was chosen for further dissection of apLSE function. **C**: Key DNA fragments within minLSE were identified with another set of 10 smaller deletions. Two of them, *ap^LSEΔ3.2^* and *ap^LSEΔ5^*, fail to promote adult survival in hemizygous flies. With *ap^LSEΔ4^*, activity drops to about 60% after 18 days, suggesting that the spacing between LSEΔ3.2 and LSEΔ5 may play a minor role (see Supp. Fig. 2C). **D**: The LSE-3.2 region was further delimited with three deletions. Among these, only *ap^LSEΔ3.2_C^* loses rescue activity comparably to *ap^LSEΔ3.2^* (see Supp. Fig. 2D). **E**: The LSE-5 region was further delimited with seven deletions. Among these, only *ap^LSEΔ5.1^* loses rescue activity comparably to *ap^LSEΔ5^* (see Supp. Fig. 2E). **F**: The double mutant *ap^LSEΔ3.2_C+Δ5.1^* behaves essentially like an *ap^null^* (see Supp. Fig. 2F).

These observations demonstrate that apLSE function is established by two short regulatory modules of 11 and 6 bp which are 136 bp apart. Although a drop in rescue activity was noted for *ap^LSEΔ4^* (Supp. Fig. 2C), the reduced spacing between the two (51 bp) does not seem to be crucial.

### 5. Phenotypes associated with *apterous* alleles

For the interpretation of phenotypes, knowledge of the molecular lesion and/or the amount of protein produced by a particular allele is helpful. Therefore, we have sequenced some of the alleles presented in this study. In addition, some alleles were subjected to proteomic analysis. The most relevant data is described below. For a detailed description of the proteomics data, see Supplementary Materials.

According to FlyBase, the *ap* locus produces 3 peptides: 469 aa (ap-PA), 468 aa (ap-PC) and 246 (ap-PB). Proteomics data indicates that only two of them are formed: ap-PA and ap-PB. Two *in situ* rescue alleles were generated that produce either ap-PA (*ap^cDNAint2.3^*; Fig. 1A; Bieli et al., 2015b) or ap-PB only (*ap^ex5-8^*; Fig. 1A). Proteomics data and whole mount embryo stainings with α-*ap* antibody demonstrate that both alleles express the expected protein variant (see Supplementary Material). *ap^cDNAint2.3^* can be maintained as a homozygous stock and is sufficient to rescue all phenotypes analyzed in this study. In contrast, *ap^ex5-8^* behaves like a null allele. Homozygous flies have no wings and die precociously (Fig. 1A; Table 1). *ap^c2.34d^* deletes a 3.95 kb fragment including part of the 5’-UTR and the first 13 aa encoded by exon 1 of the 469 aa peptide (Fig. 1D). Homo-as well as hemizygous flies survive well but produce only little wing tissue (Table 1; Fig. 3B-C). Proteomics data indicate that *ap^c2.34d^* produces normal amounts of the 246 aa peptide but also reduced quantities of a truncated version of the 469 aa peptide with undefined N-terminus. In contrast, *ap^c1.72d^* is a null allele which produces neither of the two peptides (see Supplementary data). It shares the same proximal break with *ap^c2.34d^* but the deletion also removes the transcription start site required for the produce ap-PA and the apPRE.

**Figure 3:**
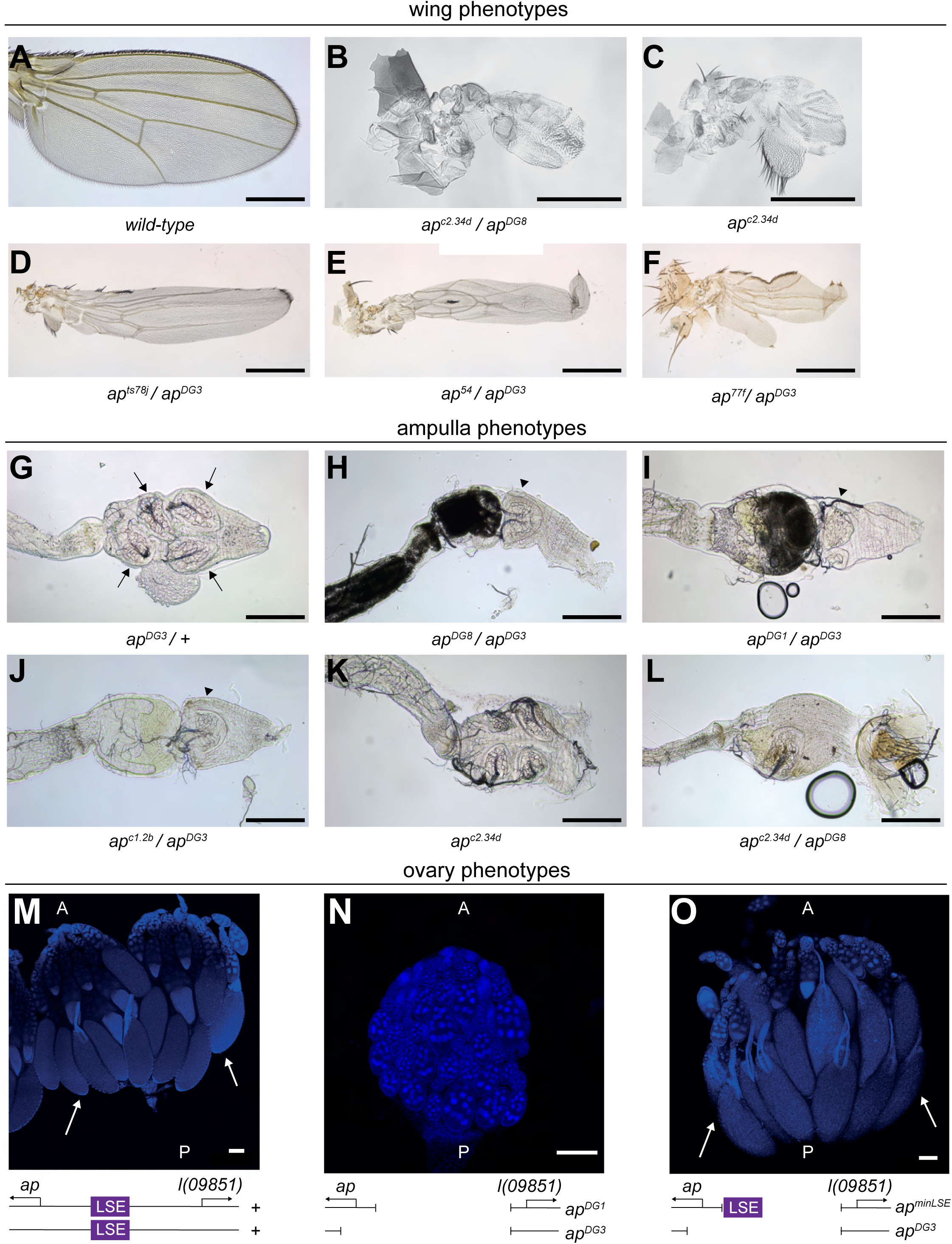
Overview of wing, ampulla and ovary phenotypes of different *ap* alleles. **A-F**: wing phenotypes of different *ap* genotypes imaged are shown. **A**: wild-type wing. Scale bar: 50 μm. **B-C**: *ap^c2.34d^* flies show strong wing phenotypes in (**B**) hemizygous and (**C**) homozygous condition. In the latter, only the most proximal structures are formed, the former phenotype is even stronger (nubbin). **D-F**: *ap^ts78j^/ap^DG3^* and *ap^54^/ap^DG3^* grown at permissive temperature and *ap^77f^/ap^DG3^* flies produce strap-like wings. **G-L**: Ampullae of different *ap* genotypes are shown with anterior to the left. Scale bar: 100μm. A normal *Drosophila* ampulla contains 4 papillae that are targeted by intricate tracheal networks (Wessing and Eichelberg, 1973). **G**: heterozygosity for *ap* (*ap^DG3^/+*) is sufficient for normal ampulla development. The four papillae are indicated by arrows. **H-J**: Absence of Ap protein (**H**), apLSE (**I**) or apPRE (**J**) lead to complete intestinal blockage and formation of a Reinger’s knot (arrow head). The dark material in the ampulla (**H,I**) is meconium whose excretion is prevented by the Reinger’s knot. **K**: in contrast to wing development, the ampulla of homozygous *ap^c2.34d^* flies is normal. It contains 4 papillae. **L**: hemizygous *ap^c2.34d^* flies fail to develop normal ampullae. Ampulla with a single papilla is shown. **M-O**: Ovary phenotypes in the presence (**M,O**) or absence (**N**) of apLSE. Anterior (A) is up, posterior is down (P). Below the ovary pictures, diagrams of the respective genotypes are shown. The box in magenta represents the apLSE. Arrows in **M** and **O** point to vitellogenic eggs. These are missing in **N** because vitellogenesis is not initiated. Note that like for ampulla development, one copy of the apLSE is sufficient to restore egg maturation and therefore fertility. Scale bar: 100 μm.

**Table 1.**
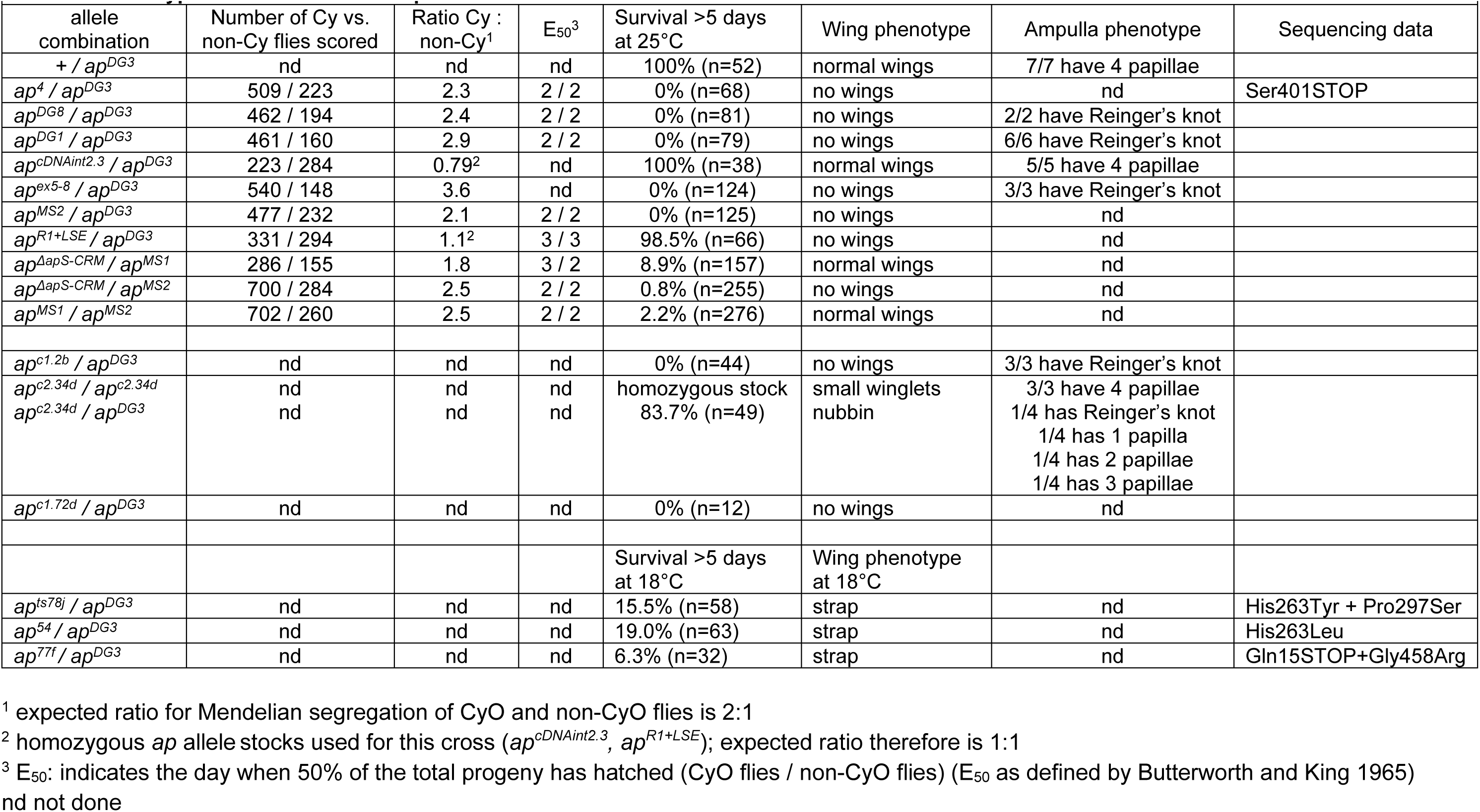
Phenotypic characterization of apterous alleles.

These results indicate that it is the 469 aa peptide that is essential to exert *ap* function, at least during wing and hindgut development. The phenotypes produced by *ap^c2.34d^* suggest that wing development is more sensitive to reduced protein levels than adult survival. The role of the 246 aa peptide remains elusive. It might play a role in other tissues where Ap is expressed. The comparison between alleles *ap^c2.34d^* and *ap^c1.72d^* suggests that deletion of the gene’s 5’ end is also affecting the production of the 246 aa peptide, although it is produced from alternate transcription start sites that are not affected by *ap^c1.72d^*.

#### A. Survival and developmental delay

In the *Drosophila* field, it is well established that hypomorphic allele combinations of essential genes can yield occasional adult escapers. This name derives from the fact that such flies are scored at a ratio clearly below that expected for Mendelian inheritance. This implies that most of their siblings die before eclosion. It is important to note that this is not the case even for *ap^null^* flies. With other words, in a cross between *ap*/+ parents, homozygous *ap* flies are expected to make up 25% percent of the total number of flies scored. Careful genetic analysis of the strong allele *ap^1^* yielded a value of 24.3% (Metz, 1914). Later, it was shown that many combinations of *ap* alleles emerge in numbers close to the expected Mendelian ratio. Also, while escapers tend to show a marked developmental delay, *ap* flies appear along with their heterozygous siblings (Butterworth & King, 1965). We have reevaluated these parameters for the classical allele *ap^4^* and some of the *ap* alleles generated in our lab (Table 1). Similar to previous studies, we find that all allelic combinations tested eclose in numbers close to the expected Mendelian ratio and that they are not developmentally delayed.

Freshly hatched *ap* flies are as agile as their heterozygous siblings. But after a few hours, their vitality decreases. Or as Metz (1914) wrote in his account on the *ap^1^* allele: “Instead of being vigorous and active they are weak and usually sluggish”. These observations indicate that *ap* flies initially are by no means handicapped by their genetic constitution. However, their lives take a fatal turn soon afterwards.

#### B. Wing phenotypes

*ap* is best known for its role in wing development. Null genotypes like *ap^DG8^/ap^DG3^* have no wings at all (Table 1). The classical *ap^4^* also belongs to this class. It produces a truncated protein due to a non-sense mutation in the homeodomain (Table 1). Depending on their strength, hypomorphic *ap* alleles can show a wide range of phenotypes. For example, when hemizygous animals of the temperature-sensitive alleles *ap^54^* and *ap^ts78j^* are grown under permissive conditions, they have strap-like wings (Fig. 3D-E). These are narrower than normal and form little or no wing margin. Molecularly, both contain a missense mutation that changes the identity of His263. This most likely destabilizes the second Zn-finger of the second LIM domain. In *ap^77f^*, a nonsense mutation truncates the protein at position Gln15 and is expected to produce a null allele. However, hemizygous flies also produce strap-like wings (Fig. 3F). Proteomic data shows that this allele produces both Ap peptides in similar relative quantities as *ap^c2.34d^* (see Supplementary Materials). This suggests that *ap^77f^* may also re-initiate translation and produce a truncated 469 aa peptide.

#### C. Precocious adult death and ampulla phenotypes

We have recently demonstrated that the hindgut of *ap^null^* and *ap^ΔLSE^* flies is malformed and blocked by a complete intestinal obstruction. Consequently, such flies do not excrete their meconium, do not properly initiate adult feeding and die within a few days after eclosion (Reinger et al., 2025). The hindgut phenotype is characterized by an abnormal rectal ampulla. In wild-type flies, it contains four rectal papillae (Fig. 3G). In *ap* mutants, they are not formed. Early in metamorphosis, the papilla precursor cells fail to reach their usual position in the ampulla. Instead, they remain clustered together and form an intestinal obstruction, which we refer to as Reinger’s knot (Reinger et al., 2025). This structure is readily detected in all flies of genotypes that fail to survive longer than 6 days (Fig. 3H-J; Table 1). It has previously been demonstrated that the presence of four papillae is not essential for survival (Schoenfelder et al., 2014). Moreover, we have generated papillae-less flies which survive well under standard laboratory conditions (Reinger et al., 2025). Ampullae with less than 4 papillae can be observed in hypomorphic *ap* alleles like for example *ap^c2.34d^*. Strikingly, while homozygous *ap^c2.34d^* flies have a strong wing phenotype (Fig. 3C), they develop normal ampullae (Fig. 3K). In contrast, hemizygous animals often contain less than four papillae (Fig. 3L) and their wing phenotype is further aggravated (Fig. 3B).

These observations suggest that the molecular requirements for wing and ampulla development react differentially to lower Ap protein concentrations.

#### D. Female sterility

Previous studies have reported that the precocious adult death phenotype observed in *ap* flies is always linked to female, but not male, sterility (Metz, 1914; Butterworth & King, 1965; Stevens & Bryant, 1985; Wilson, 1981b). When wild-type females hatch, their ovaries are not mature yet. The last step of oogenesis, vitellogenesis, is only initiated after eclosion. It starts as soon as females begin to eat and as long as the flies’ diet is supplemented with sufficient amounts of protein (Sang & King, 1961). However, although kept on rich medium, inspection of *ap* ovaries revealed that they do not undergo vitellogenesis and that they remain tiny. It is known that the transition to vitellogenesis is controlled by a check point which decides between progression through normal development or apoptosis of an egg chamber (Terashima & Bownes, 2004). Several lines of evidence suggest that this check point is under hormonal control. In insects including *Drosophila*, it has been known for a long time that juvenile hormone secreted by the corpus allatum is essential for vitellogenesis (Bodenstein, 1947; Vogt, 1948). Topical application of the Juvenile hormone analogue ZR-515 to the bellies of newly eclosed *ap* females triggered vitellogenesis and the formation of mature eggs (Postlethwait & Weiser, 1973). But importantly, Juvenile hormone treatment did not rescue their adult precocious death phenotype (Postlethwait & Weiser, 1973; Wilson, 1982).

Like observed for *ap^nulll^*, *ap^ΔLSE^* flies do not undergo vitellogenesis. In contrast, hemizygous flies of rescue allele *ap^minLSE^* have normal ovaries and they are fertile (Fig.3 M-O). It can be concluded that the presence of the LSE is not only required to rescue adult survival but also female fertility. Thus, in the context of the juvenile hormone data described above, it can be inferred that female sterility is only a secondary consequence of early adult lethality induced by the Reinger’s knot.

### 6. Regulation of the LSE

A functional LSE relies on the regulatory input mediated via two regulatory modules defined by alleles *ap^ΔLSE3_C^* and *ap^ΔLSE5.1^* (cf. Fig. 2). The latter is only 6 bp long and contains a sequence motive preferentially bound by Hox transcription factors. In support of this hypothesis, LSE function is also affected by mutagenesis of the characteristic ATTA core in an otherwise wild-type background (*ap^ΔLSE5.1_mut^*; Fig. 4C; Supp. Fig. 3C). No obvious transcription factor binding motive is contained within *ap^ΔLSE3_C^* deletion. Analysis of a library of 14 point mutations spanning the *ap^ΔLSE3_C^* interval did not detect bases crucial for binding of a transcription factor (Fig 4A; Supp. Fig. 3A). Note that we have used a similar approach to successfully characterize essential bases in the apE wing enhancer (Aguilar et al., 2023). The failure to detect relevant bases in *ap^ΔLSE3_C^* suggested that the organization of this regulatory module is more complex. *In silico* analysis indicated that *ap^ΔLSE3_C^* could disrupt a cluster of 4 binding site for the transcription factor Fork head. The data presented in Fig. 4A indicates that impairment of only one Fkh binding site at the time (as in mut_6 and _7 or _14, Fig. 4A) is not sufficient to abolish the function of the LSE3.2-C module. Therefore, the Fkh-cluster hypothesis was tested with four variants mutating an increasing number of Fkh binding site at a time (Fig. 4B; Supp. Fig. 3B). Hemizygous *ap^LSE-fkh_variation_1^* flies still survive well, even though 2/4 putative binding sites are abolished. But additional mutation of the most conserved Fkh binding site in allele *ap^LSE-fkh_variation_2^* is sufficient to elicit the precocious adult death phenotype in hemizygous flies.

**Figure 4:**
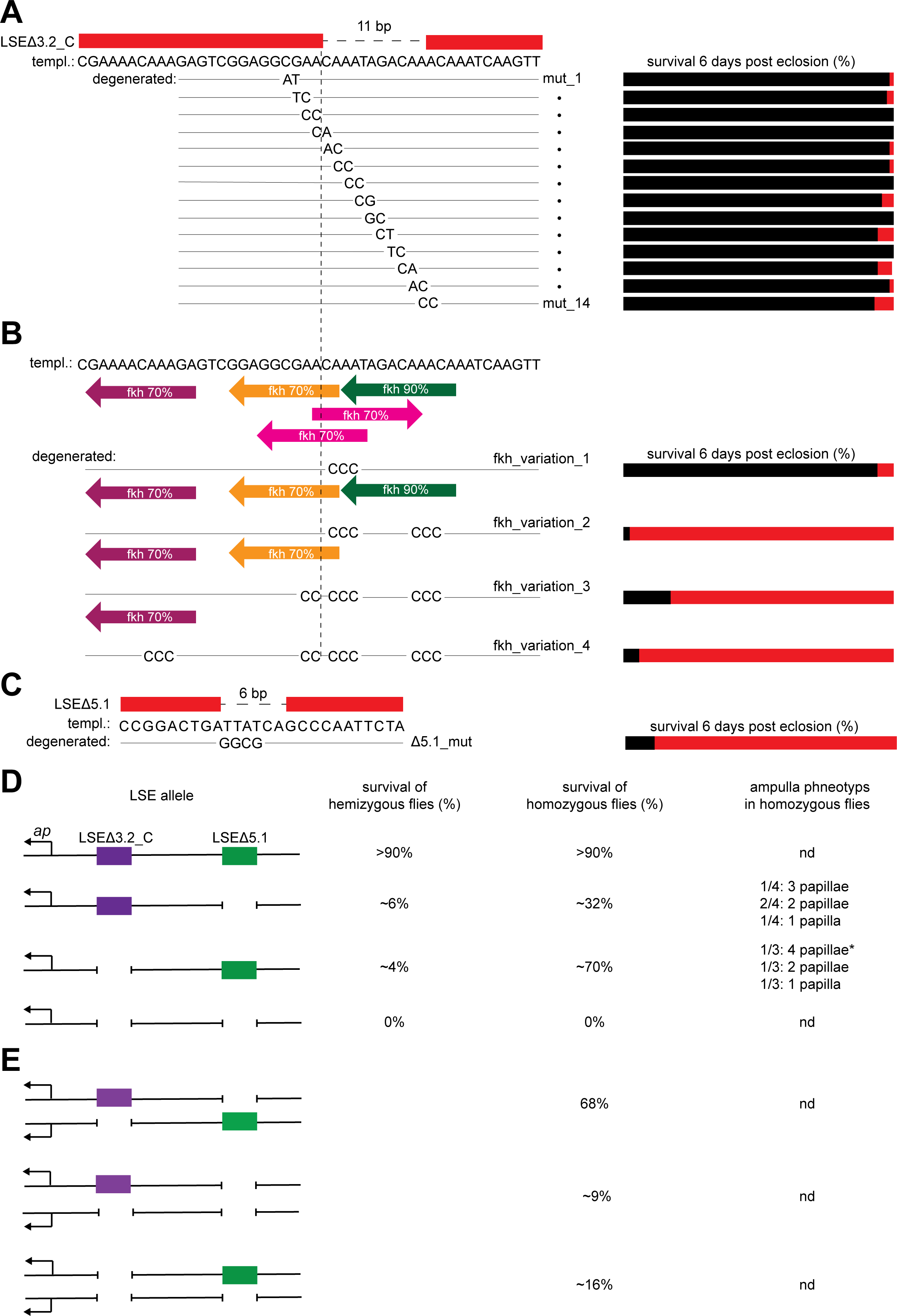
Functional dissection of alleles *ap^LSEΔ3.2^* and *ap^LSEΔ5.1^*. (**A-C**) Alleles were tested over *ap^DG3^*. Presence or absence of survival rescue activity is indicated by the same color code (black and red, respectively) as described in Fig. 2. Corresponding survival curves are shown in Supp. Fig. 3. **A**: Analysis of 14 2-bp point mutations in the *ap^LSEΔ3.2-C^* deletion interval. *Left:* At the top, the *ap^LSEΔ3.2-C^* deletion is shown. Below, 14 over-lapping 2-bp point mutations spanning the whole deletion are shown. Their names are *mut_1* to *mut_14*. *Right:* all 14 alleles efficiently rescue adult survival six days post-eclosion (see also Supp. Fig.3A). **B**: Analysis of potential Fork head (Fkh) protein binding sites in the *ap^LSEΔ3.2-C^* deletion interval. *Left:* at the top, a cluster of potential Fkh binding sites (indicated by colored arrows) predicted by JASPER is located within and around the LSE3.2 region. The percentages indicate the threshold used for *in silico* predicition. Four “*fkh_variation*” alleles were generated. In these, an increasing number of Fkh binding site was altered by changing the highly conserved “TTT” motive to “CCC.” *Right:* mutation of more than three of the five potential Fkh binding sites results in precocious adult lethality (see also Supp. Fig 3B). **C**: Analysis of a potential Hox binding site in the *ap^LSEΔ5.1^* deletion interval. *Left:* the *ap^LSEΔ5.1^* allele deletes a potential Hox binding site (ATTA motive). *Right:* mutation of this motif leads to precocious adult death in hemizygous *ap^LSE5.1_mut^* flies (see also Supp. Fig. 3C). As shown below, individual mutation of regulatory modules LSE3.2_C or LSE5.1 does not lead to a complete inactivation of the LSE. (**D-E**) Survival after 6 days of seven genotypes was scored and is indicated in % of flies collected on day 0. **D**: Four genotypes are shown. Their survival in homozygous or hemizygous (over *ap^DG3^*) condition is shown. apLSE activity is haplo-sufficient, as both homozygous (+/+) and hemizygous (+/*ap^DG3^*) flies survive normally. Individual deletion of regulatory modules LSE3.2_C or LSE5.1 generates hypomorphic *ap* alleles. In homozygotes, compared to wild-type, their rescue activity drops and is almost completely abolished in hemizygotes. Rectal ampullae of *ap^LSEΔ3.2-C^* and *ap^LSEΔ5.1^* homozygotes often contain less than 4 papillae (*: in this ampulla, two of the four papillae are smaller compared to the other two). Only simultaneous deletion of both regulatory modules phenocopies an amorphic *ap^null^* genotype where all flies die within a few days. **E**: three genotypes are shown. Survival is only mildly affected in trans-heterozygous *ap^LSEΔ3.2-C^*/*ap^LSEΔ5.1^* flies. Even a single copy of either LSE3.2_C or LSE5.1 is sufficient for a weak rescue effect. Note that in these genotypes, survival is improved compared to the hemizygous condition shown in Fig. 4D. Genotypes in 4E contain two functional ap transcription units, hemizygotes only one.

Genetic analysis of hemi- and homozygous combinations of *ap^ΔLSE3_C^* and *ap^ΔLSE5.1^* alleles suggest that the two regulatory modules are acting in an additive manner (Fig. 4D). Although most hemizygous *ap^ΔLSE3_C^*, *ap^ΔLSE5.1^* and *ap^ΔLSE3_C^*^+^*^ΔLSE5.1^* flies die precociously, the three genotypes behave differentially when analyzed in homozygous condition. *ap^ΔLSE3_C^*^+^*^ΔLSE5.1^* flies still fail to survive longer than a few days. In contrast, survival of *ap^ΔLSE3_C^* and *ap^ΔLSE5.1^* flies is clearly improved. Importantly, surviving flies are also fertile, suggesting that they are not affected by a Reinger’s knot. In fact, inspection of their hindguts reveals that ampullae with less than four papillae are formed (Fig. 4D). Trans-heterozygous combination of *ap^ΔLSE3_C^* and *ap^ΔLSE5.1^* leads to quite robust survival (Fig. 4E). And even a single copy of any of the two moduls can still mediate a mild rescue effect. These observations also provide an explanation why *ap^MS1^*/*ap^ΔapS-CRM^* flies survive somewhat better than *ap^MS2^*/*ap^ΔapS-CRM^* animals (cf. Fig. 1C’). In the former, one functional copy of the LSE5.1 module is left in place. The latter, like *ap^ΔLSE3_C^*^+^*^ΔLSE5.1^* flies, lack all four modules.

### 7. *apterous* expression in wild-type and *ap^ΔLSE^* alleles

We have previously demonstrated that apLSE activates Ap expression in the posterior hindgut (Reinger et al., 2025). A more comprehensive overview for timing and location of Ap expression in the embryonic hindgut was obtained by α-otp/α-ap double stainings. The homeodomain transcription factor Orthopedia (*otp*) is involved in hindgut development (Hildebrandt et al., 2020). *otp* mutant animals are embryonic lethal and their hindguts are only about one third of normal length. Otp expression in the hindgut is initiated in the proctodeum at stage 10 (Hildebrandt et al., 2020; Fig.5A). During germband retraction, the developing hindgut assumes the shape of a small tube and Otp is detected in the large intestine, the rectum and the anal pads (Hildebrandt et al., 2020; Fig. 5B-D’). Ap protein becomes visible at stage 13. It is restricted to the rectum and the anus in cells that also express Otp (Fig.5 C-D’). We note that *ap* transcript could already be detected in the presumptive anus of stage 10 embryos (Cohen et al 1992). Ap expression analysis in hemizygous embryos confirms that this hindgut pattern is strictly dependent on the presence of the *ap^LSE^* and the two regulatory modules *ap^LSE3.2^* and *ap^LSE5.1^* (Fig. 5E).

**Figure 5:**
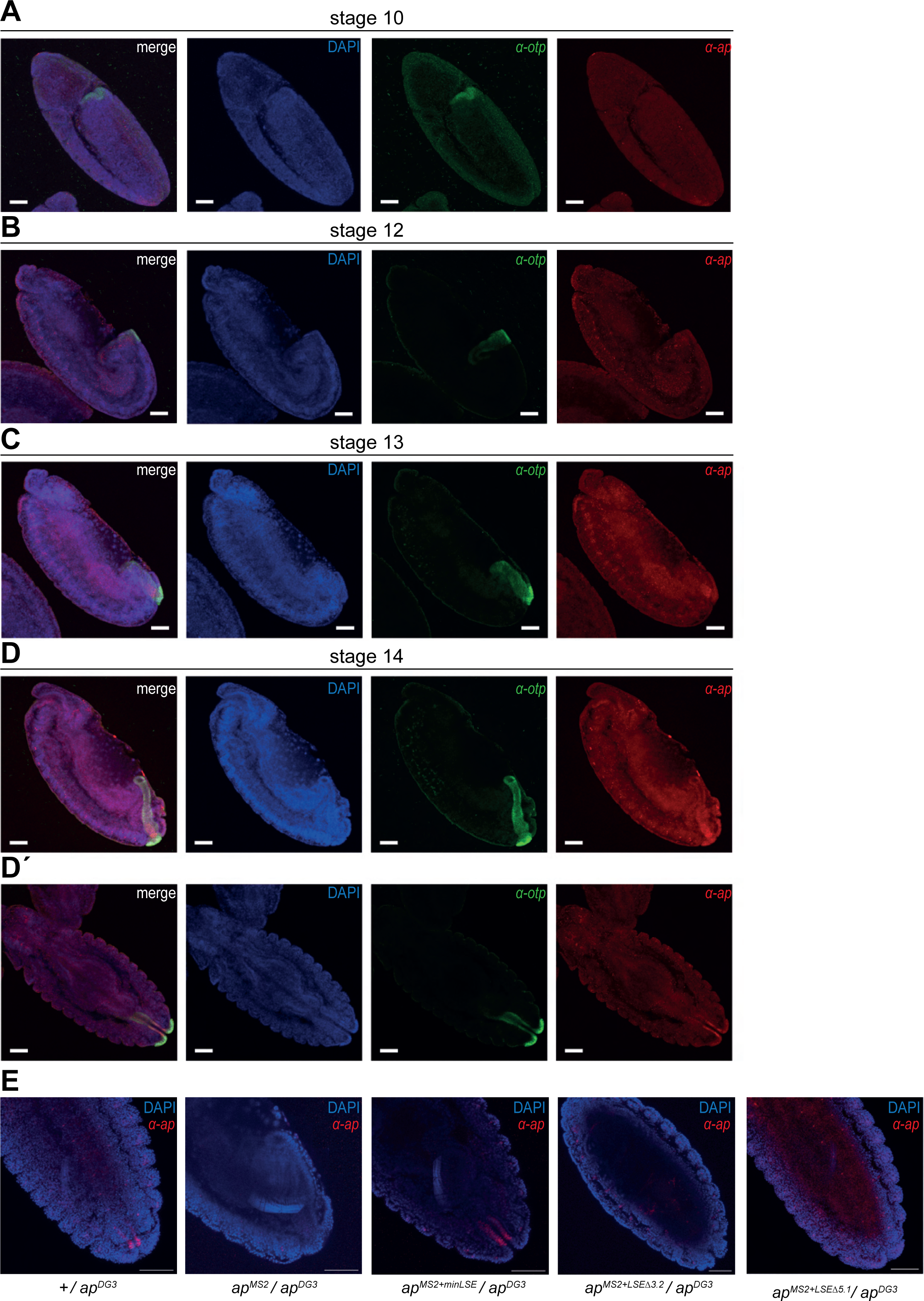
Ap expression in the embryonic hindgut. Ap expression in the embryonic hindgut was analyzed by double-immune-detection relative to the hindgut marker Orthopedia (Otp). Scale bar: 50µm. **A-D**: Lateral views of stage 10 to 14 embryos are shown (anterior in upper left corner). Otp is in green, Ap in red and DAPI in blue. Otp expression in the hindgut starts at embryonic stage 10 and remains active throughout the rest of embryonic development. Ap expression in the embryonic hindgut is first detected at stage 13. Compared to Otp, Ap expression remains restricted to the most posterior part of the hindgut, the rectum, where it persists also in older embryos. **D′**: Dorsal view. Otp and Ap seem to co-localize in the rectum and anal pads. **E**: Ap expression in hindguts of hemizygous *ap^LSE^* alleles. Dorsal views of stage 14 embryos are shown. Observed expression patterns correlate with survival: Ap is expressed in the presence of a functional apLSE (*+* and *ap^MS2+LSE^*) but not expressed in the presence of a non-functional apLSE (*ap^MS2^*, *ap^MS2+LSEΔ3.2^* or ap^MS2+LSEΔ5.1^_)._

During the course of this study, 7 *ap^Gal4^* drivers were generated (Supp. Fig. 4; Table 2). Some of them have previously been used to characterize the Ap hindgut expression pattern (Reinger et al., 2025). When crossed with a fluorescent reporter, our “wild-type” driver *ap^c1.4b-Gal4^* produces a pattern that is indistinguishable from the classical driver *ap^md544^* (Calleja et al., 1996). However, while *ap^md544^* is a strong *ap* allele, hemizygous *ap^c1.4b-Gal4^* flies show no phenotypes. But in the presence of a *UAS-ap* transgene, hemizygous *ap^md544^/ap^DG3^* are well viable and have normal wings (Table 2). This could not necessarily be expected because protein expression with the UAS/Gal4-system can lead to hypermorphic conditions. Our observations suggest that the genetic components that we are using do not lead to excessive Ap overproduction or that high Ap concentrations are tolerated as long as there is no ectopic expression (see next section). Therefore, these experiments were repeated with all other *ap^Gal4^* drivers (Table 2). In most cases, observed expression patterns and adult wing and hindgut phenotypes were as could be predicted based on observations described above. An exception is *ap^DG1-Gal4^* which causes a moderately strong GFP signal in fat body cells (Table 2).

**Table 2.** Rescue activity of various ap^Gal4^ drivers for wing development and survival.

| ap<sup>Gal4</sup> driver |  | UAS-CD8-GFP ap<sup>DG3</sup> | UAS-CD8-GFP ap<sup>DG3</sup> UAS-ap |
| --- | --- | --- | --- |
| ap<sup>md544</sup> | GFP pattern | expression in hindgut and wing disc; not in fat body | nd |
|  | wing phenotype | < nubbin | normal wings |
|  | survival >5 days | 0% (n=32) | 100% (n=38) |
| ap<sup>c1.4b-Gal4</sup> | GFP pattern | expression in hindgut and wing disc; not in fat body | nd |
|  | wing phenotype | normal wings | normal wings |
|  | survival >5 days | 98% (n=49) | 100% (n=25) |
| ap<sup>DG1-Gal4</sup> | GFP pattern | Moderate expression in fat body ; no expression in hindgut and wing disc | nd |
|  | wing phenotype | no wings | no wings |
|  | survival >5 days | 0% (n=40) | 0% (n=27) |
| ap<sup>MS3-Gal4</sup> | GFP pattern | expression in wing disc but not in hindgut and fat body | nd |
|  | wing phenotype | normal wings | normal wings |
|  | survival >5 days | 0% (n=44) | 0% (n=67) |
| ap<sup>MS5-Gal4</sup> | GFP pattern | no expression in wing disc, hindgut and fat body |  |
|  | wing phenotype | no wings | no wings |
|  | survival >5 days | 3.5% (n=57) | 0% (n=56) |
| ap<sup>R1-LSE-Gal4</sup> | GFP pattern | expression in hindgut but not in wing disc and fat body | nd |
|  | wing phenotype | no wings | no wings |
|  | survival >5 days | 100% (n=45) | 100% (n=66) |
| ap<sup>R1-LSEΔ3.2-Gal4</sup> | GFP pattern | no expression in hindgut, wing disc and fat body | nd |
|  | wing phenotype | no wings | no wings |
|  | survival >5 days | 0.8% (n=129) | 1.1% (n=180) |
| ap<sup>R1-LSEΔ5-Gal4</sup> | GFP pattern | no expression in hindgut, wing disc and fat body | nd |
|  | wing phenotype | no wings | no wings |
|  | survival >5 days | 0.8% (n=124) | 1.9% (n=103) |

For completeness, we would like to mention that we have also generated an *ap^c1.4b-LexA^* driver which faithfully reproduces the all aspects of the ap pattern (Supp. Fig. 4J; data not shown).

### 8. Ectopic *ap* expression in all 4 papillae phenocopies *ap^ΔLSE^* ampulla phenotypes

So far, we have demonstrated that complete loss of *ap* expression in two of the four papillae leads to Reinger’s knot formation and precocious adult death. Reduced *ap* expression in hypomorphic *ap* alleles often leads to a reduction in the number of papillae per ampulla, a condition which in general is viable. In a next step, we wished to address how ectopic Ap might influence ampulla development. It is well established that during wing development, ectopic Ap in the ventral compartment of the wing disc is not tolerated (Blair et al., 1994; Bieli et al., 2015a), leading to loss of wing tissue. In the embryonic hindgut, Ap is also not uniformly expressed but is restricted to two longitudinal stripes on opposite sides of the intestinal tube (Reinger et al., 2025).

As a means for ectopic Ap expression in all four papillae, the *byn^Gal4^* driver was employed. *UAS*-reporter gene expression can be detected from embryonic stages onwards in all cells of the developing hindgut (Lengyel & Iwaki, 2002). In adults, reporter gene expression is detected in all four papillae (Takashima et al., 2008). First experiments indicated that *byn^Gal4^*>*UAS*-*ap* animals grown at 25°C die before reaching the 3^rd^ larval instar stage (for details, see materials and methods). But survival to adulthood was obtained when the experiment was repeated with a *byn^Gal4^ tub-Gal80^ts^* recombinant (Table 3). Such flies were well viable and also fertile. These observations suggest that byn^Gal4^ driver activity is too powerful and that in the presence of *UAS*-*ap*, larval survival requires its modulation by Gal80^ts^. Inspection of byn^Gal4^ tub-*Gal80^ts^>UAS-ap* ampullae revealed that papilla formation is impaired. 3/3 ampullae contained only 1 or 2 papillae (Table 3; Fig. 6B). The experiment was repeated with an 18 → 29°C temperature shift during the mid-3^rd^ larval stage (for details see materials and methods). All emerging adults died within 5 days and 3/3 dissected hindguts contained a Reinger’s knot (Fig. 6C). Similar experiments were also carried out with *ap^EY03046^*, a *UAS*-*ap* analogue with reduced activity (Bieli et al., 2015a). *byn^Gal4^ tub-Gal80^ts^*>*ap^EY03046^* adults survived well and had normal ampullae (Fig. 6A). These observations indicate that restricted spatial ap expression in only two of the four papillae is essential. High levels (at 29°C) of ectopic ap phenocopy the *ap^ΔLSE^* ampulla phenotype while more moderate (at 25°C) ectopic expression interferes with normal papilla formation.

**Figure 6:**
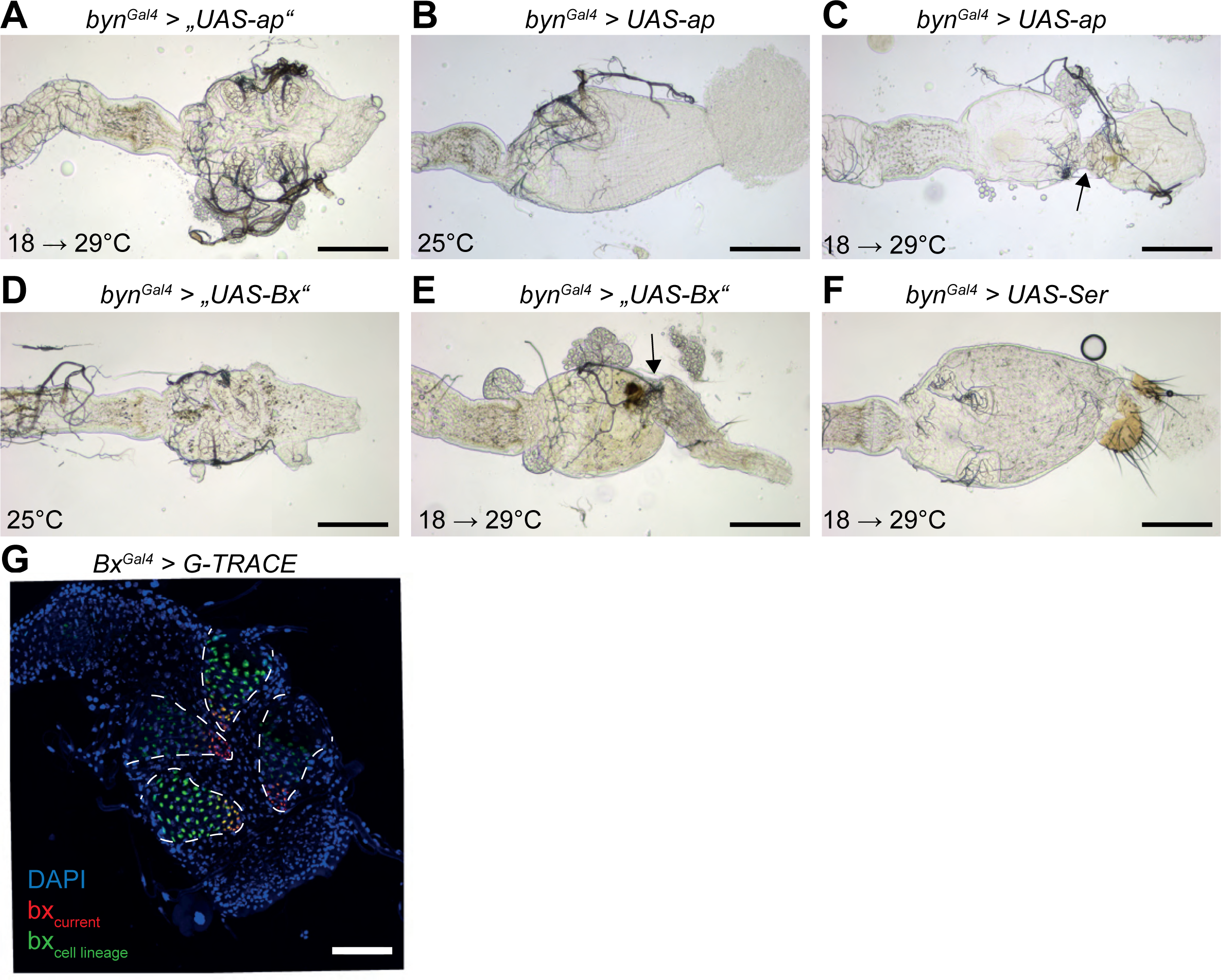
Ectopic expression of Apterous and downstream targets by *byn^Gal4^*. **A-F**: Dissected female hindguts are shown with anterior to the left. All genotypes also include *tub-Gal80^ts^*. Scale bar: 100µm. **A**: *byn^Gal4^*>*ap^EY03046^* (“*UAS-ap*”) grown with temperature shift regime. Four well developed papillae are formed. **B**: *byn^Gal4^>UAS-ap* gown at 25°C. In this ampulla, only a singe papilla is formed. **C**: *byn^Gal4^>UAS-ap* grown with temperature shift regime. A Reinger’s knot is formed (indicated by an arrow). **D**: *byn^Gal4^*>*P{GSV1}s-576* (“*UAS-Bx*”) grown at 25°C. In this ampulla, two papillae are formed. **E**: *byn^Gal4^*>*P{GSV1}s-576* (“*UAS-Bx*”) grown with temperature shift regime. Dissected ampullae invariantly display a Reinger’s knot (indicated by an arrow). **F**: *byn^Gal4^*>*UAS-Scr* grown with temperature shift regime. Ampullae containing less and smaller papillae are formed. **G**: Confocal image of a *Bx^Gal^*>*G-TRACE* ampulla is shown. The four papillae are encircled by a white dashed line. Note the differential expression of the green lineage signal.

**Table 3.**
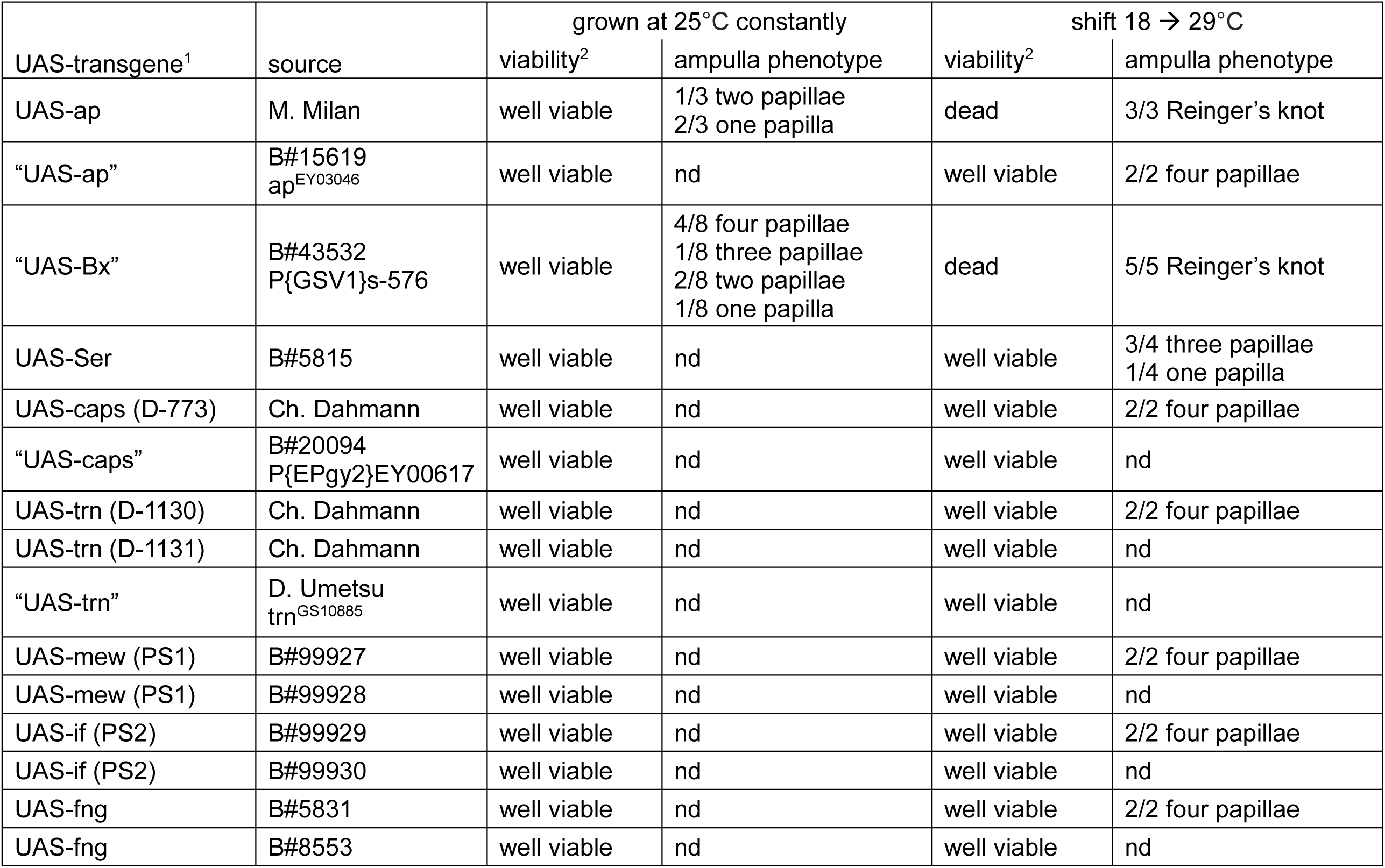
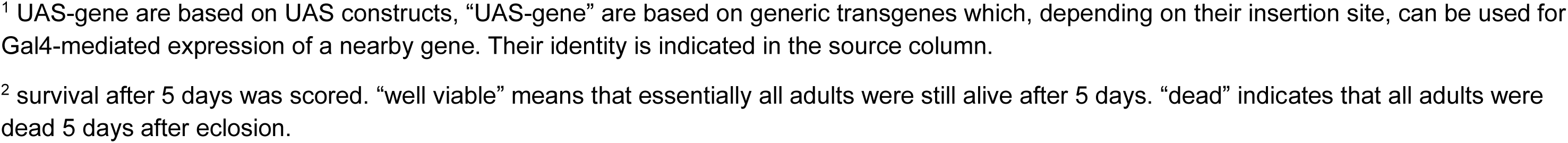
Ampulla phenotypes observed in tub-Gal80^ts^ byn^Gal4^>UAS-(ap or downstream-of-ap-target) flies.

| UAS-transgene<sup>1</sup> | source | grown at 25°C constantly |  | shift 18 → 29°C |  |
| --- | --- | --- | --- | --- | --- |
|  |  | viability<sup>2</sup> | ampulla phenotype | viability<sup>2</sup> | ampulla phenotype |
| UAS-ap | M. Milan | well viable | 1/3 two papillae<br>2/3 one papilla | dead | 3/3 Reinger's knot |
| "UAS-ap" | B#15619<br>ap<sup>EY03046</sup> | well viable | nd | well viable | 2/2 four papillae |
| "UAS-Bx" | B#43532<br>P{GSV1}s-576 | well viable | 4/8 four papillae<br>1/8 three papillae<br>2/8 two papillae<br>1/8 one papilla | dead | 5/5 Reinger's knot |
| UAS-Ser | B#5815 | well viable | nd | well viable | 3/4 three papillae<br>1/4 one papilla |
| UAS-caps (D-773) | Ch. Dahmann | well viable | nd | well viable | 2/2 four papillae |
| "UAS-caps" | B#20094<br>P{EPgy2}EY00617 | well viable | nd | well viable | nd |
| UAS-trn (D-1130) | Ch. Dahmann | well viable | nd | well viable | 2/2 four papillae |
| UAS-trn (D-1131) | Ch. Dahmann | well viable | nd | well viable | nd |
| "UAS-trn" | D. Umetsu<br>trn<sup>GS10885</sup> | well viable | nd | well viable | nd |
| UAS-mew (PS1) | B#99927 | well viable | nd | well viable | 2/2 four papillae |
| UAS-mew (PS1) | B#99928 | well viable | nd | well viable | nd |
| UAS-if (PS2) | B#99929 | well viable | nd | well viable | 2/2 four papillae |
| UAS-if (PS2) | B#99930 | well viable | nd | well viable | nd |
| UAS-fng | B#5831 | well viable | nd | well viable | 2/2 four papillae |
| UAS-fng | B#8553 | well viable | nd | well viable | nd |

<sup>1</sup> UAS-gene are based on UAS constructs, “UAS-gene” are based on generic transgenes which, depending on their insertion site, can be used for Gal4-mediated expression of a nearby gene. Their identity is indicated in the source column.
<sup>2</sup> survival after 5 days was scored. “well viable” means that essentially all adults were still alive after 5 days. “dead” indicates that all adults were dead 5 days after eclosion.

Based on these observations, it can be concluded that *ap* loss- and gain-of-function conditions produce similar phenotypes.

### 9. *Beadex*, a downstream target of *apterous*, may also play a role during hindgut formation

Genetic analyses of downstream targets of *ap* have identified several genes. Some of them are involved in cell-cell adhesion between dorsal and ventral wing epithelia (*trn*, *caps* (Milán et al., 2001), *integrin PS1* and integrin *PS2* (Blair et al., 1994)) while others are part of the signaling pathway establishing Wingless expression at the D-V compartment border (*Ser*, *fng* (Diaz-Benjumea & Cohen, 1993, 1995; Irvine & Wieschaus, 1994; Bachmann & Knust, 1998); *Bx* (Milán et al., 1998)). It appeared conceivable that some of these target genes might be reemployed during hindgut development. This possibility was tested by their ectopic expression in *byn^Gal4^ tub-Gal80^ts^*>*UAS*-(downstream target gene) animals (Table 3). We hypothesized that downstream targets normally activated by *ap* could elicit similar phenotypes as *UAS*-*ap* if they play a role in hindgut development. This expectation was fulfilled by “*UAS-Bx*”. *byn^Gal4^ tub-Gal80^ts^*>*P{GSV1}s-576* adults survived well when grown at 25°C but died within 5 days when shifted to 29°C during the 3^rd^ larval instar stage. Ampulla phenotypes closely emulated those observed with *UAS*-*ap*. At 25°C, 4/8 ampullae formed less than 4 papillae and at 29°C, 5/5 dissected hindguts showed the a Reinger’s knot (Table 3, Fig. 6 D-E). These results are only meaningful if the *Bx* gene is expressed during hindgut development. Signals detected in *Bx^Gal^*>*G-TRACE* ampullae indicate that this is the case (Fig. 6G). Intriguingly, the green lineage signal is clearly stronger in two papillae located on opposite sides of the ampulla. Assuming that this differential expression is meaningful for papilla formation, it is conceivable that ectopic Bx expression in *byn^Gal4^ tub-Gal80^ts^*>*P{GSV1}s-576* animals could interfere with this process. In wing development, it has been shown that Bx competes with Chip for Ap binding and that gain-of-function *Bx* mutants reduce Ap activity (Milán et al., 1998; Milán & Cohen, 1999; van Meyel et al., 1999; Rincón-Limas et al., 2000). Thus, Bx might perform a similar role during papilla formation.

Among the remaining target genes tested, only *UAS-Ser* produced a noticeable deviation from wild-type development. When shifted to 29°C during the 3^rd^ larval stage, adults survived well but the integrity of their ampullae was often impaired: 4/4 dissected hindguts showed less than 4 papillae which also appeared smaller than normal (Fig. 6F). Expression of Ser during papilla formation could not be monitored because *Ser^Gal4^*>*G-TRACE* animals died before eclosion. But a role for Ser during papilla formation would not be unexpected because Notch signaling is also involved (Fox et al., 2010).

### 10. The case of *ap^Xasta^*

Among the known *ap* alleles, *ap^Xa^* is special because it is dominant. *ap^Xa^/+* flies have an easily recognizable wing phenotype. The allele was induced by X-rays and was later characterized as a reciprocal translocation between chromosome arms 2R (where *ap* maps) and 3R (Serebrovsky & Dubinin, 1930; Hetherington et al., 1968). The break point in *ap* was molecularly characterized (Bieli et al., 2015a). It maps in the 27 kb intergenic spacer immediately distal of the *ap^E^* enhancer (Fig. 7A). The translocation fuses the *ap* locus to wing specific regulatory elements of the *Dad* gene located on 3R: Dadint52 and Dad4 (Weiss et al., 2010). Dad protein is expressed in a stripe along the A-P axis of the wing imaginal disc. Consequently, in *ap^Xa^*, *ap* falls under the *Dad* regulatory regime and Ap protein becomes ectopically expressed along the A-P axis also in the ventral compartment of the wing imaginal disc where it is normally absent (Bieli et al., 2015a).

**Figure 7:**
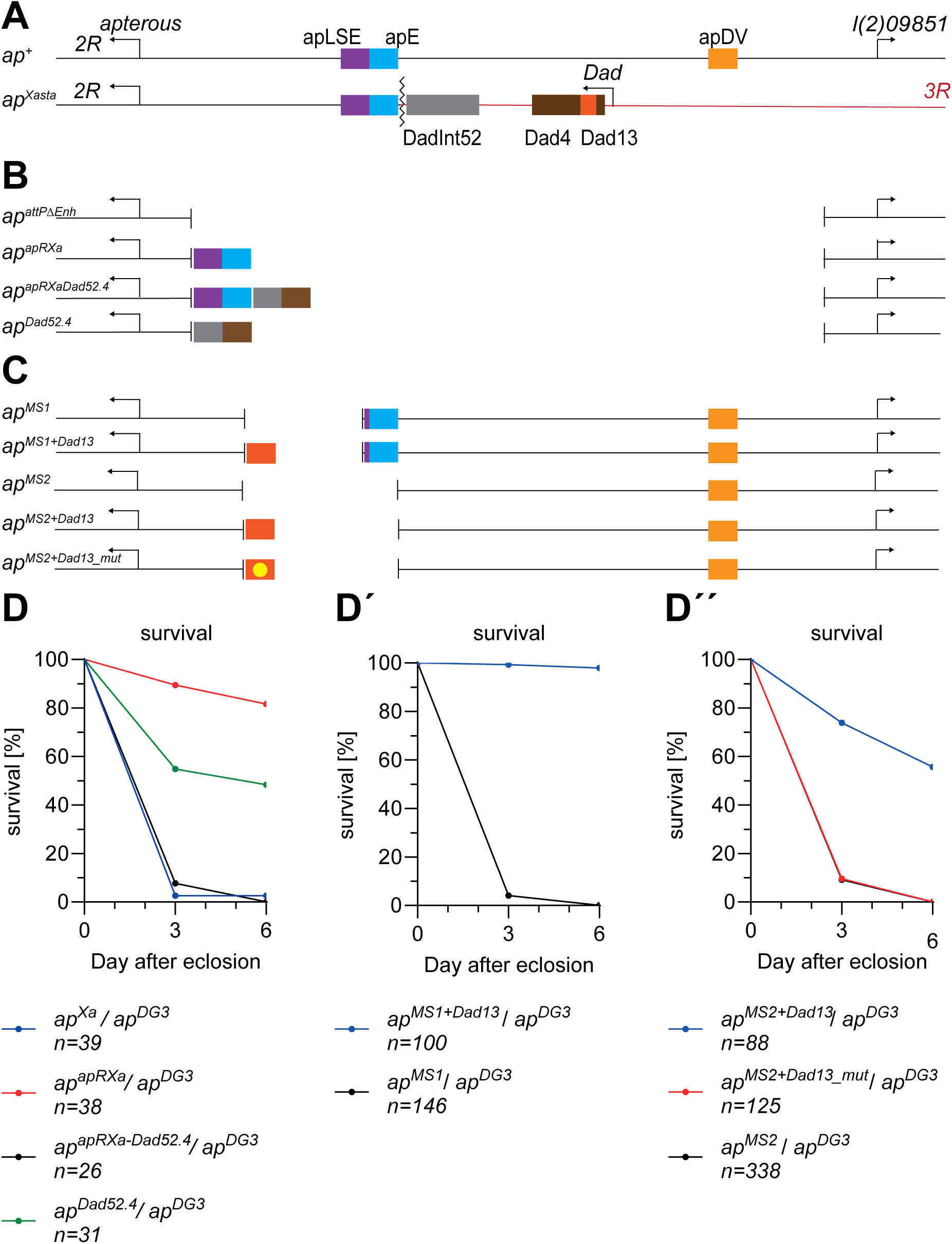
Maps and survival curves for *ap^Xa^* related alleles. **A-C**: Maps not shown to scale. **A**: Overview of *ap^+^* and *ap^Xa^* loci. At the top, the wild-type locus is shown along with the position of the hindgut specific apLSE (in magenta), the wing specific apE (in blue) and apDV (in light orange) enhancers. Below, the *ap^Xa^* locus is depicted. The wiggly line indicates the break point immediately distal to the apE enhancer which fuses the *ap* locus to the *Dad* gene normally located on 3R (Bieli et al., 2015a). The wing specific *Dad* enhancers DadInt52 (in grey), Dad4 (in brown) and Dad13 (in dark orange; a sub-fragment of Dad4) are located downstream of the *Dad* transcription start site (Weiss et al., 2010). **B**: Overview of three *ap^Xa^*-like alleles. In these, different combinations of apLSE, apE, DadInt52 and Dad4 enhancers were brought back into the *ap^attPΔEnh^* docking site. Note that in *ap^attPΔEnh^*, the whole intergenic spacer between *ap* and *l(2)09851* is missing. **C:** Overview of *ap^MS1^* and *ap^MS2^* alleles with or without Dad13 enhancer (indicated in dark orange). Note that a small part of the apLSE including module LSE5.1 (in magenta) and the apE wing enhancer (in blue) are still present in *ap^MS1^* derivatives and that apDV is present in all five alleles. In *ap^MS2+dad13_mut^*, 2/4 Mad/Medea binding sites normally present in Dad13 have been mutated (indicated by a yellow circle). **D**: Survival curves for *ap^Xa^* and alleles depicted in panel B are shown. Alleles that do not combine *ap* and *Dad* enhancers survive clearly better than those that do. **D’**: Survival curves for *ap^MS1^* derivatives are shown. **D’’**: Survival curves for *ap^MS2^* derivatives are shown. Hemizygous *ap^MS1+Dad13^* or *ap^MS2+Dad13^* flies survive clearly better than controls, indicating that Dad13 can at least partially substitute for apLSE. In contrast, rescue activity is completely abolished in hemizygous *ap^MS2+dad13_mut^* animals.

In the rearranged *ap^Xa^* locus, apLSE is still present between the *ap* promoter and the *ap^Xa^* break point (Fig. 7A). One could therefore expect that hemizygous *ap^Xa^* flies have a similar life expectancy as other genotypes where only one functional copy of *ap^LSE^* is present (e.g. *+/ap^DG3^* or *ap^D14^/ap^DG3^*; cf. Fig. 1A). This is, however, not the case. As previously demonstrated (Butterworth & King, 1965), ∼95% of hemizygous *ap^Xa^* flies die within 5 days. This experiment was repeated with very similar outcome with *ap^Xa^/ap^DG3^* flies (Fig. 7D). Inspection of their hindguts revealed that they often develop a Reinger’s knot. Sometimes, also ampullae with a single papilla are observed (Fig. 8B, B’; Table 4). Surprisingly, the integrity of the ampulla is also affected in *ap^Xa^/+* animals. They often have only 2, sometimes also 3 papillae (Fig. 8A; Table 4).

**Figure 8:**
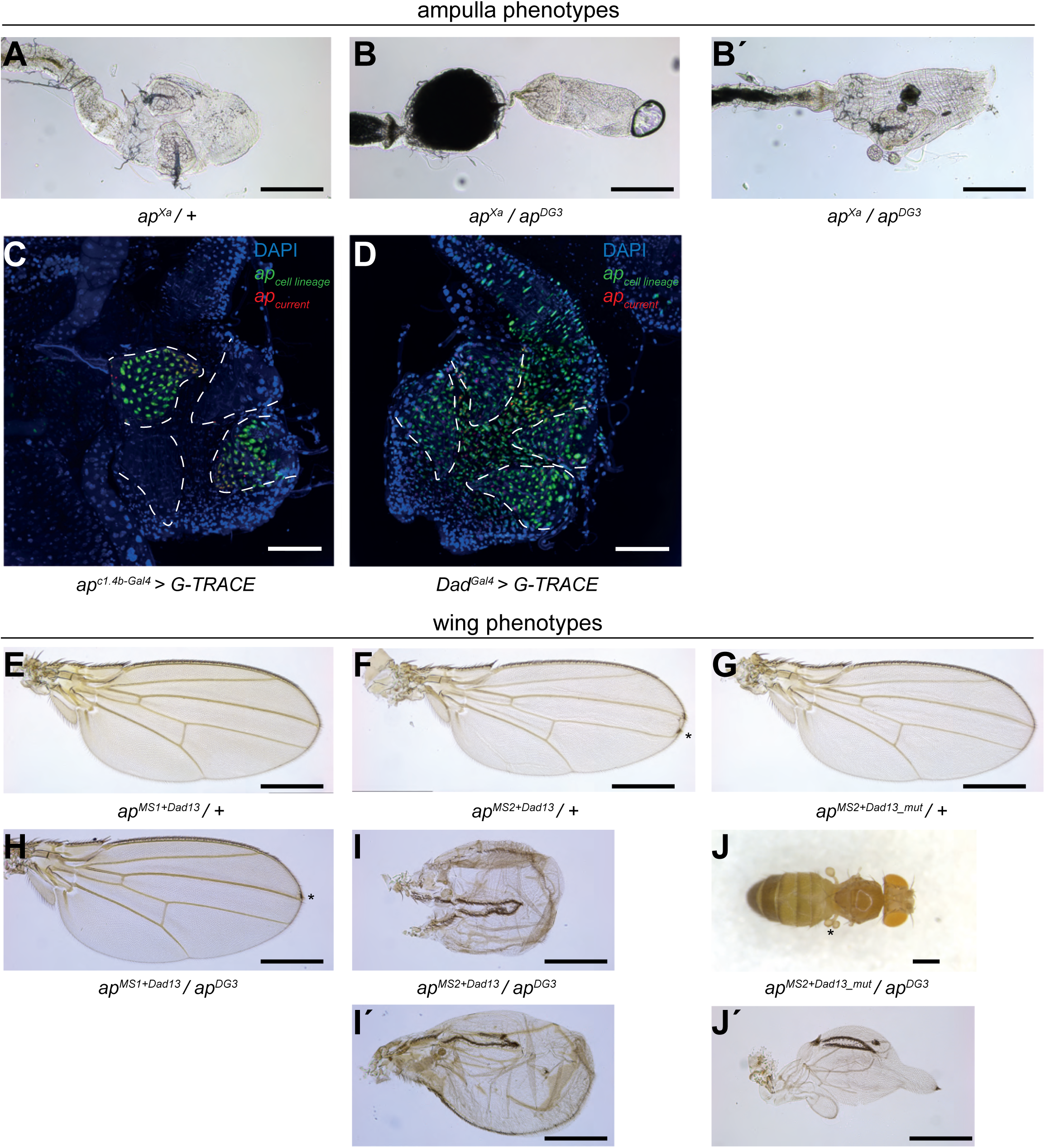
Ampulla and wing phenotypes of *ap^Xa^* and related alleles. **A**: Ampullae of *ap^Xa^/+* flies never contain 4 papillae. An example with only two papillae is shown. **B-B’**: ampullae dissected from *ap^Xa^/ap^DG3^* flies are shown. **B**: ampullae frequently develop a Reinger’s knot. **B’**: ampullae with 1 papilla can also be observed. **C-D:** G-TRACE signals detected in crosses with *ap^c1.4b-Gal4^* (C) and *Dad^Gal4^* (D) are shown. Papillae are outlined with dashed white line. **C**: as previously documented (Reinger et al., 2025), *ap^c1.4b-Gal4^* (C) leads to fluorescent signals in only 2/4 papillae. **D**: In contrast, *Dad^Gal4^* expression appears unrestricted and present in most cells of the ampulla including those forming the papillae. In both experiments, mainly the green lineage tracing signal is visible, suggesting that the Gal4 drivers are not very active in adults after eclosion. Scale bar: 50μm. **E-J:** Wing phenotypes of *ap^MS1^* and *ap^MS2^* derivatives containing Dad13 or Dad13_mut are shown. **E-F**: Most heterozygous *ap^MS1+Dad13^* and *ap^MS2-Dad13^* wings are normal but can show a weak margin defect with low frequency (indicated by *). **G**: the weak margin defect is never observed in heterozygous *ap^MS2-Dad13-mut^* animals. **H-J**: wing phenotypes of hemizygous flies (over *ap^DG3^*) are shown. **H**: the weak margin phenotype is sometimes observed in *ap^MS1+Dad13^* flies (indicated by *). **I-I’**: two examples of wings dissected from *ap^MS2-Dad13^* flies are shown. Their phenotype indicates that Dad13 partially substitutes for the missing apE enhancer. Wing margin formation suggests that sufficient Ap protein is produced to launch the auto-regulatory feedback loop via the apDV enhancer. Note that wing margin is absent in hemizygous *ap^apRXaDad52.4^* or *ap^Dad52.4^* flies which lack apDV (compare to Fig. 6B in Bieli et al., 2015a). **J**: In contrast, most *ap^MS2-Dad13-mut^*/*ap^DG3^* adults do not form wings but can show halteres with abnormal shape (indicated by *). **J’**: strap-like wings are formed by a fraction of such flies. Scale bar: 100μm, exept J: Scale bar: 0.5mm.

**Table 4.**
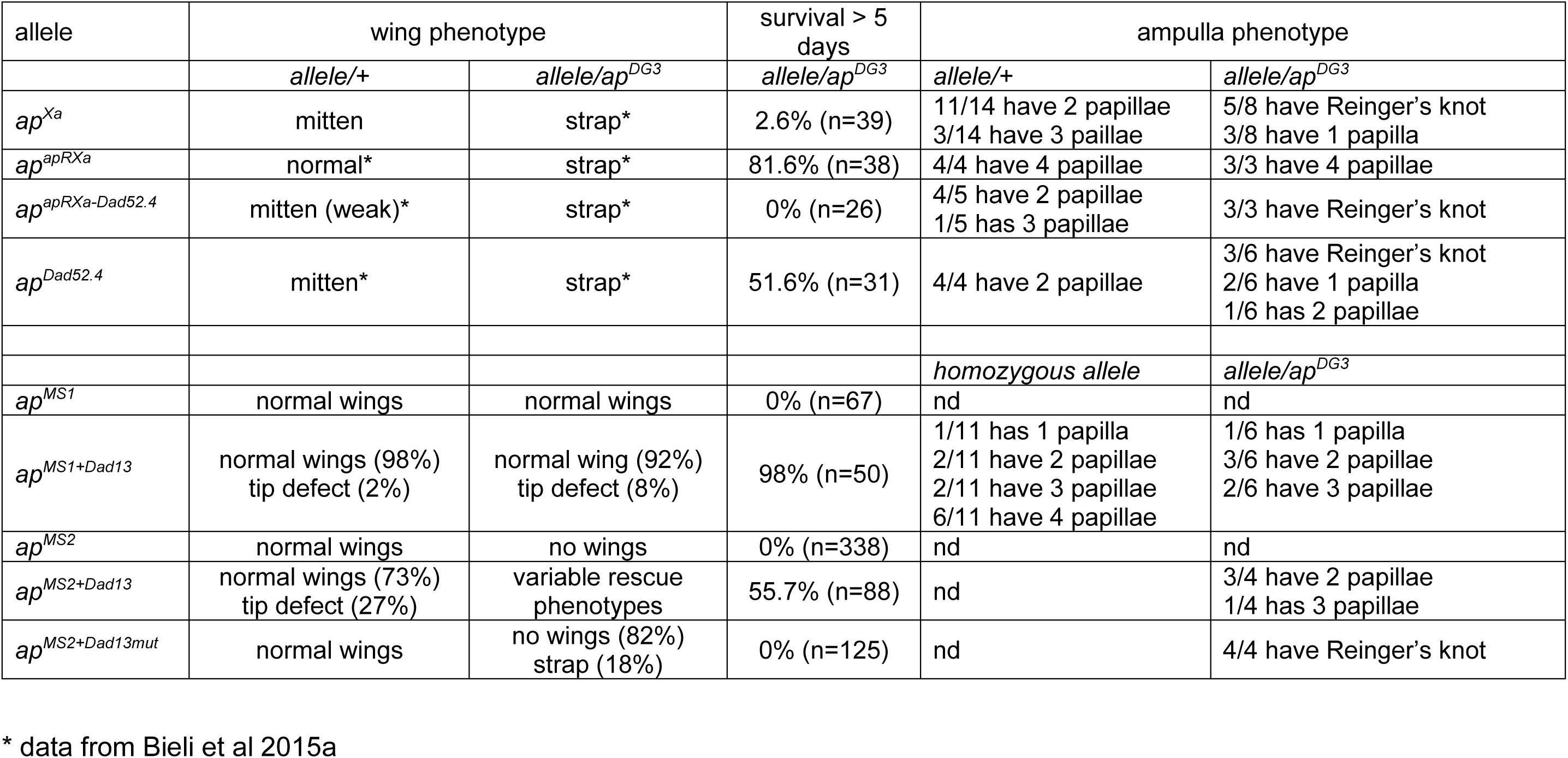
Phenotypes of *ap^Xa^*-related alleles.

During the characterization of *ap^Xa^*, three related *ap* alleles were generated. They are *ap^apRXa^*, *ap^apRXaDad52.4^* and *ap^Dad52.4^* (Bieli et al., 2015a). In these, only the wing specific regulatory components, which are juxtaposed by the translocation in *ap^Xa^* are present in place of the 27 kb intergenic spacer (Fig. 7B). The apRXa fragment (same as used in Supp. Fig. 1) contains the apLSE and apE. Dad52.4 is a 3148 bp fragment consisting of the two wing specific *dad* enhancers Dadint52 (941 bp) and Dad4 (2207 bp) (Fig. 7 A-B). The life span of these three *ap* alleles was assessed in hemizygous flies. ap*^apRXaDad52.4^* behaves very similar to *ap^Xa^*: they are short-lived and their hindgut contains a Reinger’s knot. Furthermore, the ampulla is also abnormal in heterozygous flies (Fig. 7D; Table 4). In contrast, whenever the *ap* or *Dad* components are present on their own, roughly 50% of the flies survive longer than 5 days (Fig. 7D). But inspection of their ampullae reveals that ampulla development is differentially affected. Heterozygous and hemizygous *ap^apRXa^* flies always have 4 papillae/ampulla. Although they survive significantly better than for example *ap^Xa^/ap^DG3^*, the ampulla phenotypes of heterozygous and hemizygous *ap^Dad52.4^* flies resemble those observed for *ap^Xa^* (Table 4).

These observations reveal a peculiar relationship between the *ap^LSE^* and the two *Dad* enhancers Dad4 and Dadint52. On the one hand, these dominantly interfere with papilla formation (as in heterozygous *ap^Xa^, ap^apRXaDad52.4^* and *ap^Dad52.4^* flies). On the other hand, the *Dad* enhancers can at least partially substitute for the LSE (as in hemizygous *ap^Dad52.4^* flies). These effects of the *Dad* enhancers on *ap* are analogous to those they exert on wing development (Table 4). There, they are brought about by ectopic Ap expression in the ventral wing compartment. This suggests that ectopic Ap could also be the reason for the observed ampulla phenotypes. This hypothesis was tested by monitoring the expression pattern in the rectal ampulla of *Dad^Gal4^>G-TRACE* flies. Compared to *ap^c1.4b-Gal4^*>G-TRACE, the *Dad^Gal4^*>G-TRACE signal is much less well defined but appears to be spread over all four papillae (compare Fig. 8C-D). This suggests that in *ap^Xa^* and derivatives, Ap could also be ectopically expressed at least during early metamorphosis. These observations reinforce our previous conclusion that Ap expression must be restricted to two of the four papillae.

### 11. LSE complementation by the Dad13 enhancer depends on *dpp* signaling

The effect of the Dad enhancers on Ap expression were analyzed in more detail with our *ap^MS1^* and *ap^MS2^* docking sites. They were used to introduce the 520 bp Dad13 enhancer, a sub-fragment of Dad4 (Fig. 7A), into the *ap* locus. The regulation of Dad13 has been analyzed in detail and shown to be a *bona fide* target of dpp-signaling (Weiss et al., 2010). The resulting new *ap* alleles *ap^MS1+Dad13^* and *ap^MS2+Dad13^* are depicted in Fig. 7C. Their effects on wing and ampulla development were scored in heterozygous, homozygous and hemizygous flies (Fig. 7, Fig. 8; Table 4). Heterozygous *ap^MS1+Dad13^* and *ap^MS2+Dad13^* flies show a weak wing phenotype with low penetrance (Fig. 8E-F). In hemizygous condition, the penetrance of this wing phenotype increases slightly in *ap^MS1+Dad13^* flies (Fig. 8H; Table 4). In contrast, *ap^MS2+Dad13^*/*ap^DG3^* flies show variable but strong wing phenotypes (Fig. 8I-I’’). The difference between the two genotypes can be attributed to the lack of apE in *ap^MS2^* derivatives. Thus, as described in the previous chapter for the two larger enhancers, Dad13 enhancer has a dominant effect on wing formation (as in *ap^MS1+Dad13^/+* and *ap^MS2+Dad13^/+* flies) but it can also at least partially substitute for the lack of the apE wing enhancer (as in *ap^MS2+Dad13^/ap^DG3^* flies).

Dad13 also functions as a substitute for apLSE in ampulla development. Hemizygous *ap^MS1+Dad13^* and *ap^MS2+Dad13^* flies survive much better than their hemizygous controls, *ap^MS1^* and *ap^MS2^*, respectively (Fig. 7D’-D’’). As previously noted, the small difference between *ap^MS1+Dad13^* and *ap^MS2+Dad13^* can probably be attributed to the presence of a single LSE5.1 module in the former (cf. Fig. 1C’). Dissection of the corresponding ampullae reveals that they are never normal but contain 1-3 papillae only (Table 4). This phenotype can be ameliorated in homozygous *ap^MS1+Dad13^* flies, in which 55% of the ampullae contain 4 papillae.

Since Dad13 is a target of *dpp*-signaling, we wanted to test whether *dpp*-signaling also plays a role for Dad13 function in the context of the *ap* locus. Electrophoretic mobility shift assays revealed that Dad13 contains four binding sites for the Brinker, Mad and Medea DNA binding peptides. Simultaneous mutation of binding sites 1 and 4 largely abolished Dad13 activity in co-transfection as well as in LacZ-reporter assays in transgenic embryos (Weiss et al., 2010). Therefore, the mutated Dad13m1m4 fragment of Weiss et al. was inserted into the *ap^MS2^* landing site. Then, wing and ampulla phenotypes of the new allele *ap^MS2+Dad13mut^* were analyzed. Hemizygous animals die precociously and all of their hindguts contain a Reinger’s knot (Fig. 7D’’; Table 4). The weak dominant wing phenotype is no longer seen (Fig. 8G) and 82% of hemizygous flies have no wings (Fig. 8J). The remaining 18% can have strap-like wings (Fig. 8J’). This weak residual activity during wing development might be explained by the presence of binding sites 2 and 3 identified in Dad13 by Weiss et al. These observations strongly suggest that the rescue activity of the Dad13 enhancer depends on *dpp*-signaling.

## Discussion

### 1. The *apterous* syndrome

*ap^null^* adults are characterized by two easy to spot phenotypes. They lack wings and halteres and fail to live longer than 2-3 days. The prominent role of *ap* in wing development is documented by contributions from many labs. The regulatory cascade leading to Ap expression in the dorsal wing epithelium is rather well understood (reviewed Irvine & Rauskolb, 2001). Ap expression in the wing imaginal disc is initiated via an enhancer known as apE (Bieli et al., 2015a, Bieli et al., 2015b; Aguilar et al 2023). Flies lacking this enhancer are phenotypically indistinguishable from true *ap^null^* flies: they look like ants because their wings (and halteres) are missing. However, while *ap^ΔE^* flies can be maintained as a homozygous stock, *ap^null^* flies perish soon after eclosion. This difference indicates that *ap* regulatory elements other than apE are required to warrant adult survival. Support for this hypothesis comes from two observations. First, the temporal requirement for Ap in wing development and adult survival fall into separate time intervals. While the wing function is essential during the second larval instar stage, the survival function is indispensable during the first two days of metamorphosis (Wilson, 1981b; Reinger et al., 2025). Second, mosaic analysis has indicated that Ap function essential for adult survival is required in the posterior abdomen (Wilson, 1981a). In this study, we describe the isolation and characterization of a tissue specific *ap* enhancer whose activity fits these two temporal and positional requirements. We have christened it Live Span Enhancer (apLSE). It maps right proximal to apE but does not overlap with it (Fig. 2). apLSE is active in the posterior hindgut (Fig. 5). Like *ap^null^* flies, *ap^ΔLSE^* flies are viable and show the precocious adult death phenotype. Lethality is induced by an innate blockage of the intestinal lumen in the posterior part of the rectal ampulla. The blockage, referred to as Reinger’s knot, is brought about by inappropriately positioned rectal papilla cells which fail to find their normal position in the ampulla. *ap^ΔLSE^* flies are unable to excrete their meconium, barely eat and frequently display a condition known as proboscis extension sleep. In addition, their hindguts are bloated and eventually become permeable and disintegrate. We have proposed that the character and succession of the various phenotypes indicate a condition highly reminiscent of ileus disease in humans (Reinger et al., 2025).

Prior to 1990, the strongest available *ap* allele of choice was *ap^4^*. It was demonstrated that in *ap^4^* flies, precocious adult death is always associated with two other phenotypes. They are female sterility and persistence of larval fat cells in the imago (King & Sang, 1958; Butterworth & King, 1964, 1965; Butterworth, 1972; Postlethwait & Weiser, 1973; Wilson, 1982; Richard et al., 1993; Aguila et al., 2007). Apart from the various phenotypes listed in the previous paragraph, female sterility and persistence of larval fat cells are also salient in *ap^ΔLSE^* flies. In humans, such a combination of multiple medical problems showing the existence of a particular disease is referred to as “syndrome”. With respect to the linked phenotypes in *ap^ΔLSE^* flies, it therefore seems appropriate to speak of the *ap* syndrome. Although we have not investigated larval fat cell histolysis in detail in *ap^ΔLSE^* flies, our observations strongly suggest that all phenotypes constituting the *ap* syndrome rely solely on the absence of the apLSE in *ap^ΔLSE^* flies. Thus, all phenotypic aspects of the *ap* syndrome are a consequence of Reinger’s knot formation early in metamorphosis. First signs of the looming highly stressful syndrome are the failure of meconium excretion and almost complete absence of food ingestion.

Several of the *ap* syndrome phenotypes might be elicited by inter-organ communication via the gastro-endocrine system. When wild-type flies are exposed to an environment where food resources are scarce, their physiology switches to a survival mode. For example, a nutrient dependent checkpoint in mid oogenesis is triggered before egg chambers enter vitellogenesis (Drummond-Barbosa & Spradling, 2001). Hence, yolk deposition into the maturing oocyte is blocked. In addition, programmed cell death of nascent oocytes is a well-characterized mechanism for re-allocation of energy stores (Pritchett et al., 2009). Similar to *ap^ΔLSE^* flies (Fig. 3P-R), ovaries of starved flies are clearly smaller and do not contain vitellogenic eggs. Considerable evidence indicates that an environmental stress response to starvation is mediated by ecdysone signaling (Hirashima et al., 2000; Ishimoto et al., 2009). A more recent report has shown how ecdysone titers are influencing oogenesis via the peptide hormone Ecdysis Triggering Hormone (Meiselman et al., 2018). Experiments with *ap^4^* flies have shown that vitellogenesis and larval fat cell histolysis can be rescued by the topical application of a juvenile hormone analogue on the female abdomen. But importantly, precocious adult death cannot be prevented by this treatment. These observations strongly suggest that the female sterility and delayed fat cell histolysis phenotypes observed in *ap^ΔLSE^* flies are induced by aberrant hormonal signaling as a secondary consequence of complete intestinal blockage.

### 2. Regulation of apLSE

The Ap protein can be detected in multiple tissues at different developmental time points. Tissue specific enhancers have been described for several of these organs, including muscle (Capovilla et al., 2001) and nervous system (Lundgren et al., 1995; Stratmann & Thor, 2017). Over the last decade, our lab has concentrated on deciphering the regulatory logic behind wing development. Three regulatory elements are essential to establish a normal wing. First, Ap expression is initiated by the apE enhancer targeted by proteins binding to well-defined Hox- and GATA-transcription factor binding sites. Then, ensuing Ap expression initiates an auto-regulatory feedback loop required to activate the apDV enhancer. A Polycomb Response Element (apPRE; Fig.1A) located immediately upstream of the *ap* transcription start site could transmit enhancer activity to the Pol II transcription machinery (Bieli et al., 2015b). The succession of these events starts during the second larval instar stage and coincides with the temperature sensitive period for wing development of an *ap^ts^* allele (Wilson, 1981b). In this study, the regulation of Ap expression in the posterior hindgut has been characterized to a similar level of resolution as for wing development. A single enhancer, the apLSE, is sufficient to launch Ap expression in combination with the apPRE during late embryogenesis. Like apE, apLSE is activated by two regulatory modules. One of them is most likely targeted by an undefined Hox-transcription factor. A possible candidate is the homeobox gene *otp* which is expressed in the hindgut and whose protein co-localizes with Ap (Fig. 5). The second module could be targeted by the Fork head transcription factor. *fkh* embryos fail to from a hindgut while it is clearly shorter in *otp* embryos. Unfortunately, these morphological shortcomings preclude an analysis of Ap expression in *fkh* and *otp* mutant embryos.

Curiously, and in contrast to wing development, the onset of Ap expression in the hindgut happens 4-5 days (at 25°C) before the protein becomes essential during early metamorphosis when the rectal papillae are formed. In fact, for successful papilla formation, it is sufficient to provide Ap activity (Wilson, 1981b) or initiate Ap expression in an ap^ΔLSE^ background (Reinger et al., 2025) late during the 3^rd^ larval instar stage. Hence, the question arises why Ap expression is already initiated during late embryogenesis and not only by the onset of metamorphosis. Patterning of the *Drosophila* hindgut relies on the localized expression of several signaling pathways as well as transcription factors (reviewed in Lengyel & Iwaki, 2002). Importantly, it occurs when the relevant tissue is still small and expression patterns of downstream targets can be more precisely defined.

As alluded to above, it is possible that apLSE is a target of the transcription factor Fork head. This is the founding member of a large family of proteins which share a characteristic winged helix DNA-binding domain (reviewed in Golson & Kaestner, 2016). Fork head has been classified as a pioneer factor that can bind and initiate opening of silent chromatin regions, recruit chromatin remodelers as well as epigenetic histone modification enzymes. As a result, a targeted enhancer is activated and remains so also after multiple cell divisions (reviewed in Larson et al., 2021; Barral & Zaret, 2024). Thus, it is possible that Ap expression is established early because the transcription factor cocktail required for apLSE activation is only available during late embryogenesis and that apLSE remains active because of the biochemical properties of the proteins involved. Our results show that the effects of the LSE3.2 and LSE5.1 regulatory modules are additive and that absence of Fkh binding sites (as for example in homozygous *ap^MS2+LSEΔ3.2-C^* animals) does not abolish apLSE function. In this case, the pioneer factor hypothesis implies that also the Hox transcription factor binding to the LSE5.1 module should have pioneer factor characteristics. To which degree the apPRE is part of this putative cellular memory mechanism remains to be shown.

## Acknowledgements

We would like to thank Gustavo Aguilar, Milena Bauer and Sophie Schnider for fruitful discussions and a helping hand whenever needed. The Imaging Core Facility of the Biozentrum Basel was invaluable for the whole LSE project. A big thank you to Johannes Stratmann, Stefan Thor, Johannes Bischof, Donald T. Fox, Marco Milán, the Bloomington and Kyoto Stock Center for sending flies and Uwe Walldorf for an aliquot of the anti-otp antibody as well as discussions. Help from Matthias Ostermaier with acquisition of data presented in Supplementary Figure 11 is also acknowledged. Beat Suter gratefully instructed us on how to dissect ovaries. We are indebted to Gina Evora, Consuelo Zuluaga Gomez, Adela Garzia and Patrick Groelly for constant and reliable supply of world’s best fly food. The work was supported by grants from the Swiss National Science Foundation (310030_192659 and 310030B_176400) and by funds from the Kanton Basel-Stadt and Basel-Land.

## Materials and Methods

### Fly stocks

Flies were housed in a 25°C incubator with 60% humidity and reared on a standard cornmeal agar diet (0.07% w/v granulated agar-agar, 0.1% w/v corn meal, 0.06% w/v granulated sugar, 0.015% w/v granulated dry yeast, 0.003% w/v granulated Nipagin). The solidified agar plug is supplemented with a drop of fresh baker’s yeast unless otherwise stated.

*ap^Xa^* is a classic *ap* allele (Serebrovsky and Dubinin, 1930) whose break in *ap* has been molecularly characterized (Bieli et al., 2015a). *ap^DG1^* is described in Gohl et al., 2008 (where it is called ap*^DG^*_)_. ap*^apRXa^*, ap*^RXa-Dad52.4^*, ap*^Dad52.4^*, ap*^c1.4b^*, ap*^attPΔEnh^*, ap*^DG3^* and *ap^DG8^* are described in Bieli et al 2015a. *ap^DG12^, ap^DG14^, ap^t11b^, ap^c1.2^, ap^cDNAint2.3^* and *ap^attPΔCDS^* are described in Bieli et al 2015b. *ap^MS1^*, *ap^MS1+dad13^, ap^MS2^, ap^c1.4b-Gal4^, ap^DG1-Gal4^, ap^R1+minLSE-Gal4^ and ap^MS3-Gal4^* are described in Reinger et al, 2025. *ap^R2^* is described in Aguilar et al., 2023. *ap^ΔapS-CRM^* was obtained from Johannes Stratmann and Stefan Thor. *y M{vas-int.Dm}zh-2A w* was obtained from Johannes Bischof. *UAS-ap* was obtained from Marco Milan. *P{GawB}Dad^NP0974^* (#103843; *Dad^Gal4^*) was purchased from the Kyoto Stock Center. The following stocks were purchased from the Bloomington Stock Center: *ap^4^* (#223), *ap^54^* (#3500), *ap^77f^* (#3495), *ap^ts78j^* (#3495), *ap^md544^* (#3041), *P{y[+t7.7][+mC]=10XUAS-IVS-mCD8::GFP}attP40* (#32186), *fkh^6^* (#28281), *otp^13064^* (#94293), *otp^1024^* (#94292) and *G-TRACE* (#28281).

The following two deletions were obtained by imprecise excision of insert *ap^MM^* (Gohl et al., 2008) during the generation of gene conversion events *ap^c1.2b^* and *ap^c2.73c^* (for details see Bieli et al 2015a):

*Df(2R)ap^c2.34d^*: contains a 3951 bp deletion starting in the 5’-UTR and ending in the large intron 1 (Fig. 1D). Sequences flanking the break points are CTTATATCGGTAGGTATACC-(Δ)-TACGCGCACCGCTTCCAAAA. At the *ap^MM^* insertion site, 35 bp of the P-element ends flanked by the 8 bp target site duplication (GGATGGAC) remain in place. Referred to as *ap^c2.34d^* in the text.

*Df(2R)ap^c1.72d^*: contains a 4477kb deletion ending in intron 1 at exactly the same bp as *ap^c2.34d^*, but it starts at the *ap^MM^* insertion site (Fig. 1D). Sequences flanking the break points are CTTATATCGGTAGGTATACC-(Δ)-<u>GGATGGAC</u>TCCAGACTCGAA. The 8 bp target site duplication on the distal side of *ap^MM^* is underlined. Between the two break points, the 3’ most 2138 bp of the *ap^MM^* insert remain in place. Referred to as *ap^c1.72d^* in the text.

The following two alleles were generated by Flp-mediated recombination (Golic and Golic, 1996) between FRT sites in transgene inserts located *in trans*:

*Df(2R)ap^DG10^*: FRT sites in transgenes *ap^MM^* and *ap^EE23.9^* (Bieli et al., 2015b) were used to generate this 2814 bp deletion (Fig. 1D). *ap^EE23.9^* is a ФC31-integrase mediated insertion of a plasmid containing *mini-white*, *FRT* and *mini-yellow* in docking site *MI00964* (Venken et al 2011). After recombination, *FRT* and *mini-yellow* remain in place. Can be maintained as a homozygous stock. Referred to as *ap^DG10^* in the text.

*Df(2R)ap^DG13^*: FRT sites in transgenes *ap^f00451^* (Thibault et al 2004) and *ap^DD8.1^* (Bieli et al., 2015b) were used to generate this 1110 bp deletion. *ap^DD8.1^* is a ФC31-integrase mediated insertion of a plasmid containing *mini-white*, *FRT* and *mini-yellow* in docking site *MI02330* (Venken et al., 2011). Note that the distal *attP* site of *MI02330* is missing (FBrf0243345). Therefore, distal to *ap^DG13^*, *FRT*, *mini-white* and the *yellow* and *EGFP* reporters of *Mi{MIC}* remain in place. Referred to as *ap^DG13^* in the text.

### Construction of new *ap^Gal4^* and *ap^LexA^* driver lines

*ap^LSEΔ3.2-Gal4^* and *ap^LSEΔ5-Gal4^* were constructed the same way as *ap^minLSE-Gal4^* described in a previous paper (Reinger et al., 2025). In short: plasmids pCR60 and pCR61 were integrated into the *ap^attPΔEnh^* landing site (Supp. Fig.1). In the final stock, the *yellow* marker present on pCR60 and pCR61 remains in place between the 3’ end of Gal4 and 5’ end of *l(2)09851*.

*ap^MS5-Gal4^* was obtained by Flp-mediated recombination between FRT sites in *ap^c1.4b-Gal4^* and *ap^R2^*. Note that the *mini-yellow* marker of *ap^c1.4b-Gal4^* is lost and that the distal breakpoint of *ap^MS2^* and *ap^MS5-Gal4^* are identical. Hence, the apE wing enhancer is deleted.

For the generation of *ap^c1.4b-LexA^*, plasmid pCR62 was produced by Genewiz. A cassette was synthesized that combines the minLSE, a synthetic core promotor (Pfeiffer et al 2008) and the LHV2 fragment of Yagi et al (2010). The cassette is flanked by NotI and a SpeI restriction sites and was cloned into pMS399 (Reinger et al 2025) cut with the same enzymes and pCR62 was obtained. *ap^c1.4b-LexA^* was generated by injection of plasmid pCR62 into *y w vas-integrase; ap^c1.4b^/CyO* embryos. pCR62 contains a *mini-yellow* marker devoid of all *yellow* enhancers. Injectees were crossed with *y w* partners. Transgenic flies could be detected by their y^+^ wings and a *y w; ap^c1.4b-LexA^/CyO* stock was established.

### Construction of ap^ex5-8^

The *ap* locus produces two proteins of 469 and 246 amino acids (see text and Supp. Material). In *ap^cDNAint2.3^* flies, only the long 469 aa peptide is produced (Bieli et al., 2015b). In order to elucidate the function of the short 246 aa peptide on its own with allele *ap^ex5-8^*, plasmid pCR76 was synthesized by Genewiz. pCR76 contains a 1786 bp NheI/AgeI fragment comprising the coding genomic DNA from exon 5 to exon 8 up to the NheI site (1279 bp) and the proximal 507 bp of the *ap* 5’-UTR up to the AgeI site. Then, the NheI/AgeI fragment of pCR76 was isolated and cloned into plasmid pDB92 cut with the same enzymes resulting in plasmid pCR77. In pCR77, the apPRE and the complete 5’-UTR are fused directly to the ATG start codon of the short peptide in exon 5. From the distal side, the whole interval starts with GTCCATCCTCATTCATGTGG. At the junction between 5’-UTR and exon 5 (ATG underlined), the sequence is AAAAGACGCAC<u>ATG</u>CGCGCCA. At the NheI site (underlined), the sequence is AGCACG<u>GCTAGC</u>TCTTGAAT. pCR77 was injected into *y M{vas-int*.*Dm}zh-2A w; ap^attPΔCDS^*/*CyO* embryos and allele *ap^ex5-8^* was obtained.

### Establishing recombinant stocks

Recombinants between genetic components *P{y[+t7.7][+mC]=10XUAS-IVS-mCD8::GFP}attP40*, *ap^DG3(w+)^* and *P{UAS-ap, w+}* were isolated according to standard meiotic recombination protocols. Presence of genetic components was verified with appropriate test crosses.

### Plasmids for *in vivo* rescue of deletion interval *ap^MS2^*

Ten DNA fragments (Fig. 2A) flanked by HindIII and SpeI restriction sites were amplified with appropriate PCR primers (see Suppl. Table 1). These fragments were cloned into reentry plasmid pDB345 cut with the same enzymes (Supp. Fig. 5C-C’). Resulting plasmids were validated by sequencing (Microsynth, Balgach). Midi-prep grade plasmid DNA was produced (Macherey-Nagel #740422.50) and injected into *y M{vas-int.Dm}zh-2A w; ap^MS2^/ CyO* embryos. Surviving injectees were crossed with *y w* flies and the offspring was selected for y^+^ wings. Stocks with genotype *y w; ap^MS2+FragX(y+)^/SM6a* were established. All new alleles were verified by sequencing using the PCR primers P307 and P309 (for sequences, see Supp. Table 1).

### *in silico* analysis of the minLSE

An *in silico* analysis on the minLSE was done to identify regions conserved between 27 closely related *Drosophila* species (UCSC Genome Browser (*D. melanogaster* Aug 2014, BDGP Release 6 + ISO1 MT/dm6; http://genome-euro.ucsc.edu/; University of California Santa Cruz)). In summary, six conserved regions were defined within the minLSE. Together they span most of the ∼560bp and are named as LSE-1 to LSE-6. The conserved regions were enlarged by an arbitrary amount of bps. In addition, two deletions were generated spanning the rest of the apLSE which were not covered by the conserved regions and their arbitrary extensions and were named LSE-7 and LSE-8 (Fig. 2C).

### Deletion mutagenesis of minLSE by Gibson Assembly

Oligonucleotide sequences synthesized for of Gibson Assembly can be found in Supp. Table 2. The chimeric primers were designed as described in the Gibson Assembly instruction manual. Therefore, they combined ∼15 bp of the distal and ∼15 bp of the proximal part directly adjacent of the sequences to be deleted. The outer, non-chimeric primers were designed in a way to share an overlapping sequence of ∼15-25 bp of the minLSE fragment followed by ∼15-25 bp of the vector sequence including a HindIII or a SpeI restriction site for cloning into pDB345 (see Supp. Fig. 5). These fragments (LSE-X.1 and LSE-X.2; Supp. Table 3) were amplified with standard PCR reactions and pDB345 was cut with HindIII and SpeI. All three compounds were mixed with the Gibson Assembly® Master Mix 2x (New England BioLabs, E2611) and were incubated at 50°C for 1h. 4 µL of this reaction was dialyzed and transformed into electro-competent *E. coli* cells. Midi-prep grade plasmid DNA was produced (Macherey-Nagel #740422.50) from clones verified by sequencing (Microsynth, Balgach). Plasmids were injected into the *y M{vas-int.Dm}zh-2A w;ap^MS2^/CyO* landing site. Surviving injectees were crossed with *y w* flies and the offspring was selected for y^+^ wings. Stocks with genotype *y w;ap^MS2+FragX(y+)^/SM6a* were established. All new alleles were verified by sequencing using the PCR primers P307 and P309.

### Point mutation analysis of the LSEΔ3.2_C region

#### 2bp point mutations

Survival assays identified the LSE-3.2C region as being of particular importance (Fig. 2D). Therefore, this region was scanned for critical bases by overlapping 2-bp point mutations. 14 mutations spanning the LSE-3.2C region were generated (plasmid library synthesized by Genewiz). Adenine was changed to Cytosine, Thymine to Guanine and vice versa (Fig. 4A). The plasmid library mix was injected into *y M{vas-int.Dm}zh-2A w;ap^MS2^/ CyO* embryos and transgenic flies were identified by expression of the *mini-yellow* marker. The 14 possible alleles (*ap^LSEΔ3.2_C_mut_1(y+)^* to *ap^LSEΔ3.2_C_mut_1(y+)^*) were identified by sequencing of hemizygous flies. *y w;ap^LSEΔ3.2_C_mut_X(y+)^/SM6* stocks were established from selected candidates.

#### Fkh cluster point mutations in LSEΔ3.2_C region

Since none of the induced 2 bp-point mutations indicated the existence of a relevant protein binding site in the *ap^LSEΔ3.2_C^* interval, JASPAR (https://jaspar.elixir.no/) was used to search for transcription factor binding sites within the LSE-3 region. The threshold was set to 70%. The software located a cluster of five fkh transcription factor binding sites in and around the LSE-3.2C region. The conserved motif within the predicted Fkh binding site, TTT, was mutated to CCC. In total, four plasmids carrying these point mutations in the 562 bp minLSE were synthesized by Genewiz (Fig. 4B). The plasmids were integrated into the *y M{vas-int.Dm}zh-2A w;ap^MS2^/ CyO* landing site and transgenic fly lines with genotype *y w;ap^LSE_fkh_variation_X(y+)^/SM6* stocks were established.

### Point mutation analysis of the LSEΔ5.1 region

In *ap^LSEΔ5.1^*, a putative Hox factor binding site is deleted (Fig. 4E). To corroborate this hypothesis, the motive TTAG was changed to GGCG. The mutated plasmid (CR84) was synthesized by Genewiz. It was integrated into the *y M{vas-int.Dm}zh-2A w; ap^MS2^/ CyO* landing site and a transgenic fly line with genotype *y w; ap^LSEΔ5.1_mut(y+)^/CyO,Dfd::YFP* stocks was established.

### *in vivo* rescue assays of *ap^LSE^* alleles

Before newly established *ap^LSE(y+)^* alleles were tested for their rescue potential, the *mini-yellow* marker present on the backbone of the reentry plasmid was removed by Flp-treatment and *y w; ap^LSE^/CyO,Dfd::YFP* stocks were established (Supp. Fig. 5D-E). The presence of the desired allele was again verified by sequencing. Then, the rescue potential of each *ap^LSE^* allele was determined by careful assessment of its survival rate. *y w; ap^LSE^/CyO, Dfd::YFP* males were crossed with *y w; ap^DG3^/CyO, Dfd::YFP* virgins. Hemizygous male and female flies were collected during one day and kept together at 25°C in pools of 20-30 flies. Flies were kept in glass vials (∼56 cm^3^) on normal cornmeal agar diet not supplemented with fresh baker’s yeast to prevent flies from sticking to the food. Flies were transferred to fresh tubes at least every second day without CO_2_ anesthesia. The number of surviving flies was scored every day or every third day during a period of 5, 6 or 18 days, respectively. Survival curves are plotted as survival in % ([survivors on day X]/[total of flies scored]x100) versus day X after eclosion.

### Sample preparation and immunohistochemistry

#### G-TRACE

Gal4 driver males were crossed with virgins collected from the real-time and lineage expression stock (G-TRACE; Evans et al., 2009) and Gal4 expression in the hindgut of adult progeny was monitored by confocal microscopy. Ampullae were dissected in PBS and fixed with 4% paraformaldehyde in PBS for 25 min at room temperature (RT). They were washed 3x for 20 min with PBST at RT and 3x with PBS and finally mounted in Vectashield containing DAPI (Vector Laboratories). Confocal imaging of adult guts was done with an Olympus SpinD. Image processing was done with the Omero or with the ImageJ software.

#### Immunostaining

Protein expression was investigated by antibody stainings. Embryos were dechorionated and fixed in 1:1 heptane:4% paraformaldehyde in 1xPBS for 25 min at room temperature (RT) with vigorous shaking (∼850 rpm). The fixative was removed and replaced by an equal amount of MeOH. The vitellin membrane was removed by vigorous shaking for 1 min. Embryos were washed 3x with MeOH. Then, standard immune-detection protocols were followed. Primary antibody: Rabbit α-Ap (1:800-1000, Bieli et al., 2015b). Guinea-pig α-Otp (1:1000, Hildebrandt et al., 2020). Secondary antibody: α-rabbit Alexa Fluor 568 and α-guinea-pig Alexa Fluor 488 used at 1:500. Samples were mounted in Vectashield containing DAPI. Confocal imaging of embryos was performed using a Leica SP5 or SP8 microscope. Image processing was done with the Omero or with the ImageJ software.

#### Ampulla preparations

Adult ampullae were dissected in 1xPBS on a siliconized slide and mounted in 1x PBS/50% Glycerol. Coverslips with tiny playdough feet were used to prevent tissue crushing. Preparations were immediately inspected with a Leica DM2700M microscope equipped with Normarski optics and imaged with a Leica Flexacam C3 camera.

#### Wing preparations

Adult female wings were removed from anesthetized flies and dipped for 10 sec. in 100% EtOH. Excess EtOH was removed with a tissue. Then, wings were transferred into a drop of Hoyer’s solution on a microcopy slide. They were mounted dorsal side up and a cover slip was applied. Preparations were baked at 58°C over night and weighed down with a 40-g metal cylinder. Images were taken with a Leica DM2700M microscope equipped a Leica Flexacam C3 camera.

#### Ovary preparation

2-3 days old, well-fed virgins were used for dissection. Ovaries were dissected in PBS and ovarioles were separated from each other with a sharp needle. Then, they were transferred into an Eppendorf tube containing 4% paraformaldehyde in 1xPBS and fixed for 20min at RT. The fixing solution was removed and ovaries were rinsed 3x with PBST followed by 2×15min washes at RT with PBST. Finally, the ovaries were quickly washed with PBS and mounted on a glass slide with Vectashield containing DAPI. Samples were imaged with a point scanning confocal Leica SP5-II-Matrix microscope.

### Ectopic expression of UAS transgenes with *byn^Gal4^*

A *byn^Gal4^/TM3,Sb* stock (Iwaki and Lengyel, 2002) was obtained from Donald T. Fox. It was recombined with *P{tub-Gal80^ts^}* (B#7019) and a *byn^Gal4^ tub-Gal80^ts^/TM6C* stock was established. The sources for the various UAS transgenes are indicated in Table 3.

Pre-3^rd^ larval instar lethality of *byn^Gal4^*>*UAS-ap* animals was deduced from the following two crosses:

1. *ap^MS2^/CyO;byn^Gal4^/TM2*,*Ubx* virgins and *ap^DG3^ UAS-ap UAS-CD8-GFP*/*CyO,Dfd::YFP* males (cross grown at 25°C).
2. *ap^MS2^/CyO, Dfd::YFP;byn^Gal4^ tub-Gal80^ts^/TM2, Ubx* virgins and *ap^DG3^ UAS-ap UAS-CD8-GFP/CyO, Dfd::YFP* males (cross grown at 29°C).

A control cross showed that a strong fluorescent signal is readily detectable in the hindgut and anal pads of *byn^Gal4^>UAS-CD8-GFP* larvae. No such animals were observed among the 264 3^rd^ instar larvae obtained from cross (1). In contrast, cross (2) produced *byn^Gal4^>UAS-CD8-GFP* larvae close to the expected Mendelian ratio (n=483; data not shown).

Data presented in Table 3 was obtained by crossing *byn^Gal4^ tub-Gal80^ts^/TM6C* virgins with males selected from the respective UAS transgene stocks. Two sets of crosses were set up. One was kept at 25°C constantly. For the other, embryos were collected at 18°C for 24 hrs. Parents were removed and the cultures were aged at 18°C for 144 hrs. Finally, cultures were shifted to 29°C when developing larvae were 144-168 hrs old. This temperature regime makes sure that *byn^Gal4^>UAS-transgene* expression kicks in sufficiently early to influence papilla formation during the first two days of metamorphosis (Wilson, 1981a; Reinger et al., 2025). Survival of progeny eclosing from these crosses was scored by determining the number of surviving flies after a 5-day period. Hindguts were dissected and documented as described above.

The *Bx^Gal4^* driver used in Fig. 6G corresponds to *^BxMS1096-KE^* (B#8696; Milan et al., 1998). The *Ser^Gal4^* driver was *P{UAS-Ser.mg5603}SS1* (B#5815).

## Notes

### Competing Interest Statement

The authors have declared no competing interest.

