## Supplementary Material for "Genetic characterization of the *apterous* Life Span Enhancer in *Drosophila melanogaster*"

#### Supp. Fig. 1

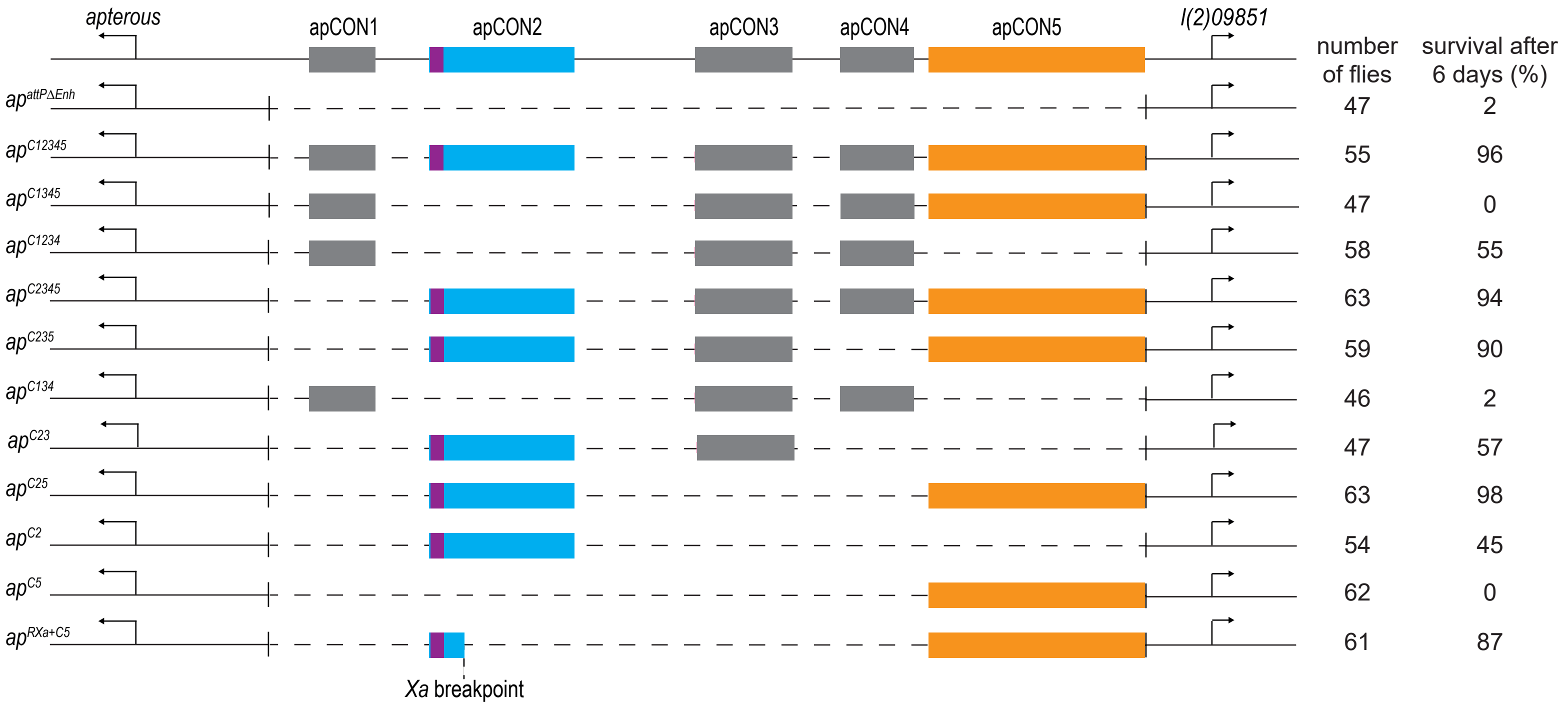

*apterous*
apCON1
apCON2
apCON3
apCON4
apCON5
*l(2)09851*
number of flies
survival after 6 days (%)
$ap^{attP\Delta Enh}$ 47 2
$ap^{C12345}$ 55 96
$ap^{C1345}$ 47 0
$ap^{C1234}$ 58 55
$ap^{C2345}$ 63 94
$ap^{C235}$ 59 90
$ap^{C134}$ 46 2
$ap^{C23}$ 47 57
$ap^{C25}$ 63 98
$ap^{C2}$ 54 45
$ap^{C5}$ 62 0
$ap^{RXa+C5}$ 61 87
*Xa* breakpoint

### Supp. Fig. 2

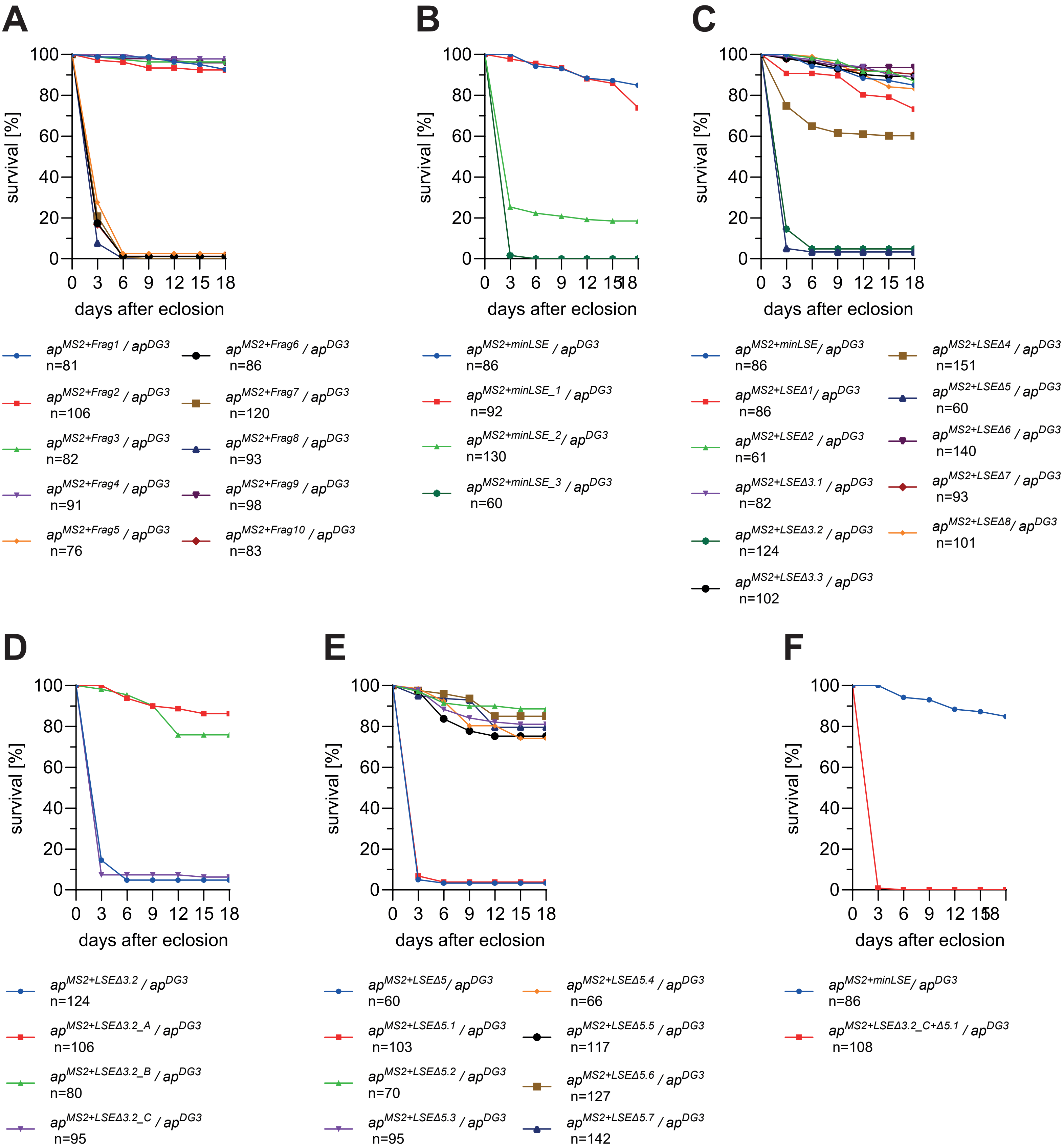
A
B
C
D
E
F
survival [%]
days after eclosion
$ap^{MS2+Frag1}$ / $ap^{DG3}$ n=81
$ap^{MS2+Frag2}$ / $ap^{DG3}$ n=106
$ap^{MS2+Frag3}$ / $ap^{DG3}$ n=82
$ap^{MS2+Frag4}$ / $ap^{DG3}$ n=91
$ap^{MS2+Frag5}$ / $ap^{DG3}$ n=76
$ap^{MS2+Frag6}$ / $ap^{DG3}$ n=86
$ap^{MS2+Frag7}$ / $ap^{DG3}$ n=120
$ap^{MS2+Frag8}$ / $ap^{DG3}$ n=93
$ap^{MS2+Frag9}$ / $ap^{DG3}$ n=98
$ap^{MS2+Frag10}$ / $ap^{DG3}$ n=83
$ap^{MS2+minLSE}$ / $ap^{DG3}$ n=86
$ap^{MS2+minLSE\_1}$ / $ap^{DG3}$ n=92
$ap^{MS2+minLSE\_2}$ / $ap^{DG3}$ n=130
$ap^{MS2+minLSE\_3}$ / $ap^{DG3}$ n=60
$ap^{MS2+minLSE}$/ $ap^{DG3}$ n=86
$ap^{MS2+LSE\Delta1}$/ $ap^{DG3}$ n=86
$ap^{MS2+LSE\Delta2}$ / $ap^{DG3}$ n=61
$ap^{MS2+LSE\Delta3.1}$ / $ap^{DG3}$ n=82
$ap^{MS2+LSE\Delta3.2}$ / $ap^{DG3}$ n=124
$ap^{MS2+LSE\Delta3.3}$ / $ap^{DG3}$ n=102
$ap^{MS2+LSE\Delta4}$ / $ap^{DG3}$ n=151
$ap^{MS2+LSE\Delta5}$ / $ap^{DG3}$ n=60
$ap^{MS2+LSE\Delta6}$ / $ap^{DG3}$ n=140
$ap^{MS2+LSE\Delta7}$ / $ap^{DG3}$ n=93
$ap^{MS2+LSE\Delta8}$/ $ap^{DG3}$ n=101
$ap^{MS2+LSE\Delta3.2}$ / $ap^{DG3}$ n=124
$ap^{MS2+LSE\Delta3.2\_A}$ / $ap^{DG3}$ n=106
$ap^{MS2+LSE\Delta3.2\_B}$ / $ap^{DG3}$ n=80
$ap^{MS2+LSE\Delta3.2\_C}$ / $ap^{DG3}$ n=95
$ap^{MS2+LSE\Delta5}$/ $ap^{DG3}$ n=60
$ap^{MS2+LSE\Delta5.1}$ / $ap^{DG3}$ n=103
$ap^{MS2+LSE\Delta5.2}$ / $ap^{DG3}$ n=70
$ap^{MS2+LSE\Delta5.3}$ / $ap^{DG3}$ n=95
$ap^{MS2+LSE\Delta5.4}$ / $ap^{DG3}$ n=66
$ap^{MS2+LSE\Delta5.5}$ / $ap^{DG3}$ n=117
$ap^{MS2+LSE\Delta5.6}$ / $ap^{DG3}$ n=127
$ap^{MS2+LSE\Delta5.7}$ / $ap^{DG3}$ n=142
$ap^{MS2+minLSE}$/ $ap^{DG3}$ n=86
$ap^{MS2+LSE\Delta3.2\_C+\Delta5.1}$ / $ap^{DG3}$ n=108

### Supp. Fig. 3

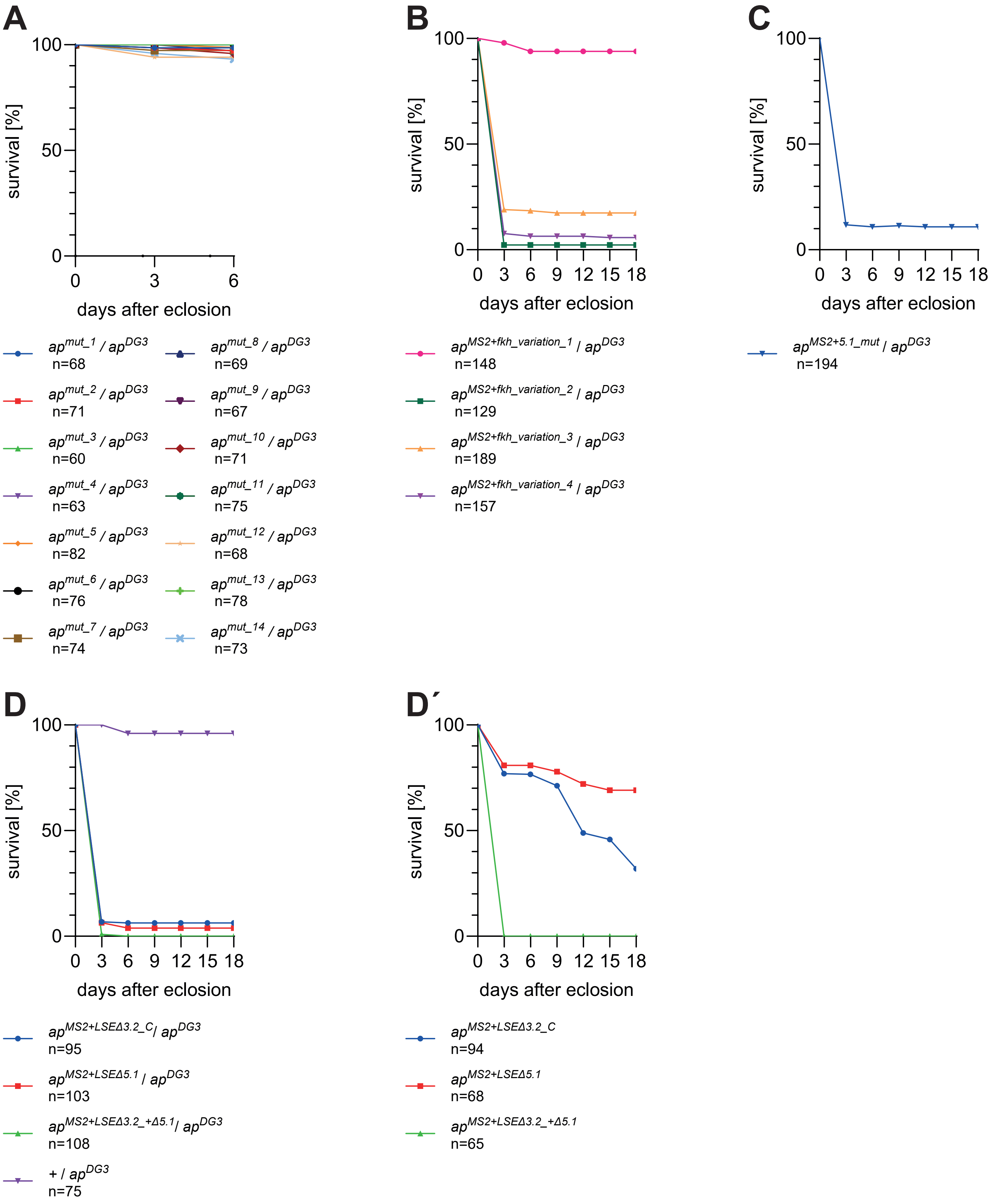
A
survival [%]
100
50
0
0
3
6
days after eclosion
$ap^{mut\_1}$ / $ap^{DG3}$ n=68
$ap^{mut\_2}$ / $ap^{DG3}$ n=71
$ap^{mut\_3}$ / $ap^{DG3}$ n=60
$ap^{mut\_4}$ / $ap^{DG3}$ n=63
$ap^{mut\_5}$ / $ap^{DG3}$ n=82
$ap^{mut\_6}$ / $ap^{DG3}$ n=76
$ap^{mut\_7}$ / $ap^{DG3}$ n=74
$ap^{mut\_8}$ / $ap^{DG3}$ n=69
$ap^{mut\_9}$ / $ap^{DG3}$ n=67
$ap^{mut\_10}$ / $ap^{DG3}$ n=71
$ap^{mut\_11}$ / $ap^{DG3}$ n=75
$ap^{mut\_12}$ / $ap^{DG3}$ n=68
$ap^{mut\_13}$ / $ap^{DG3}$ n=78
$ap^{mut\_14}$ / $ap^{DG3}$ n=73
B
survival [%]
100
50
0
0 3 6 9 12 15 18
days after eclosion
$ap^{MS2+fkh\_variation\_1}$ / $ap^{DG3}$ n=148
$ap^{MS2+fkh\_variation\_2}$ / $ap^{DG3}$ n=129
$ap^{MS2+fkh\_variation\_3}$ / $ap^{DG3}$ n=189
$ap^{MS2+fkh\_variation\_4}$ / $ap^{DG3}$ n=157
C
survival [%]
100
50
0
0 3 6 9 12 15 18
days after eclosion
$ap^{MS2+5.1\_mut}$ / $ap^{DG3}$ n=194
D
survival [%]
100
50
0
0 3 6 9 12 15 18
days after eclosion
$ap^{MS2+LSE\Delta3.2\_C}$/ $ap^{DG3}$ n=95
$ap^{MS2+LSE\Delta5.1}$ / $ap^{DG3}$ n=103
$ap^{MS2+LSE\Delta3.2\_+\Delta5.1}$/ $ap^{DG3}$ n=108
+ / $ap^{DG3}$ n=75
D´
survival [%]
100
50
0
0 3 6 9 12 15 18
days after eclosion
$ap^{MS2+LSE\Delta3.2\_C}$ n=94
$ap^{MS2+LSE\Delta5.1}$ n=68
$ap^{MS2+LSE\Delta3.2\_+\Delta5.1}$ n=65

Supp. Fig. 4

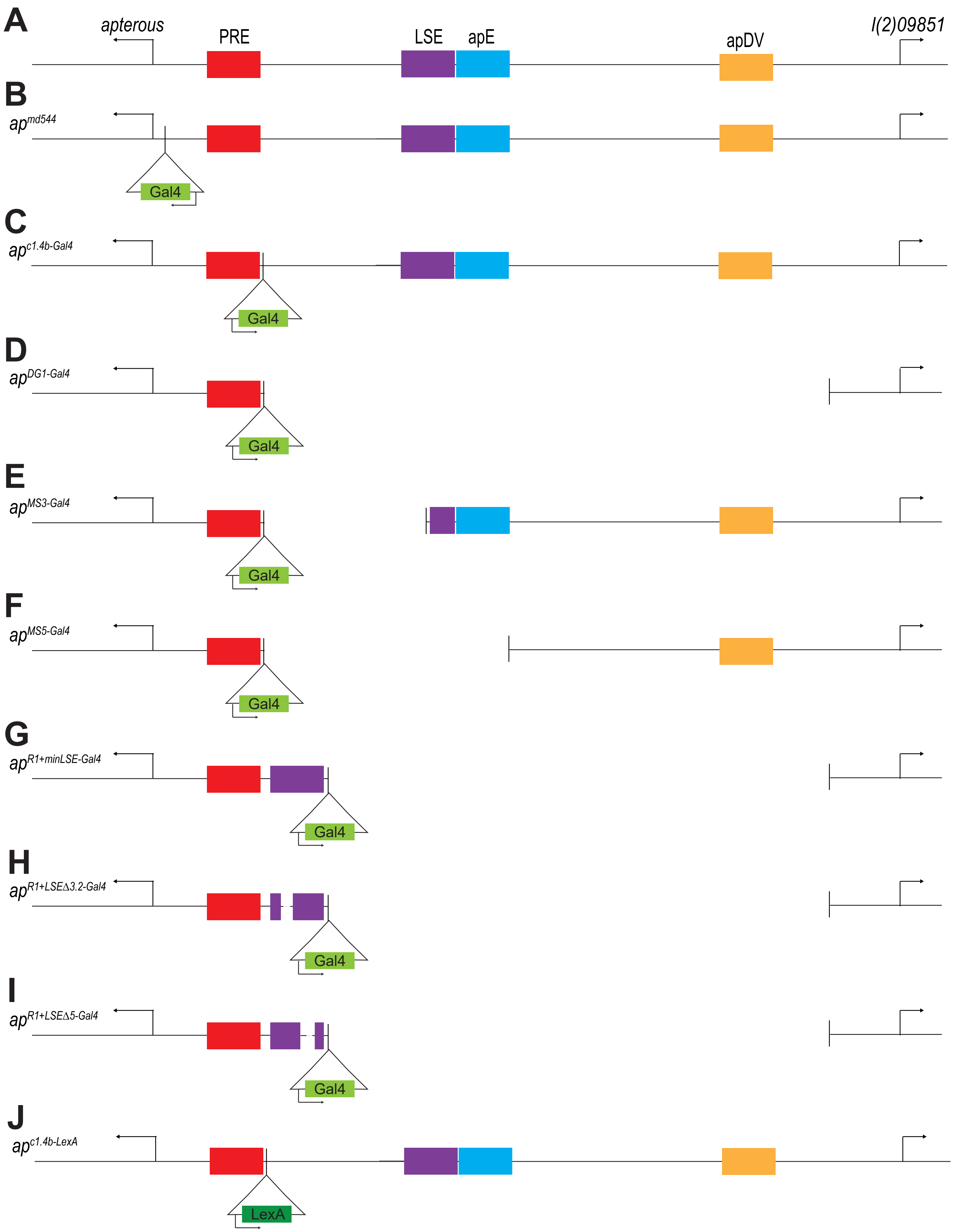
A
apterous
PRE
LSE
apE
apDV
l(2)09851
B
ap md544
Gal4
C
ap c1.4b-Gal4
Gal4
D
ap DG1-Gal4
Gal4
E
ap MS3-Gal4
Gal4
F
ap MS5-Gal4
Gal4
G
ap R1+minLSE-Gal4
Gal4
H
ap R1+LSEΔ3.2-Gal4
Gal4
I
ap R1+LSEΔ5-Gal4
Gal4
J
ap c1.4b-LexA
LexA

#### Supp. Fig. 5

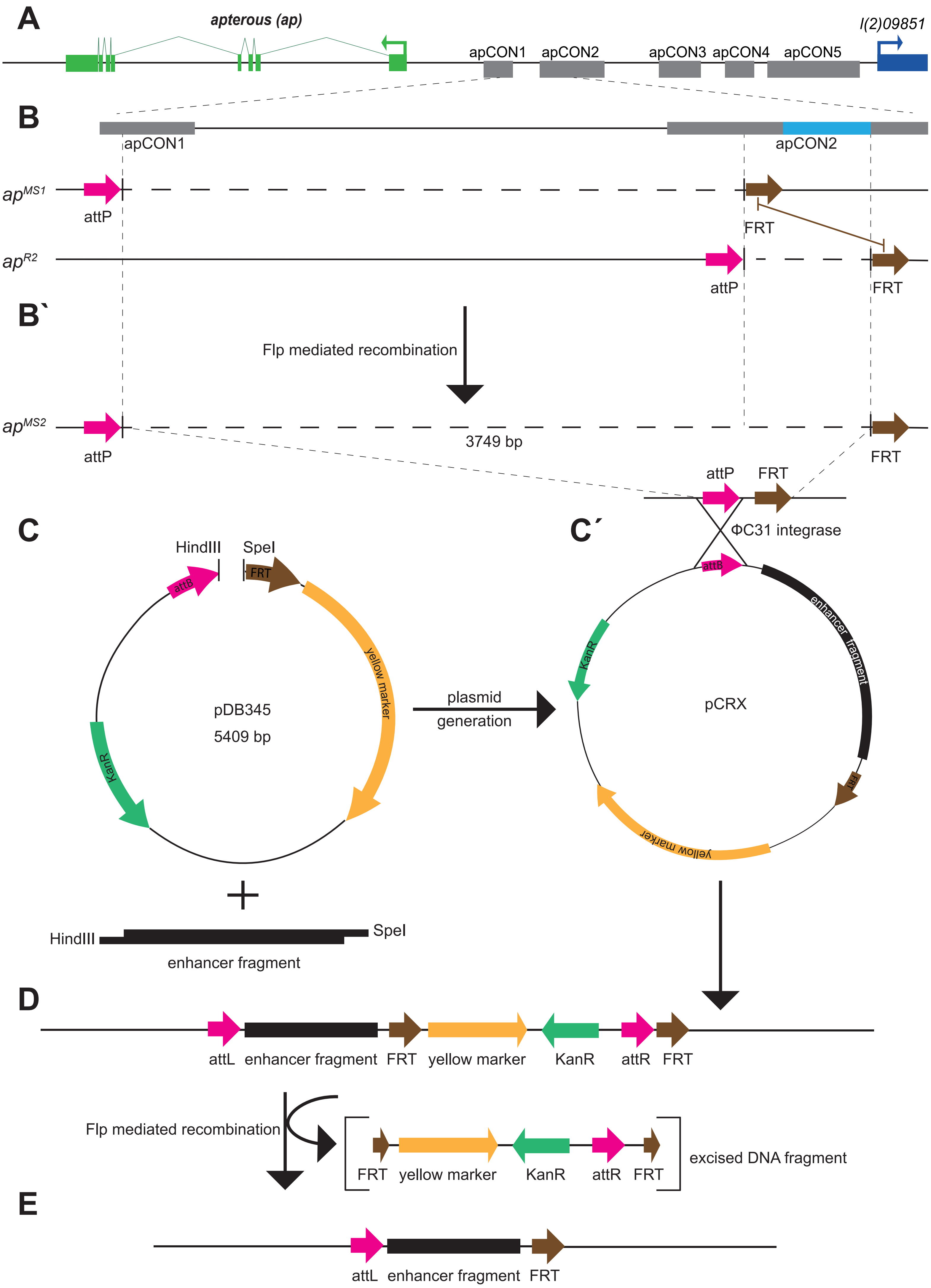

A
apterous (ap)
l(2)09851
apCON1
apCON2
apCON3
apCON4
apCON5
B
apCON1
apCON2
ap^MS1
attP
FRT
ap^R2
attP
FRT
B`
Flp mediated recombination
ap^MS2
attP
3749 bp
FRT
attP
FRT
ΦC31 integrase
C
HindIII
SpeI
attB
FRT
yellow marker
KanR
pDB345
5409 bp
plasmid
generation
C´
attB
enhancer fragment
KanR
pCRX
FRT
yellow marker
HindIII
SpeI
enhancer fragment
D
attL
enhancer fragment
FRT
yellow marker
KanR
attR
FRT
Flp mediated recombination
FRT
yellow marker
KanR
attR
FRT
excised DNA fragment
E
attL
enhancer fragment
FRT

**Supp. Figure 1: Rescue of survival by *ap* alleles affecting the 27 kb intergenic spacer.**

At the top, the ~27 kb intergenic spacer separating *ap* (on the left) and *l(2)09851* (on the right) is depicted. The five conserved regions apCON1 to apCON5 (Bieli et al., 2015b) are indicated by grey or colored boxes. apCON2 is shown in blue and apCON5 in orange. The box in magenta in apCON2 represents apLSE. DNA represented by dashed lines connecting the apCON elements are not present in the rescue constructs. The two columns on the right show the survival data for each allele as determined in hemizygous flies (over $ap^{DG3}$). A sub-fragment of apCON2 termed $ap^{RXa}$ represents the smallest fragment tested with this assay. Importantly, it contains the apLSE. Its distal end corresponds to the break point in $ap^{Xa}$.

**Supp. Figure 2: Survival curves for $ap^{LSE}$ alleles**

Survival of hemizygous flies was scored over a period of 18 days. **A**: data for alleles presented in Fig. 2A are shown. >80% of flies with a functional apLSE survive longer than 18 days. In the absence of apLSE activity, >80% of the flies are dead after 3 days. **B**: data for alleles presented in Fig. 2B are shown. Allele $ap^{MS2+minLSE2}$ retains only residual rescue activity. **C**: data for alleles presented in Fig. 2C are shown. Survival function is almost completely abolished in alleles $ap^{MS2+LSE\Delta3.2}$ and $ap^{MS2+LSE\Delta5.1}$, indicating that they affect important regulatory modules within apLSE. Reduced survival in $ap^{MS2+\ LSE\Delta4}$ flies could indicate that spacing between LSE3.2 and LSE5.1 could play a minor role. **D**: data for alleles presented in Fig. 2D are shown. Precocious adult death is only observed with $ap^{MS2+LSE\Delta3.2\_C}$. **E**: data for alleles presented in Fig. 2E are shown. Precocious adult death is only observed with $ap^{MS2+LSE\Delta5.1}$. **F**: data for allele presented in Fig. 2F is shown. It is concluded that apLSE function is dependent on regulatory modules defined by the 11 bp deletion $ap^{MS2+LSE\Delta3.2\_C}$ and the 6 bp deletion $ap^{MS2+LSE\Delta5.1}$.

**Supp. Figure 3: Survival curves for $ap^{LSE3.2\_C}$ and $ap^{LSE5.1}$ derived alleles**

**A-D**: Survival of hemizygous flies was scored over a period of 18 days. **A**: data for alleles presented in Fig. 4A are shown. None of the overlapping 2 bp point mutations affects LSE function. **B**: data for alleles presented in Fig. 4B are shown. Mutation of three or more of the putative Fkh protein binding sites results in a strong reduction of apLSE activity. **C**: data for allele presented in Fig. 4C is shown. Mutation of the putative Hox binding site results in a strong reduction of LSE activity. **D**: data for alleles presented in Fig. 4D are shown. **D'**: Survival of homozygous flies presented in Fig. 4D are shown. The hypomorphic nature of alleles $ap^{MS2+LSE\Delta3.2}$ and $ap^{MS2+\ LSE\Delta5.1}$ is indicated by the fact that homozygotes survive much better than hemizygotes. But concomitant mutation of both regulatory modules in allele $ap^{MS2+LSE\Delta3.2+\ \Delta5.1}$ abolishes apLSE activity in both genotypes, indicating that it is an $apLSE^{null}$.

**Supp. Figure 4: Overview of available $ap^{Gal4}$ drivers**

**A**: The ~27 kb intergenic spacer separating *ap* (on the left) and *l(2)09851* (on the right) is depicted. Relevant cis-regulatory elements shown are apPRE (in red), apLSE (in magenta), apE (in blue) and apDV (in yellow). Map is not shown to scale. **B**: the classic *ap*$^{Gal4}$ driver *ap*$^{md544}$ is depicted (Calleja et al., 1996). It contains a *P{GawB}* insertion (indicated by a triangle and a green box representing Gal4; Brand and Perrimon, 1993) very close to the transcription start site which almost completely inactivates the gene. **C**: allele *ap*$^{c1.4b\text{-}Gal4}$ is shown (Reinger et al., 2025). It contains a Gal4 gene distal to the apPRE in the *ap*$^{c1.4b}$ docking site. In contrast to *ap*$^{md544}$, hemizygous *ap*$^{c1.4b\text{-}Gal4}$ are well viable and fertile and develop normal wings. **D**: *ap*$^{DG1\text{-}Gal4}$ lacks the 27 kb intergenic spacer and therefore also all relevant enhancers (Reinger et al., 2025). **E-F**: *ap*$^{MS3\text{-}Gal4}$ and *ap*$^{MS5\text{-}Gal4}$ are derivatives of *ap*$^{c1.4b\text{-}Gal4}$ (for details see materials and methods). While *ap*$^{MS3\text{-}Gal4}$ contains a partially functional apLSE (it retains the LSE5.1 module) and an intact apE (Reinger et al., 2025), these regulatory elements are missing in *ap*$^{MS5\text{-}Gal4}$ (this study). **G-I**: these Gal4 drivers were obtained by insertion of minLSE-Gal4, LSEΔ3.2-Gal4 and LSEΔ5-Gal4 cassettes into the *ap*$^{attP\Delta Enh}$ docking site located at the same position as that of *ap*$^{c1.4b}$ (Reinger et al., 2025 and this study). All regulatory elements of the 27 kb intergenic spacer are absent and replaced by only the apLSE or derivatives, Gal4 and the *mini-yellow* marker (not shown). **J**: allele *ap*$^{c1.4b\text{-}LexA}$ is shown. For the construction of this allele, the LHV2 fragment of Yagi et al., 2010 consisting of the DNA-binding domain of LexA and the activator domain of Gal4 was used. Note that the Gal4 and LexA genes in C-J are transcribed under the control of the same synthetic core promoter described in Pfeiffer et al., 2008.

**Supp. Figure 5: Construction of the *ap*$^{MS2}$ landing site and its use in establishing *ap*$^{MS2+LSE}$ alleles**

**A**: The *ap* locus is depicted. On the left, the exon-intron structure of the gene is indicated in green. The conserved regions apCON1 to apCON5 are shown as grey boxes. To the left, the 5' end of gene *l(2)09851* is indicated in dark blue. **B**: A zoom-in of the apCON1-apCON2 interval is shown. The position of apE is indicated as a blue box. Below, allele *ap*$^{MS1}$ is shown. The dashed line represents the DNA missing in this deletion. On its left, there is an attP site (arrow in magenta) for ΦC1-integrase mediated insertion of attB containing plasmids (Bischof et al., 2007). On its left, there is a FRT site (brown arrow). This FRT was used to extend the *ap*$^{MS1}$ deletion on its distal side to the FRT of *ap*$^{R2}$ (described in Aguilar et al., 2023). **B'**: the resulting new deletion is *ap*$^{MS2}$ (3749 bp long). It contains the same genetic components as *ap*$^{MS1}$ but it also deletes the apE wing enhancer. **C**: Cloning of plasmids for ΦC1-integrase mediated insertion. Putative enhancer elements were cloned as HindIII/SpeI-fragments into plasmid pDB345 cut with the same enzymes. **C'**: Plasmids ready for ΦC1-integrase mediated insertion are called pCRX where X represents a number. They contain the *mini-yellow* transformation marker. Such plasmids were injected into *y M{vas-int.Dm}zh-2A w;ap*$^{MS2}$*/CyO* embryos and successful integration was monitored by the $y^+$ wing phenotype. **D**: the genetic structure of a newly established allele still containing the *mini-yellow* marker is shown. The cassette flanked by FRT sites (brown arrows) was deleted by Flp-mediated recombination (Golic and Golic, 1996). **E**: final structure of *ap*$^{LSE}$ alleles used for genetic testing. The enhancer fragment is flanked by a 54 bp attL and a 48 bp FRT.

#### Supp. Table 1: Plasmids generated by restriction cloning

| Name of final plasmid* | PCR Primer 1** | PCR Primer 2** |
|---|---|---|
| CR1(LSE-Fragment1) | tgtaagcttgcattttccatcggcgttttg | gcgactagtggaggccagctctaaaatcgag |
| CR2(LSE-Fragment2) | tgtaagcttgcattttccatcggcgttttg | gcgactagtaggaataaaaggaggcaagcctc |
| CR3(LSE-Fragment3) | tgtaagcttgcattttccatcggcgttttg | gcgactagtgcgtaaggattccatgaaccttc |
| CR4(LSE-Fragment4) | tgtaagcttgcattttccatcggcgttttg | gcgactagtatttgaatcgattcgaggggg |
| CR5(LSE-Fragment5) | tgtaagctttccagctctgaatctcactccc | gcgactagtggaggccagctctaaaatcgag |
| CR6(LSE-Fragment6) | tgtaagcttgcattttccatcggcgttttg | gcgactagtgcggggtaacttgatttgtttg |
| CR7(LSE-Fragment7) | tgtaagcttgaacaagaccctctcacctacg | gcgactagtggaggccagctctaaaatcgag |
| CR8(LSE-Fragment8) | tgtaagcttgcattttccatcggcgttttg | gcgactagtagaaatcccgtgttcattaaagtt |
| CR9(LSE-Fragment9) | tgtaagcttcacctgactcaatagcaagactac | gcgactagtggaggccagctctaaaatcgag |
| CR10(LSE-Fragment10) | tgtaagcttccgattccgatacttgag | atatctcgaggaagctgcactatgactgagg |
| CR24(minLSE) | tgtaagcttgaacctttttgttcctatacccc | gcgactagtgcgtaaggattccatgaaccttc |
| CR25(minLSE1) | tgtaagcttcacctgactcaatagcaagactac | gcgactagtatttgaatcgattcgaggggg |
| CR50(minLSE2) | attaagcttccctgtaaaggcagtcgaaaac | gcgactagtatcgattcgagggggctac |
| CR39(minLSE3) | agtaagcttcaaatagacaaacaaatcaagttaccc | agtactagtgattttttgtcgggccttgc |
| CR46(Dad13) | gtgaagcttcctcaaccttaaattgttgattcttcaacttg | gtgactagtcgaacgggagagcgcc |
| CR49(Dad13_mut) | gtgaagcttcctcaaccttaaattgttgattcttcaacttg | gtgactagtcgaacgggagagcgcc |

| Primer name for verification | sequence |
|---|---|
| project_LSE_sfp_for_P(307) | gtaaaaacatcgtcccgaacg |
| project_LSE_sfp_rev2_P(309) | gcttatgctgttcccatagatcg |

* Name in parenthesis used in main text.
**All cutting sites within the primer sequences are underlined.

**Supp. Table 2: Overview of plasmids generated with the Gibson Assembly Method**

| Final plasmid | Forward primer 1 | Reverse primer 1 | Forward primer 2 | Reverse primer 2 |
|---|---|---|---|---|
| CR26(LSEΔ1) | cgtactccacctcgaattcaagcttgaaccttttgttcctataccccc | atttgggtctggaagttaataaacggctatttaggttc | ggaacttccggatcccactagtgcgtaaggattccatgaacc | gtttattaacttccagacccaaatgacg |
| CR27(LSEΔ2) | cgtactccacctcgaattcaagcttgaaccttttgttcctataccccc | gggaggacaagtattaatgtaaaatctcgggtttgtcgattg | ggaacttccggatcccactagtgcgtaaggattccatgaacc | caatcgacaaacccgagattttacattaatacttgtcctccc |
| CR41(LSEΔ3.1) | cgtactccacctcgaattcaagcttgaaccttttgttcctataccccc | cttgatttgtttgtctatttgcctttacagggaggacaag | ggaacttccggatcccactagtgcgtaaggattccatgaacc | cttgtcctccctgtaaaggcaaatagacaaacaaatcaag |
| CR28(LSEΔ3.2) | cgtactccacctcgaattcaagcttgaaccttttgttcctataccccc | gggtcttgttcggggtaattcgcctccgactctttg | ggaacttccggatcccactagtgcgtaaggattccatgaacc | caaagagtcggaggcgaattaccccgaacaagaccc |
| CR57(LSEΔ3.2_A) | cgtactccacctcgaattcaagcttgaaccttttgttcctataccccc | gtcttgttcggggtaacttgattctatttgttcgcctccgactc | ggaacttccggatcccactagtgcgtaaggattccatgaacc | gagtcggaggcgaacaaatagaatcaagttaccccgaacaagac |
| CR63(LSEΔ3.2_B) | cgtactccacctcgaattcaagcttgaaccttttgttcctataccccc | gttcggggtaacttgatttgtttcgcctccgactctttg | ggaacttccggatcccactagtgcgtaaggattccatgaacc | caaagagtcggaggcgaaacaaatcaagttaccccgaac |
| CR64(LSEΔ3.2_C) | cgtactccacctcgaattcaagcttgaaccttttgttcctataccccc | agggtcttgttcggggtaattgtctatttgttcgcctccg | ggaacttccggatcccactagtgcgtaaggattccatgaacc | cggaggcgaacaaatagacaattaccccgaacaagaccc |
| CR42(LSEΔ3.3) | cgtactccacctcgaattcaagcttgaaccttttgttcctataccccc | ctatgactgagggggtcatacttgatttgtttgtctatttg | ggaacttccggatcccactagtgcgtaaggattccatgaacc | caaatagacaaacaaatcaagtatgaccccctcagtcatag |
| CR29(LSEΔ4) | cgtactccacctcgaattcaagcttgaaccttttgttcctataccccc | ggctgataatcagtcataaagcgtagatgag | ggaacttccggatcccactagtgcgtaaggattccatgaacc | acgctttatgactgattatcagcccaattc |
| CR31(LSEΔ5) | cgtactccacctcgaattcaagcttgaaccttttgttcctataccccc | gcttcgatgaaacagtccggtacataagtcg | ggaacttccggatcccactagtgcgtaaggattccatgaacc | tgtaccggactgtttcatcgaagcgggtg |
| CR51(LSEΔ5.1) | cgtactccacctcgaattcaagcttgaaccttttgttcctataccccc | gaaggcaatttagaattgggctcagtccggtacataagtcg | ggaacttccggatcccactagtgcgtaaggattccatgaacc | cgacttatgtaccggactgagcccaattctaaattgccttc |
| CR52(LSEΔ5.2) | cgtactccacctcgaattcaagcttgaaccttttgttcctataccccc | ctcaagaatgacttcatgaagggaattgggctgataatcagtccg | ggaacttccggatcccactagtgcgtaaggattccatgaacc | cggactgattatcagcccaattcccttcatgaagtcattcttgag |
| CR53(LSEΔ5.3) | cgtactccacctcgaattcaagcttgaaccttttgttcctataccccc | gattgactttactcaagaatgactggcaatttagaattgggctgataatcag | ggaacttccggatcccactagtgcgtaaggattccatgaacc | ctgattatcagcccaattctaaattgccagtcattcttgagtaaagtcaatc |
| CR54(LSEΔ5.4) | cgtactccacctcgaattcaagcttgaaccttttgttcctataccccc | cgggccttgcttgattgactcatgaaggcaatttagaattgggc | ggaacttccggatcccactagtgcgtaaggattccatgaacc | gcccaattctaaattgccttcatgagtcaatcaagcaaggcccg |
| CR55(LSEΔ5.5) | cgtactccacctcgaattcaagcttgaaccttttgttcctataccccc | gattttttgtcgggccttgcactttactcaagaatgacttcatg | ggaacttccggatcccactagtgcgtaaggattccatgaacc | catgaagtcattcttgagtaaagtgcaaggcccgacaaaaaatc |
| CR65(LSEΔ5.6) | cgtactccacctcgaattcaagcttgaaccttttgttcctataccccc | gacttcatgaaggcaatttatgataatcagtccggtacataagtcg | ggaacttccggatcccactagtgcgtaaggattccatgaacc | cgacttatgtaccggactgattatcataaattgccttcatgaagtc |

| | | | | |
|---|---|---|---|---|
| CR66(LSEΔ5.7) | cgtactccacctcgaattcaagcttgaaccttttgttcctatacccc | ctacacccgcttcgatgaaacttgattgactttactcaagaatgacttc | ggaacttccggatcccactagtgcgtaaggattccatgaacc | gaagtcattcttgagtaaagtcaatcaagtttcatcgaagcgggtgtag |
| CR30(LSEΔ6) | cgtactccacctcgaattcaagcttgaaccttttgttcctatacccc | ggaacttccggatcccactagtatttgaatcgattcgagggggc | | |
| CR75(LSEΔ7) | gaaggcaatttagaattgggctcagtccggtacataagtcg | cgactctttgttttcgactgggtaactggtcatctgaggc | ggaacttccggatcccactagtgcgtaaggattccatgaacc | gcctcagatgaccagttacccagtcgaaaacaaagagtcg |
| CR67(LSEΔ3.2_C+LSEΔ5.1) | cgtactccacctcgaattcaagcttgaaccttttgttcctatacccc | gttcggggtaacttgatttgtttcgcctccgactctttg | ggaacttccggatcccactagtgcgtaaggattccatgaacc | cgacttatgtaccggactgagcccaattctaaattgccttc |
| | gaaggcaatttagaattgggctcagtccggtacataagtcg | caaagagtcggaggcgaaacaaatcaagttaccccgaac | | |
| CR37(MS399[Gal4] +minLSE) | gacggtcacggcgggcatgcgaacctttgttcctatacccc | ggcgagctcggccgggctaggcgtaaggattccatgaacc | | |
| CR38(minLSE-Gal4) | agaaagtataggaacttcgcgaacctttgttcctatacc | gcttccacaaatcagatccattactcttttttgggtttggtgg | | |
| CR58(MS399[Gal4] + LSEΔ3.2) | gacggtcacggcgggcatgcgaacctttgttcctatacccc | ggcgagctcggccgggctaggcgtaaggattccatgaacc | | |
| CR61(LSEΔ3.2-Gal4) | agaaagtataggaacttcgcgaacctttgttcctatacc | gcttccacaaatcagatccattactcttttttgggtttggtgg | | |
| CR58(MS399[Gal4] +LSEΔ5) | gacggtcacggcgggcatgcgaacctttgttcctatacccc | ggcgagctcggccgggctaggcgtaaggattccatgaacc | | |
| CR60(LSEΔ5-Gal4) | agaaagtataggaacttcgcgaacctttgttcctatacc | gcttccacaaatcagatccattactcttttttgggtttggtgg | | |

**Supp. Table 3: Overview of LSEΔ1 to LSEΔ8**

| Deletion | Size of deletion | Sequence proximal to breakpoint | Sequence distal to breakpoint |
|---|---|---|---|
| LSEΔ1 | 66 | aaatagccgtttattaactt | ccagacccaaatgacgcctc |
| LSEΔ2 | 35 | tgcaatcgacaaaccccgag | attttacattaatacttgtc |
| LSEΔ3.1 | 27 | acttgtcctccctgtaaagg | caaatagacaaacaaatcaa |
| LSEΔ3.2 | 21 | aacaaagagtcggaggcgaa | ttaccccgaacaagaccctc |
| LSEΔ3.2_A | 7 | gtcggaggcgaacaaataga | atcaagttaccccgaacaag |
| LSEΔ3.2_B | 10 | ggaggcgaacaaatagacaa | ttaccccgaacaagaccctc |
| LSEΔ3.2_C | 11 | aacaaagagtcggaggcgaa | acaaatcaagttaccccgaa |
| LSEΔ3.3 | 33 | aatagacaaacaaatcaagt | atgaccccctcagtcatagt |
| LSEΔ4 | 85 | tctcatctacgctttatgac | tgattatcagcccaattcta |
| LSEΔ5 | 79 | gcgacttatgtaccggactg | tttcatcgaagcgggtgtag |
| LSEΔ5.1 | 6 | cgacttatgtaccggactga | gcccaattctaaattgcctt |
| LSEΔ5.2 | 9 | actgattatcagcccaattc | ccttcatgaagtcattcttg |
| LSEΔ5.3 | 7 | cagcccaattctaaattgcc | agtcattcttgagtaaagtc |
| LSEΔ5.4 | 7 | attctaaattgccttcatga | gtcaatcaagcaaggcccga |
| LSEΔ5.5 | 17 | aagtcattcttgagtaaagt | gcaaggcccgacaaaaaatc |
| LSEΔ5.6 | 7 | atgtaccggactgattatca | taaattgccttcatgaagtc |
| LSEΔ5.7 | 20 | tcttgagtaaagtcaatcaa | gtttcatcgaagcgggtgta |
| LSEΔ6 | 53 | cccctcgaatcgattcaaat | tgaataatttcaacatatcg |
| LSEΔ7 | 33 | gcctcagatgaccagttacc | cagtcgaaaacaaagagtcg |
| LSEΔ8 | 42 | gcaaggcccgacaaaaaatc | tccagctctgaatctcactc |

**Supplementary material:**

#### Liquid chromatography-Mass spectrometric (LC-MS) analysis of *ap* alleles

##### Material and Methods

Sample preparation:

Homozygous or *CyO, Dfd::YFP* balanced *apterous* alleles were used to generate the desired genotypes. A fluorescent bino served to select against the Dfd::YFP marker in 3rd instar larvae. Selected YFP- larvae were washed in Ringer's solution and dissected in 1xPBS. One tissue sample for proteomic analysis contained the central nervous system and attached imaginal disc tissue of four larvae. Dissected tissues were separated from the rest of the carcasses and transferred into a 2 ml Eppendorf tube on ice containing 50 µl of 1xPBS. After a brief centrifugation step, the supernatant was removed and tissue samples were immediately frozen on dry ice. Samples were kept at -80ºC until they were further processed for proteomic analysis. Triplicate samples were prepared per genotype.

Then, tissue samples were lysed in 50µl lysis buffer (5% SDS, 100mM TEAB, 10mM TCEP) by ultrasonication using a Bioruptor Pico ultrasonicator (Diagnode, Belgium) for 20 min at 4°C with subsequent heating to 95°C for 10 min. Proteins were alkylated using 15 mM iodoacetamide at 25°C in the dark for 30 min and protein concentration was determined using BCA colorimetric assay (Biorad). 20 ug of proteins were purified and digested by the SP3 approach (Hughes et al., 2019) using a Freedom Evo 100 liquid handling platform (Tecan Group Ltd., Männedorf, Switzerland). In brief, Speed BeadsTM (#45152105050250 and #65152105050250, GE Healthcare) were mixed 1:1, rinsed with water and diluted to the 8 µg/µL stock solution. Samples were adjusted to the final volume of 90 µL and 10 µL of the beads stock solution was added to them. Proteins were bound to the beads by addition of 100 µL of 100% acetonitrile to the samples, which were then incubated for 8 min at RT with a gentle agitation (200 rpm). Afterwards, samples were placed on a magnetic rack and incubated for 5 minutes. Supernatants were removed and discarded. The beads were washed twice with 160 µL of 70 % (v/v) ethanol and once with 160 of 100% acetonitrile. Samples were removed from the magnetic rack and 50 µL of digestion mix (10 ng/µL of trypsin in 50 mM triethylammonium bicarbonate) was added to them. Digestion was allowed to proceed for 12 hrs at 37°C. After digestion, samples were placed back on the magnetic rack and incubated for 5 minutes. Supernatants containing peptides were collected and dried under vacuum. Before LC-MS analysis, dried peptides were resuspended in 0.1% aqueous formic acid at a concentration of 0.2 ug/ul and a mix of heavy reference peptides (Supp. Table 4) was added to reach a final concentration of 10 fmol/ul/peptide.

Liquid chromatography-Mass spectrometric (LC-MS) analysis:

In a first step, parallel reaction-monitoring (PRM) assays (Peterson et al., 2012) were generated from a mixture containing 50 fmol of each proteotypic heavy reference peptide. Peptides were subjected to LC–MS/MS analysis using an Orbitrap Exploris

480 Mass Spectrometer fitted with an Vanquish Neo (both Thermo Fisher Scientific) and a custom-made column heater set to 60°C. Peptides were resolved using a RP-HPLC column (75μm × 30cm) packed in-house with C18 resin (ReproSil-Pur C18–AQ, 1.9 μm resin; Dr. Maisch GmbH) at a flow rate of 0.2 μLmin-1. Separation of peptides was achieved by using the following gradient: 4% Buffer B to 10% Buffer B in 3 min, 10% Buffer B to 35% Buffer B in 30min, 35% Buffer B to 50% Buffer in 7 min. Buffer A was 0.1% formic acid in water and buffer B was 80% acetonitrile, 0.1% formic acid in water. The mass spectrometer was operated in DDA mode with a total cycle time of approximately 1 s. Each MS1 scan was followed by high-collision-dissociation (HCD) of the 20 most abundant precursor ions with dynamic exclusion set to 5 seconds. For MS1, AGC was set to 300% in the Orbitrap over a maximum time of 50 ms and scanned at a resolution of 120,000 FWHM (at 200 m/z). MS2 scans were acquired at a target setting of 100%, maximum accumulation time of 50 ms and a resolution of 30,000 FWHM (at 200 m/z). Singly charged ions, ions with charge state ≥ 6 and ions with unassigned charge state were excluded from triggering MS2 events. The normalized collision energy was set to 30%, the mass isolation window was set to 1.4 m/z and one micro-scan was acquired for each spectrum. The acquired raw-files were searched using the MaxQuant software (Version 1.6.2.3) against a *Drosophila melongaster* database (consisting of 22088 protein sequences downloaded from Uniprot on 2022/02/22) using default parameters except protein, peptide and site FDR were set to 1 and Lys8 and Arg10 were added as variable modifications. The best 6 transitions for each peptide were selected automatically using an in-house software tool and imported into Skyline (Version 21.2). A scheduled (window width 4 min) mass isolation list containing the light and heavy peptide precursor masses was generated and imported into the MS operating software for PRM analysis. For PRM acquisition, total cycle time was set to not exceed approximately 4 s. For MS1, the following parameters were set: Resolution: 120,000 FWHM (at 200 m/z), Scan Range: 350-1600 m/z, Injection time: 25 ms, Normalized AGC Target: 300%. MS2 scans were acquired using the following parameters: Isolation Window: 0.4 m/z, HCD Collision Energy (normalized): 30, Normalized AGC target: 3000%, DataType: Centroid. For synthetic isotopically heavy labeled peptides, 22 ms injection time and 15,000 resolution (at 200 m/z) were set whereas for endogenous peptides 120,000 resolution and 250 ms injection time were set. All raw-files were imported into Skyline (Version 21.2) for protein / peptide quantification. To control for variation in injected sample amounts, the total ion chromatogram was used for normalization.

#### Results and discussion

Note that some results presented in this section are also mentioned in the main text. Below, emphasis is on the description of the proteomics data.

Four splice as well as three protein variants have been annotated for the *ap* locus (Supp. Fig. 6). Transcript ap-RA encodes the 469 aa ap-PA protein, ap-RC the 468 aa ap-PC and ap-RB and ap-RE the 246 aa protein (proteins ap-PB and ap-PE are identical). Note that protein variant specific reference peptides required for mass spectroscopy could only be designed for the Ap-469 and Ap-468. The 246 aa constituting Ap-246 are also part of the longer protein variants.

**1. Detection of annotated Ap proteins in tissue dissected from *y w* larvae**

In a first set of experiments, the presence of the two longer proteins was tested in samples containing central nervous system and imaginal disc tissue dissected from 3rd instar *y w* larvae. Only Ap-469 could be detected. Its N-terminal Methionine was found to be cleaved and acetylated (Supp. Fig. 8). Analysis with Ap-468 specific reference peptides indicated that this protein variant is not represented in our tissue samples (Supp. Fig. 7). However, it can't be excluded that it is produced in other tissues and/or developmental stages.

Based on the sequence of Ap-246, it completely lacks the first LIM domain. Of the second LIM domain, only the second Zn-finger is represented. The rest of the protein including the homeodomain is identical to Ap-469. Ap LIM domains are considered to be important in mediating protein–protein interactions (van Meyel et al 1999; Milan M, Cohen SM 1999; Rincon-Limas et al 2000). Thus, it seems conceivable that the ability of Ap-246 to interact with other LIM domain proteins is mutilated. Unfortunately, our proteomics data provides no direct evidence for the existence of the Ap-246. Its N-terminus was not found by SID-LC-MS having assays for all 4 possible variants. This included SID LC-MS analysis of the +/- methionine and +/- acetylation N-termini (analogous analysis as in Supp. Figs. 7 and 8). However, the low MS abundance of the reference peptides indicated that sensitivity of these assays was not very high and therefore not suited for firm conclusions (data not shown). Instead, using the absolute quantitative ability of our SID-LC-MS analysis, we calculated the relative concentrations of Ap-469 and Ap-246 from peptides shared between the two proteins and determined the ratio to Ap-469 protein alone using the Ap-469 specific peptides (Supp. Table 5). As a result, Ap-469 makes up 88% of the total Ap protein. This suggests that the remaining 12% must be contributed by Ap-246. More direct evidence for the existence of Ap-246 in $ap^{c2.34d}$ animals is presented below. At this point, it is unclear why Ap-246 is about 10 times less abundant than Ap-469 and what its role in development could be.

**2. Detection of Ap protein in various *ap* allele combinations**

The following measurements for a set of *ap* allele combinations were obtained with two reliable reference peptides. One is specific for Ap-469 (FYLSAVEK, 159-166), the other detects Ap-469 and Ap-246 (VLQVWFQNAR, 410-419). Several of the analyzed *ap* alleles do not produce any of the two proteins. They are $ap^{DG3}$, $ap^{DG8}$, $ap^{c1.72b}$ and $ap^{t11b}$ (Supp. Fig. 10). While this could be expected for $ap^{DG8}$, which deletes the entire *ap* ORF of both protein variants, the other three alleles could potentially produce one ($ap^{DG3}$) or two ($ap^{c1.72d}$, $ap^{t11b}$) transcripts that could be translated and produce Ap-246 (compare Figure 1 and Supp. Fig. 6). However, also this variant is not detected in these three alleles. A hypothesis to explain this observation posits that DNA at the 5' end of the ap-RA transcript is of relevance. $ap^{c1.72d}$ and $ap^{t11b}$ are two rather small deletions that only appear to affect expression of transcript ap-RA and hence the production of Ap-469. Absence of Ap-246 could therefore imply that regulatory elements deleted by these two deletions could also affect ap-RB transcription. However, deletions $ap^{c1.72d}$

and *ap*$^{t11b}$ are located far from the ap-RB transcription start site. Their deletion break points suggest that a 526 bp fragment at the 5' end of transcript ap-RA could also be responsible for the production of transcript ap-RB. Remarkably, the 526 bp fragment contains the single Polycomb Response Element which has been mapped to the *ap* locus (Schwartz et al 2006; Oktaba et al 2008). It remains to be shown how the apPRE influences the promoter of the ap-RB transcript at a distance of ~17'500 bp.

In general, it is assumed that animals with two gene copies produce double the amount of protein than animals with only one gene copy. Surprisingly, heterozygous *ap*$^{DG8}$/+ larval tissue produces similar amounts of Ap as *y w* controls (Supp. Figure 9). A linear correlation between gene dosage and protein concentration is also not observed in tissue samples dissected from *ap*$^{cDNAint2.3}$ larvae. While homozygotes produce similar amounts of Ap as *y w* controls, heterozygous (over *ap*$^{DG8}$) larval tissue still contain about 80% instead of only 50% compared to the *y w* control. A possible explanation for this discrepancy is that in these animals, the remaining gene copy is still controlled by two regulatory regions known to contain imaginal disc and brain specific enhancers, not just one as in true hemizygotes.

##### 3. Detailed analysis of *ap*$^{c2.34d}$ reveals presence of truncated Ap-469 protein

Allele *ap*$^{c2.34d}$ was analyzed in more detail because homozygotes are viable, fertile and their ampulla contains 4 papillae (Figure 3K). However, they only develop small winglets (Figure 3C). These observations suggested that *ap*$^{c2.34d}$ flies produce a protein that still can promote ampulla but not wing development. The associated deletion removes the translation start site of Ap-469 but not the corresponding transcription start site (Figure 1D). Nevertheless, this allele still produces a truncated version of this protein (Supp. Fig. 10). In agreement with the deletion break points, measurements with Ap-469 specific reference peptide FLGPGAR (19-25) failed to detect Ap-469. In contrast, this protein could be detected with reference peptides NDYYSFFGTR (199-208) and CLASISSNELVMR (213-225). This suggests that translation of a truncated protein is reinitiated from a codon between reference peptides FLGPGAR (19-25) and NDYYSFFGTR (199-208). Previous measurements with reference peptide FYLSAVEK (159-166) also detected a truncated Ap-469 (Supp. Figure 9), thus further reducing the interval for reinitiation by another 40 aa (positions 25 to 159 of the Ap-469 ORF). Determination of the putative translation start site was not successful (data not shown). But compared to *y w* larval tissue, truncated Ap-469 is detected at clearly lower concentrations with Ap-246 specific reference peptides NDYYSFFGTR (199-208) and CLASISSNELVMR (213-225). In contrast, similar Ap levels were measured with reference peptides SYFAINHNPDAK (384-395) and VLQVWFQNAR (410-419) (Supp. Fig. 10). A possible explanation for this difference is that Ap-246 is expressed in *ap*$^{c2.34d}$ animals. Note that the apPRE is present in this allele while it is gone in *ap*$^{c1.72d}$.
Measurements made with only two instead of five reference peptides suggest that allele *ap*$^{77f}$ might produce similar protein variants as *ap*$^{c2.34d}$ (Supp. Fig. 9). In *ap*$^{77f}$, a non-sense mutation at the 15$^{th}$ codon of the Ap-469 ORF was detected, in theory leading to an apparent null-allele (Table 1). Our data suggests that in hemizygous

*ap*$^{77f}$/*ap*$^{DG8}$ animals, translation might also be reinitiated from an alternate start codon as in *ap*$^{c2.34d}$//*ap*$^{DG8}$ animals. Genetically, *ap*$^{c2.34d}$ as well as *ap*$^{77f}$ are classified as hypomorphic, which implies that these truncated Ap-469 proteins can still exert part of their normal function.

**4. Detection of Ap-246 protein in *ap*$^{ex5\text{-}8}$ larvae**

In an attempt to elucidate the function of Ap-246, flies producing only this version of Ap were generated. The allele is called *ap*$^{ex5\text{-}8}$ (Figure 1A). In such flies, the corresponding transcript is produced under the control of the transcription start site normally leading to Ap-469 (ap-PA in Supp. Figure 6). Hemizygous *ap*$^{ex5\text{-}8}$ flies have no wings and die precociously. This suggests that Ap-246 has no role in wing and ampulla development. A trivial explanation for this observation could be that the gene product is not translated. This possibility was ruled out by protein spectroscopy and anti-ap immune detection in embryos (Supp. Figure 11). As above, tissue samples were dissected from late 3$^{rd}$ instar larvae and protein extracts were analyzed with two reference peptides, one specific for Ap-469, the other detecting both protein variants. Ap protein in *ap*$^{ex5\text{-}8}$ tissue was only detected with the latter. In *ap*$^{cDNAint2.3}$ control tissue samples, Ap was detected with both reference peptides. But unexpectedly, *ap*$^{ex5\text{-}8}$ flies produce about 6 times more Ap-246 than Ap-469 is detected in *ap*$^{cDNAint2.3}$ (Supp. Figure 11A). Whether this difference contributes to the apparent phenotypic differences, is not clear. At least in embryos, expression patterns obtained with an Ap homeodomain specific antibody are very similar in hemizygous *ap*$^{cDNAint2.3}$ and *ap*$^{ex5\text{-}8}$ embryos (Supp. Figure 11B, C).

#### Figures

**Supp. Fig. 6:** 4 transcripts and 3 proteins annotated for the *apterous* locus

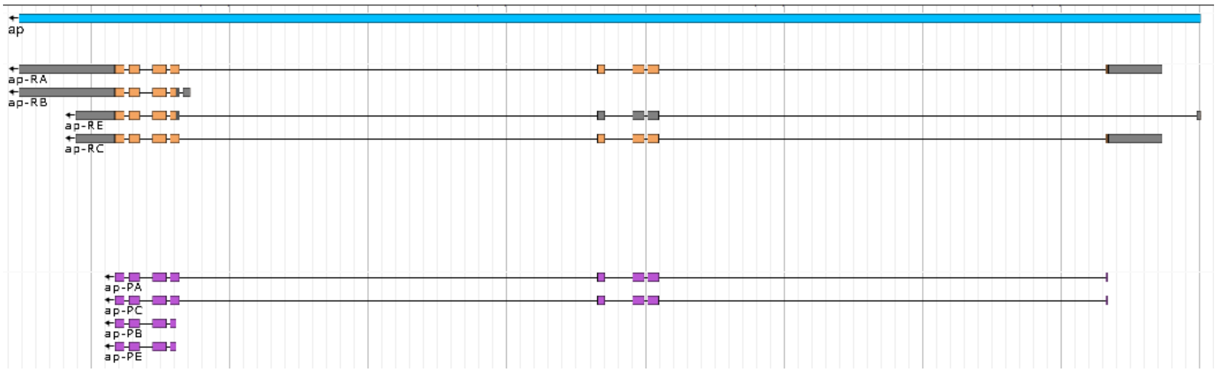

**Supp. Fig. 7:** Targeted SID LC-MS analysis of N-terminal peptides of Ap-468 in *y w* larval tissue.

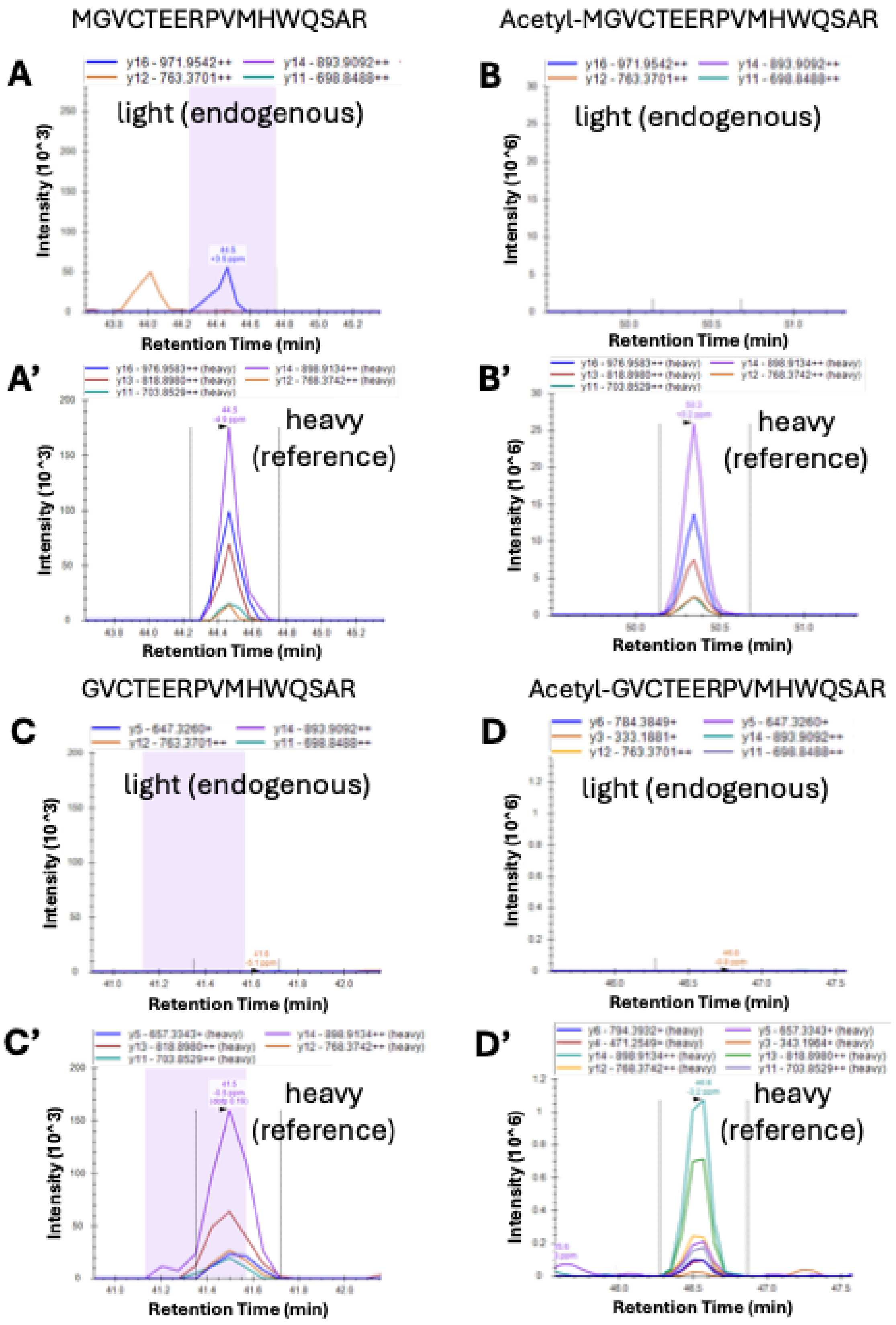

**Supp. Fig. 8:** targeted SID LC-MS analysis of N-terminal peptides of Ap-469 in *y w* larval tissue.

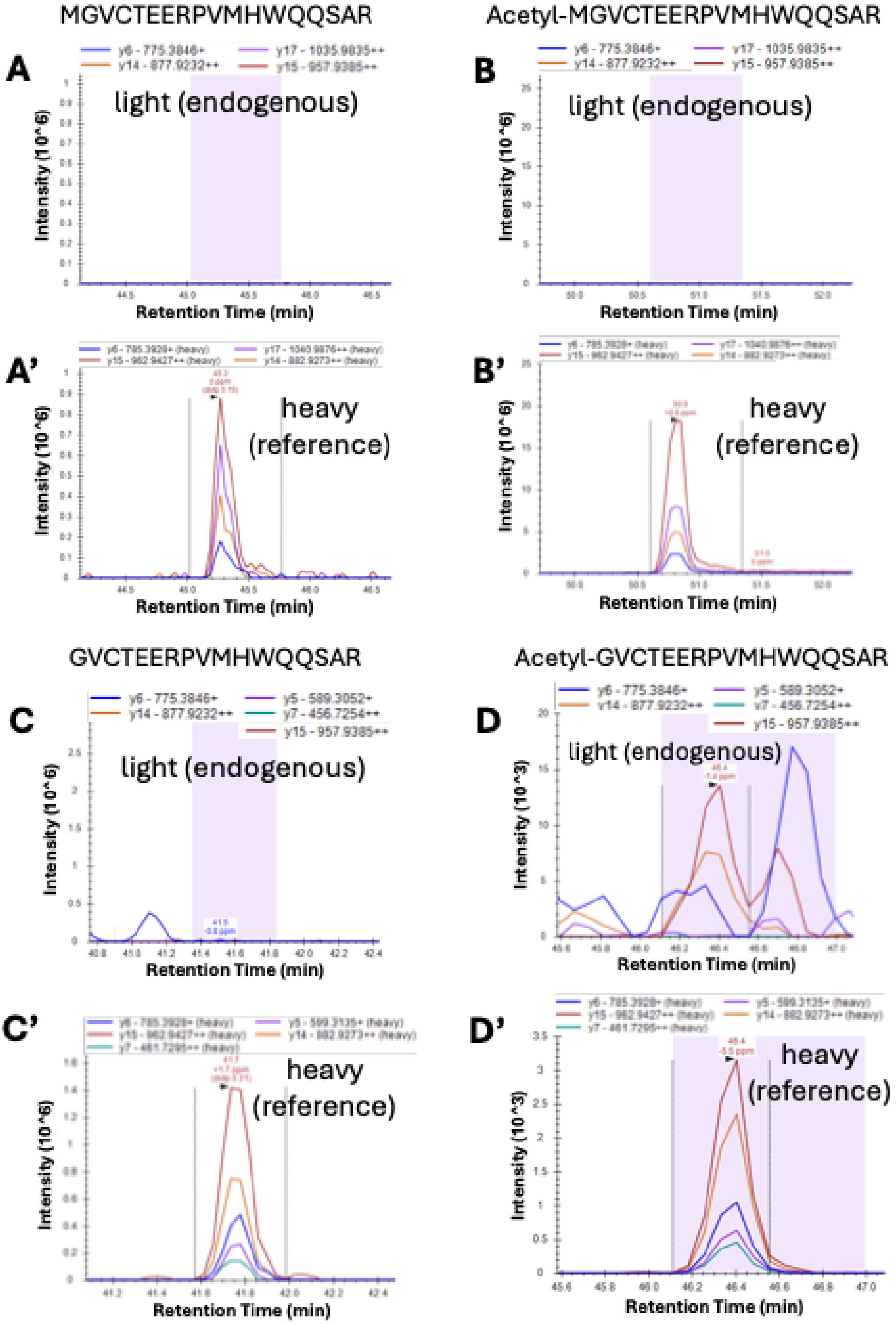

**Supp. Fig. 9:** analysis of Ap-469 and Ap-246 in several *ap* allele combinations

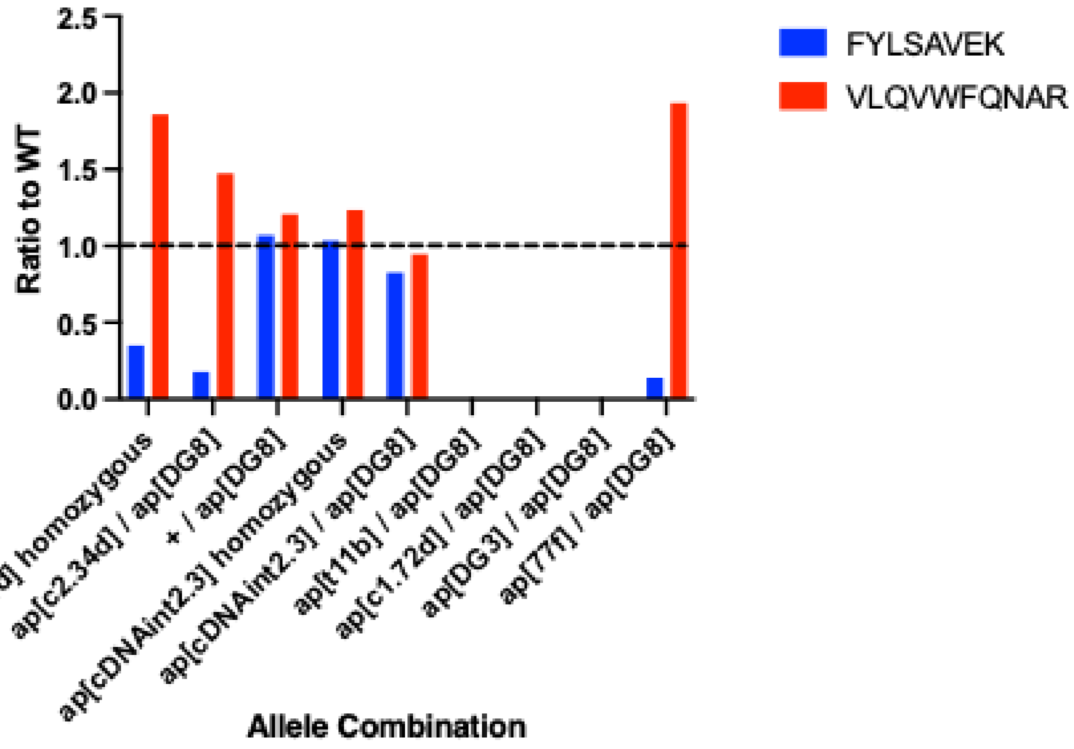

**Supp. Fig. 10:** analysis of *ap*$^{c2.34d}$ reveals usage of alternative translation start site

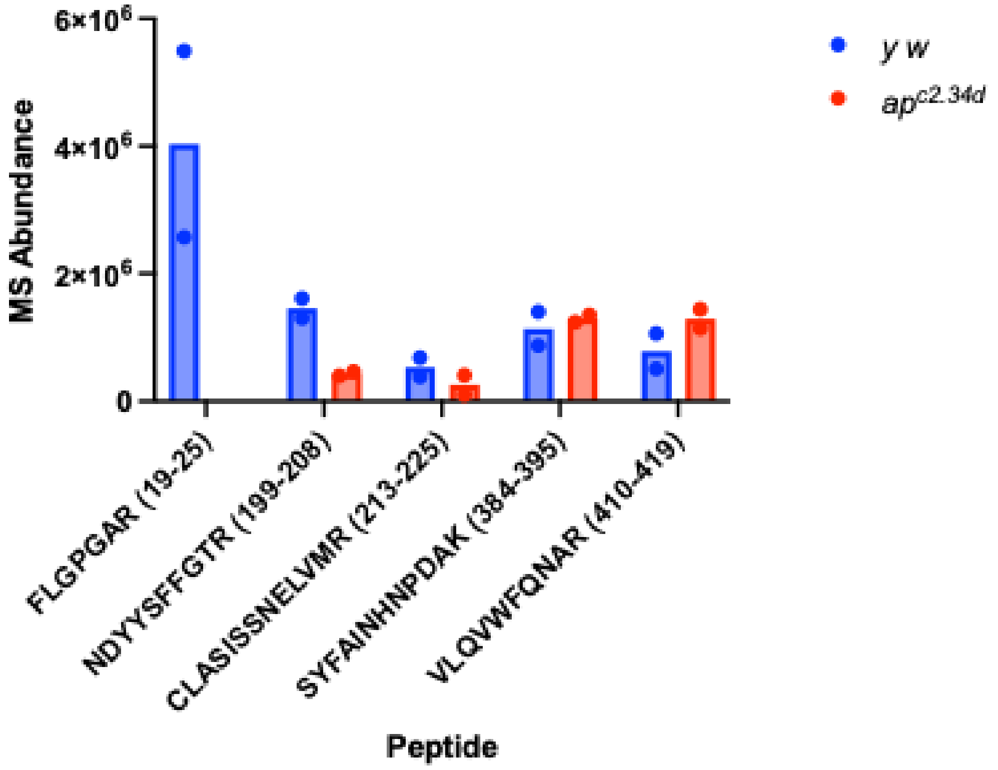

**Supp. Fig. 11:** Identification of Ap-469 in $ap^{cDNAin2.3}/ap^{DG3}$ and Ap-246 in $ap^{ex5\text{-}8}/ap^{DG3}$ larval tissue

(A)

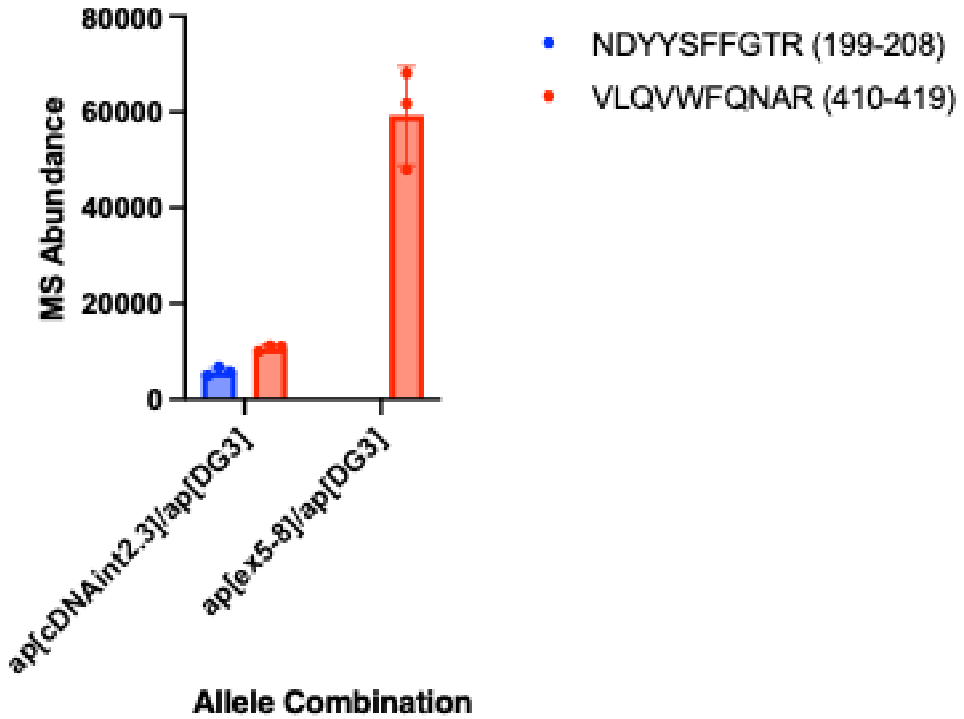


(B)

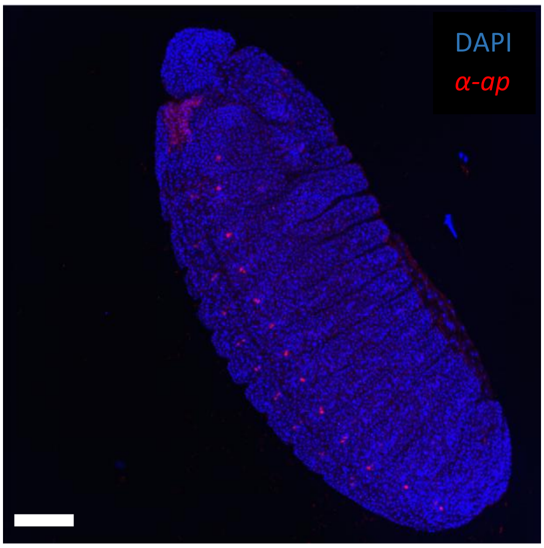


$ap^{cDNAint2.3}/ap^{DG3}$

(C)

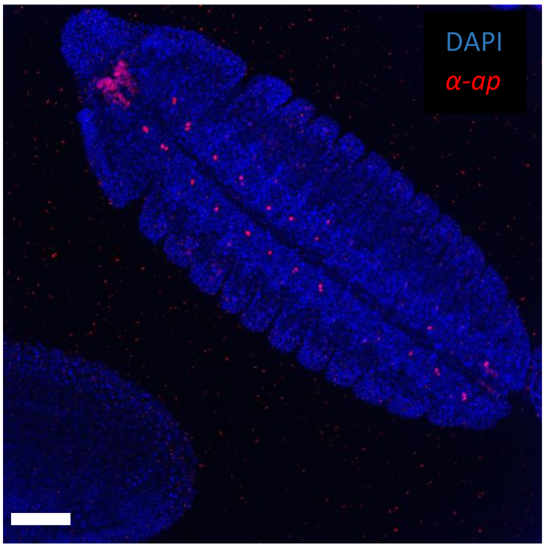


$ap^{ex5\text{-}8}/ap^{DG3}$

#### Figure Legends

**Supp. Fig. 6:** Annotated *apterous* transcripts and proteins are shown.
At the top of the panel, the blue bar depicts the 21.3 kb gene span region extending between the most 5' transcription start site and the most 3' UTR. Below, the 4 transcripts ap-RA, ap-RB, ap-RC and ap-RE are shown in grey and orange. The latter indicates the coding parts in each transcript. At the bottom, the 3 putative translated protein products are shown in magenta. ap-PA corresponds to Ap-469, ap-PB to Ap-468 and ap-PC and ap-PE to Ap-246. FlyBase (release 6.64) was used to find information on annotated mRNAs and protein variants (Jenkins et al., 2022).

**Supp. Fig. 7:** Targeted SID LC-MS analysis of N-terminal peptides of Ap-468 in *y w* larval tissue.
Four isotopically marked (heavy) reference peptides were used for the identification and quantification of the N-terminal end of Ap-468. Their sequence is indicated at the top of each panel. They differ in their N-terminal ends: (A,A') unmodified Methionine; (B,B') acetylated Methionine; (C,C') unmodified Glycine (Methionine cleaved); (D,D') acetylated Glycine. A-D: transitions obtained with proteolytically cleaved tissue samples ("light (endogenous)"). A'-D': transitions obtained with the spiked heavy reference peptides that have been added to the samples ("heavy (reference)"). These serve as an internal reference for the corresponding light peptide generated by proteolysis of the endogenous target protein shown in A-D. The Skyline software tool was used for peptide quantification and identification. At least 2 transitions (fragment traces) represented in A'-D' were required per peptide for its identification in A-D, respectively. Using this threshold, none of the N-terminal peptides of Ap-468 were identified by targeted SID-LC-MS considering +/- methionine and +/- acetylation. It can thus be concluded that Ap-468 is not represented in the tissue samples.

**Supp. Fig. 8:** Targeted SID LC-MS analysis of the N-terminal peptides of Ap-469 in *y w* larval tissue.
Targeted LC-MS data obtained with Ap-469 specific reference peptides depicted analogously as in Supp. Fig. 7. Only the N-terminal methionine cleaved and N-terminal acetylated proteoform of Ap-469 was identified requiring at least 2 transitions per peptide. Hence, it can be concluded that Ap-469 is contained in the tissue samples.

**Supp. Fig. 9:** Ap concentration in various *ap* allele combinations relative to *y w*.
Measurements obtained with reference peptides FYLSAVEK (159-166; Ap-469 specific; ratio to wild-type (*y w*) indicated by blue bars) and VLQVWFQNAR; 410-419; detects Ap-469 and Ap-246; ratio to wild-type (*y w*) indicated by red bars) are shown. $ap^{DG3}$, $ap^{DG8}$ $ap^{c1.72d}$ and $ap^{t11b}$ are lacking both protein variants and thus can be considered as true $ap^{null}$ alleles. For alleles $ap^{c2.34d}$ and $ap^{cDNAint2.3}$, homozygous and hemizygous conditions were assayed. Note that hemizygosity only extends over the entire ORF and the apPRE but not the intergenic spacer between *ap* and *l(2)09851* (see Figure 1). Hence two copies of all enhancers located in the intergenic spacer are influencing ap transcription in $ap^{c2.34d}/ap^{DG8}$ and $ap^{cDNAint2.3}/ap^{DG8}$, not only one. This

could account for the observation that such hemizygous tissue produces more than the expected 50% of Ap protein. Allele $ap^{77f}/ap^{DG8}$ tissue yielded a similar ratio to wild-type as $ap^{c2.34d}/ap^{DG8}$ did. This suggests that $ap^{77f}$ might also produce a truncated Ap-469 protein because the ratio is much lower for reference peptide FYLSAVEK (159-166) than for VLQVWFQNAR; 410-419).

**Supp. Fig.10:** Evidence for a truncated Ap-469 protein in $ap^{c2.34d}$ tissue.
Relative amounts of Ap (Mass Spectrometric (MS) abundance in vertical axis) detected in *y w* (blue bars) and $ap^{c2.34d}$ (red bars) tissue for five reference peptides located along the *ap* ORF are shown. Sequence and position of the reference peptides is indicated along the horizontal axis. The first three peptides on the left are Ap-469 specific, the last two detect Ap-469 and Ap-246. Note that aa positions 19 to 25 of Ap-469 are not detected in $ap^{c2.34d}$ tissue samples. The next two Ap-469 specific peptides are detected at clearly reduced levels compared to *y w*. Finally, similar Ap amounts are detected with the two most C-terminal reference peptides which detect Ap-469 and Ap-246. The increase in Ap levels between the 3$^{rd}$ and the 4$^{th}$ reference peptide could suggest that Ap-246 is expressed in $ap^{c2.34d}$ tissue and thus contributes to the total Ap expression. The individual data points and the mean represented by the bar are indicated.

**Supp. Fig. 11:** Detection of Ap-469 in $ap^{cDNAint2.3}/ap^{DG3}$ and Ap-246 in $ap^{ex5\text{-}8}/ap^{DG3}$ larval tissue
**(A)** MS abundance of Ap-469 in $ap^{cDNAint2.3}/ap^{DG3}$ and Ap-246 in $ap^{ex5\text{-}8}/ap^{DG3}$ larval tissue are shown. Reference peptide NDYYSFFGTR (199-208) is specific for Ap-469 and VLQVWFQNAR (410-419) is specific for Ap-469 and Ap-246. In $ap^{cDNAint2.3}/ap^{DG3}$ tissue samples, both peptides were identified by LC-MS. For the protein produced by $ap^{ex5\text{-}8}/ap^{DG3}$ animals, only VLQVWFQNAR was identified. This strongly suggests that allele $ap^{ex5\text{-}8}$ only produces Ap-246. The individual data points, mean (height of the bar) and SD are indicated.
**(B, C)** Immune detection of Ap-469 in $ap^{cDNAint2.3}/ap^{DG3}$ (B) and Ap-246 in $ap^{ex5\text{-}8}/ap^{DG3}$ (C) embryos shown with anterior to the left. Note that Ap positive cells are detected in a very similar pattern in both genotypes. Scale bar: 50µm.

#### Tables

**Supp. Table 4:** Heavy reference peptides used for targeted LC-MS*

| Peptide | Length | Specificity | Position |
|---|---|---|---|
| MGVCTEERPVMHWQQSAR | 18 | 469 | 1 - 18 |
| Ac-MGVCTEERPVMHWQQSAR | 18 | 469 | 1 - 18 |
| GVCTEERPVMHWQQSAR | 17 | 469 | 2 - 18 |
| Ac-GVCTEERPVMHWQQSAR | 17 | 469 | 2 - 18 |
| MGVCTEERPVMHWQSAR | 17 | 468 | 1 – 17 |
| Ac-MGVCTEERPVMHWQSAR | 17 | 468 | 1 – 17 |
| GVCTEERPVMHWQSAR | 16 | 468 | 2 – 17 |
| Ac-GVCTEERPVMHWQSAR | 16 | 468 | 2 – 17 |
| FLGPGAR | 7 | 468+469 | 19 – 25 |
| FYLSAVEK | 8 | 468+469 | 159 - 166 |
| NDYYSFFGTR | 10 | 468+469 | 199 – 208 |
| CLASISSNELVMR | 13 | 468+469 | 213 – 225 |
| GDQYGIIDALIYCR | 14 | all | 248 – 261 |
| SYFAINHNPDAK | 12 | all | 384 – 395 |
| VLQVWFQNAR | 10 | all | 410 - 419 |

* Sequences of 15 reference peptides are shown in order along the Ap protein sequence. Their length, specificity and position are also indicated. The N-terminal Met (M) of Ap-469 is labeled in blue, the one of Ap-246 in red.

**Supp. Table 5**: Estimating the relative amounts of Ap-469 and Ap-246n in *y w* larval tissue*

| Peptide | Ratio to reference | Average | Ratio 469aa/both |
|---|---|---|---|
| FLGPGAR (469aa) | 0.0133 | 0.0129 | 0.8843 |
| NDYYSFFGTR (469 aa) | 0.0126 | | |
| GDQYGIIDALIYCR (both) | 0.0145 | 0.0146 | |
| VLQVWFQNAR (both) | 0.0147 | | |

* Using the absolute quantitative ability of our SID-LC-MS analysis, we calculated the relative concentrations of Ap-469 and Ap-246 from peptides shared between the two proteins and determined the ratio to Ap-469 protein alone using the Ap-469 specific peptides. The ratio of 0.88 indicated that the long version made up 88% of the Ap molecules whereas the short version contributed 12%. This also shows that the short version is expressed in wild type samples.
